# Antimicrobial resistant bacteria in wastewater-irrigated Mexican soils and transfer of resistant bacteria from irrigated soils to cilantro plants

**DOI:** 10.64898/2026.05.17.725719

**Authors:** Dipen Pulami, Deeksha Bhati, Sara Gallego, Kornelia Smalla, Kathia Lüneberg, Christina Siebe, Benjamin Heyde, Jan Siemens, Stefanie P. Glaeser

**Author notes:** Correspondence: Stefanie P. Glaeser.

## Abstract

Agricultural fields in the Mezquital Valley, Mexico, were irrigated with untreated wastewater over several decades. Following the construction of a wastewater treatment plant (WWTP) in Atotonilco de Tula, WWTP effluent is used for irrigation. To evaluate the effects of changed irrigation, a soil incubation experiment was performed. Soils of the Mezquital Valley long-term irrigated with untreated wastewater were irrigated with WWTP influent or effluent, both unspiked and spiked with antibiotics and biocidal compounds and incubated four weeks. We investigated the effects of shifted irrigation on the abundance of cultivable total heterotrophic and resistant bacteria (RB). Additionally, RB were cultivated from *Coriandrum sativum* (cilantro) sown in soil of the incubation experiment. While wastewater treatment significantly reduced the bacterial abundance in effluent, spiking increased RB abundance in both wastewater types including ciprofloxacin (CIP) RB. Before wastewater addition, all soils contained cultivable RB. Irrigation increased the relative abundance of RB cultivated on Mueller Hinton (MH) agar in Leptosols and Phaeozems, compared to soils prior to wastewater addition irrespective of the water type, but not in Vertisols, suggesting the soil type rather than water qualities influenced the RB abundance. Diverse CIP RB were cultivated including strains of 14 genera of three phyla. Among those, *Achromobacter* spp. strains related to potentially pathogenic *A. spanius* originating from soil were abundant in both leaves and roots of cilantro. Our results showed that the implementation of wastewater treatment does not reduce the abundance of cultivable RB in Mezquital Valley soils and cilantro plants. Health risk associated monitoring should include long-term persistent RB colonizing plants cultivated in wastewater irrigated soils.

## Introduction

Irrigation with treated or reclaimed wastewater from wastewater treatment plants (WWTPs) is practiced increasingly to compensate for freshwater shortage, droughts, and meeting growing global food demands.^1–4^ A WWTP is considered as an interface between the sewage and the receiving environment. Wastewater treatment processes reduce but do not efficiently remove pharmaceuticals, pesticides, bacteria carrying antibiotic resistance genes (ARGs), and antimicrobial resistant bacteria (ARB) from wastewater.^5–8^ Therefore, not only untreated wastewater, but also treated wastewater (WWTP effluent) contains pharmaceuticals and disinfectants, heavy metals, ARGs, and ARB that can contribute to the exchange and spread of antimicrobial resistance among bacteria in the environment.^9–13^ Long-term irrigation with wastewater leads to the accumulation of chemical pollutants in soils, which can be taken up by plants.^14–16^ Moreover, plants can be colonized by resistant environmental and/or facultative pathogenic bacteria.^17–19^ In consequence humans might get exposed to ARB and/or facultative pathogenic bacteria via consumption of fresh produce which was irrigated with wastewater or grown in wastewater-irrigated soils. In this study, we used the term resistant bacteria (RB) to designate bacteria resistant to antibiotics (ARB) and tolerant to biocidal compounds with the focus on quaternary alkylammonium compounds (QAACs).

Owing to their chemical stability, several antibiotics, especially fluoroquinolones, sulfonamides, and tetracyclines, are known to degrade slowly, resulting in their persistence and accumulation in the environment.^20–26^ A strong accumulation of antibiotics (ciprofloxacin, CIP and sulfamethoxazole, SUL), biocidal compounds assigned to the class of QAACs, and heavy metals was found in soils of agricultural fields of the Mezquital Valley. Those soils were irrigated with untreated wastewater for more than a century.^27–31^ The Mezquital Valley is a major agricultural area located in the federal state of Hidalgo in Mexico. Dalkmann *et al.* showed an enrichment of enterococci and ARGs in long-term irrigated soils in the Mezquital Valley.^28^ Broszat *et al*. showed that culturable *Gammaproteobacteria*, including facultative pathogens, such as bacteria of the genera *Pseudomonas*, *Stenotrophomonas*, and *Acinetobacter,* and several resistance genes accumulated in the long-term irrigated soils.^32^ The fraction of cultivable RB was increased in wastewater-irrigated compared to rain-fed soils and multi-resistant *Stenotrophomonas* strains were selectively cultivated from irrigated soils.^32^

Most studies considering wastewater use in agriculture hitherto have extensively focused on the effects of irrigation with WWTP effluent upon replacing fresh water.^1,16,33,34^ Other studies have focused on effects of treated wastewater (WWTP effluent) on soil properties and bacterial community.^35–41^ The transition from long-term irrigation with untreated wastewater to irrigation with WWTP effluent has rarely been investigated.^42^ Changing irrigation with untreated wastewater to WWTP effluent could open a window of transiently elevated risk of the release of pollutants such as antibiotics, biocidal compounds, and heavy metals which were accumulated over time in irrigated soils. This may foster the proliferation of antimicrobial resistance.^42–44^

Since the construction of a large WWTP in Atotonilco de Tula at the entrance of the Mezquital Valley, WWTP effluent has been integrated into the irrigation regime of fields in the Mezquital Valley.^45–47^ This makes the Mezquital Valley a unique environment were consequences associated with changes in the irrigation with untreated wastewater to WWTP effluent get relevant. Ziegler Rivera *et al*.^48^ and Heyde *et al.*^49^ showed that this irrigation shift can increase the mobility of potential toxic elements including heavy metals in soils of the Mezquital Valley.

Up to date only the study of Soufi *et al.* investigated the effects of the transition from irrigation with untreated wastewater to irrigation with WWTP effluent on the soil microbiome and ARG abundance in more detail.^50^ A soil incubation experiment was performed with soil samples of different soil types (Leptosol, Phaeozem, Vertisol) from the Mezquital Valley. The soils have been irrigated with untreated wastewater for at least 60 years. Soil samples were irrigated either with untreated wastewater (WWTP influent, representing ongoing irrigation with untreated wastewater) or the WWTP effluent (mimicking transition to irrigation with treated wastewater) from Atotonilco WWTP. In addition, both irrigation water types were spiked (spiked-influent and spiked-effluent) with an additional level of antibiotics and QAACs (mimicking enhanced pollutant exposure).^50^ Soufi *et al.* used cultivation independent DNA-based approaches to study the relative abundance of ARGs and investigated the bacterial communities present in the water types and the irrigated soils. It was shown that ARG abundances were affected by the spiking. Contrarily the composition of the soil bacterial communities was neither affected by the irrigation water types nor the spiking of irrigation water with increased concentrations of antibiotics and biocidal compounds but by the soil type.^50^ Heyde *et al.*^49^ reported strong soil type effects on soluble concentrations of nutrients, metals, and carbon in the soils from the same soil incubation experiment described by Soufi *et al*.^50^ We aimed to complement the study of Soufi *et al.* with a cultivation-dependent approach by the investigation of cultivable RB present in the soils from the same soil incubation experiment.^50^ We cultivated RB from all wastewater types and soils before (0 days) and 4 weeks incubation post water addition.^49,50^ We targeted heterotrophic bacteria growing under higher nutrient conditions at 37°C (among them facultative pathogenic bacteria) and heterotrophic bacteria growing under lower nutrient conditions at 25°C (environmental adapted bacteria) , respectively. In addition, we aimed to cultivate faecal indicator bacteria including *Escherichia coli* and enterococci and their resistant subgroups.

In addition, we considered the aspect that the changes in the irrigation regime may also shift the use of agricultural fields in the Mezquital Valley. Traditionally wastewater-irrigated fields in the Mezquital Valley have been mainly used for cultivation of Alfalfa and maize as livestock fed and food. However, an extended use e.g. for the cultivation of cilantro (*Coriandrum sativum L.*), a common fresh produce cultivated in Mexico, may be possible after the implementation of wastewater treatment.^32,51^ For that reason, we used the irrigated soils (incubated for 14 days post irrigation) from the incubation experiment to study the transfer of RB from soil to cilantro (*Coriandrum sativum L.*) plants in a subsequently performed greenhouse experiment.

Our study tested the following hypotheses: (I) The total abundance of RB increases after changing irrigation from untreated wastewater to irrigation with effluent in Mexican soils with a history of long-term irrigation with untreated wastewater, (II) spiking of irrigation water with antibiotics and QAACs increases the RB abundance, (III) soil types modulate the abundance of RB, and (IV) differently irrigated soils all carry RB which can be transferred to fresh produce, such as *Coriandrum sativum* (cilantro), a herb commonly grown in the Mezquital Valley and consumed raw.

## Materials and Methods

### Wastewater and soil sampling

The Atotonilco WWTP has a treatment capacity of 35 m^3^ per second and a maximal capacity of 50 m^3^ per second (World Bank, 2018).^46^ The WWTP started operation at 30% of its capacity in 2017 (10.5 m^3^ per second) and continued at full capacity from 2020 onward. The main aim of the wastewater treatment is to provide high-quality water (treated wastewater or WWTP effluent) for farmland irrigation, while preserving dissolved nutrients.^45–47^ Composite wastewater samples were collected in April 2022. The Atotonilco WWTP (19°57’28.52"N, 99°17’51.02"O) influent or effluent transported via the Salto-Tlamaco channel (sampled circa 1 km downstream from the WWTP and upstream of any settlements) were collected by mixing two individual samples of 10 L each taken daily over three consecutive days (i.e., a total 60 L of each wastewater type). Samples were stored at 4°C and transported cooled via airfreight to Germany for the experiment.

Soil samples were taken from four different locations in the Mezquital Valley area: “Ulapa de Melchor Ocampo”, “Tetepango”, “Tlaxcoapan”, and “Tlahuelilpan” (16-23 km Northeast from Atotonilco WWTP) (Figure S1). Soils were taken from fields planted with Alfalfa (*Medicago sativa*) representing three typical soil types found in the region: Leptosol, Phaeozem, and Vertisol, respectively. Untreated wastewater had been used for irrigation of those fields for more than 60 years in the past. Soil samples were taken from a transect 5 to 10 m parallel to the water inlet of the fields. Four composite samples per field site were taken by mixing eight individual samples from four distinct locations from 0-30 cm depth. Four different fields per soil type were sampled. Soil samples were stored at ambient temperature and transported under cooled conditions (<8°C) to Germany. Further details to the soil and wastewater sampling procedures and physicochemical characteristics are provided by Soufi *et al*. and Heyde *et al.*^49,50^

### Soil Incubation experiment (soil microcosms)

The experimental setup is explained in detail by Soufi *et al.*^50^ An overview of the setup is additionally provided in Figure S2. The four composite samples per field were sieved to 8 mm size, homogenized, and mixed to one composite sample per field. The obtained composite soil samples were used as soil type replicates. Composite samples were obtained from four fields per soil type. Individual soil microcosms were filled with 200 g (dry weight equivalent) of soil. For each sampling time point, one microcosm per replicate was prepared. To each of the soil microcosms 25 mL of one of the four water type was added: (i) unspiked-influent, (ii) unspiked-effluent, (iii) spiked-influent, or (iv) spiked-effluent (Figure S2). Spiking of water was done by adding six antibiotics namely azithromycin (AZI), CIP, clindamycin (CLIN), erythromycin (ERY), SUL, trimethoprim (TRIM), and a mixture of different QAACs including benzyl alkyldimethyl ammonium chloride (BACs), dialkyldimethyl ammonium chloride (DADMACs), and alkyltrimethyl ammonium chloride (ATMACs) Spiking raised the antibiotic and QAAC levels by 500-fold compared to those measured in the influent (for details see Table S1). The water content of the soils was then adjusted to 80% water holding capacity by adding deionized water, considering the initial water content of the field-moist samples. The soil microcosms were covered with sterile surgical masks, placed in a climatic chamber at 95% humidity and 22°C temperature, and incubated for a period of up to 8 weeks. For more details see Soufi *et al.* and Heyde *et al.*^49,50^ In this study, we investigated the cultivable bacterial abundance in soil samples before water addition (0 days, non-incubated, field composite samples), i the four water types used for irrigation, and in soil microcosms four weeks after addition of the four types of irrigation water (Figure S2).

### Enumeration of total and antibiotic and QAAC tolerant heterotrophic bacteria

We applied two cultivation strategies to determine the abundance of total and antibiotic and QAAC tolerant heterotrophic bacteria. We aimed to differentiate between fast-growing, copiotrophic bacteria, which grow well under high nutrient conditions on Mueller Hinton agar (MH; Carl Roth GmbH) at 37°C within 24 h, and those which grow well at lower nutrient conditions on Reasoner’s 2A agar (R2A; Carl Roth GmbH) at 25°C within 4 days. To keep it simple we used the terms "MH and R2A grown bacteria" throughout this study. MH agar is a non-selective and non-differential nutrient-rich medium and commonly used for antibiotic susceptibility testing (AST), isolation, and enumeration of fast-growing heterotrophic bacteria, including pathogens associated with animals and humans.^52,53^ R2A agar contains lower nutrient concentrations than MH and is commonly used to obtain also slower growing oligotrophic environmental bacteria.^54,55^ In addition, we applied cultivation conditions for the selective cultivation of faecal indicator bacteria (*E. coli,* enterococci) and resistant subgroups targeting 3rd generation cephalosporin-resistant (3GCR) *E. coli* and vancomycin-resistant enterococci (VRE).

Briefly, from each field composite sample and each soil microcosm 1 g of soil was suspended in 9 mL tetra sodium pyrophosphate buffer (TSPP, 0.2% filter sterilized using 0.45 µm sterile 50 mL syringe filter, Whatman) in a sterile 15 mL falcon tube (Greiner Bio-one) to detach bacteria from the soil matrices. After incubation on a horizontal shaker (150 rpm, 10 min) in the dark, soil suspensions were poured inside stomacher bags (Seward circulator, 400) and mechanically treated in a stomacher (Seward Bio-master Lab System) at maximum speed for 3 min. The suspensions were transferred again to the 15 mL tubes and left in the dark for 10 to 15 min to allow soil particles to settle down by sedimentation. Further analysis was performed with 2 mL of the supernatants containing bacterial cells detached from the soil matrix. Next, 30 µL of each supernatant and each irrigation water type were serially diluted (ten-fold up to 10^-6^) in sterile 96 wells plates filled with 270 µL sterile (autoclaved) sodium chloride solution (NaCl, 0.9% w/v). Dilution series of each individual sample were performed in quadruplets (technical replications). Subsequently, 5 µL of each dilution were spotted on MH and R2A plates without and with antibiotics and biocidal compounds: CIP (1 and 4 µg mL^-1^, Sigma Aldrich), TRIM/ SUL (4/76 µg mL^-1^, Sigma Aldrich), ERY (2 µg mL^-1^, Cayman Chemical Company)/ CLIN (0.5 µg mL^-1^, Sigma Aldrich), and BAC-C12 (50 µg mL^-1^, Tokyo Chemical Company).

For the cultivation of faecal indicator bacteria the dilution series were spotted on selective differential media including Tryptone Bile X-Glucuronide agar (TBX, Carl Roth GmbH, Germany) without and with cefotaxime (CTX, 1 µg mL^-1^, Sigma Aldrich) and Slanetz and Bartley (SB) agar (Carl Roth GmbH) without and with vancomycin (VAN, 4 µg mL^-1^, Sigma Aldrich) to cultivate *Escherichia coli*, 3GCR *E. coli*, enterococci, and VRE, respectively. Plates for total and 3GCR *E. coli* were incubated under oxic conditions for 3 h at 37°C and moved to 44°C for 21 h. Plates for total enterococci and VRE were incubated under microaerophilic conditions using Anaerocult C gas Packs (Sigma-Aldrich) at 37°C for 48 h. All agar media were supplemented with cycloheximide (100 µg mL^-1^, Biochemica) to inhibit fungal growth. Following incubation, colonies were counted manually to calculate colony forming units (CFUs) per mL or g analysed sample. Counting was performed after 24 and 48 h for MH and R2A plates, respectively. Longer incubation times (48 h for MH, and more than 4 days for R2A) hindered an efficient colony counting. As 5 µL were spotted per dilution, 3 to 30 CFUs were assumed as a countable range. Additionally, most probable numbers (MPNs) of bacteria per mL or g were calculated based on the number of spots per dilution that showed growth using the method of Jarvis et al.^56^ Because the concentrations determined by CFU and MPN counts showed homogenous trends throughout the samples, only CFU counts are presented in this study. Stored wastewater samples (4°C) were studied here in parallel to soil samples after the soil incubation experiment was started.

### Cultivation and sampling of cilantro grown in soil samples of the incubation experiment

Leptosol, Phaeozem, and Vertisol soils which were incubated for 14 days with the four different water types (influent, effluent, spiked-influent, spiked-effluent) in the soil incubation experiment were used for a greenhouse experiment. The soil samples were stored at 4°C in the dark and used for the cultivation of cilantro seedlings. The greenhouse experiment comprised 48 pots (3 soil types x 4 different water types x 4 replicates). Each pot contained 15 g of irrigated soil. Five cilantro (Hortaflor variety) seeds were sown per pot. Pots were irrigated daily with sterile deionized water to keep the soil moisture content at 80% water holding capacity. The pots were incubated for 6 weeks at 22°C (day) and 18°C (night) with a 12-hour photoperiod. The position of the pots was randomized daily. After 6 weeks of incubation, aboveground (mainly leaves) and belowground (roots) parts of the cilantro plants were separated carefully by placing plants on sterile petri dishes (Greiner Bio-one). Roots with adhering soil were carefully separated by hand (using gloves) from bulk soil. The aboveground and belowground parts of the plants were placed into sterile 2 mL Eppendorf tubes containing 750 µL autoclaved NaCl (0.3% w/v) and washed by vortexing for 1 min to remove the rhizosphere and detach bacteria from the plant surface. The supernatant was decanted and fresh NaCl solution was added. The process was repeated two times. The residual plant samples (both, “leaves” and “roots”) were transferred to sterile 15 mL tubes containing 5 mL autoclaved buffered peptone water (BPW; Sigma Aldrich). The plant material contained bacteria that were too strongly attached to the root and leave surfaces to be washed off and endophytic bacteria. All samples were transported on the day of sampling on cool bags (6°C) to the laboratory and stored at 4°C until further analysis.

### Pre-enrichment cultivation and isolation of RB from washed cilantro roots and leaves

Total and resistant heterotrophic bacteria as well as faecal indicator bacteria were cultivated from washed plant material. Small fractions (at least 0.5 g) of surface washed plant samples (stored in BPW) were incubated overnight in 1 mL autoclaved BPW at 37°C and 25°C to reactivate the plant associated bacteria. After overnight cultivation, serial dilutions were performed in 0.3% NaCl in sterile 96 wells plates as described above. Four technical replications were set up per sample. All dilutions (5 µL per well) were spotted on MH and R2A agar plates without and with antibiotics or biocidal compounds as described above. Following the spot assay, highest dilutions that showed growth in the presence of CIP were selected for cultivation of CIP RB. Therefore 100 µL of those dilutions were plated on MH and R2A agar containing CIP (1 and 4 µg mL^-1^). All agar plates were additionally supplemented with cycloheximide (100 µg mL^-1^) to avoid fungal growths. Plates were incubated under oxic conditions at 37°C (24 h) and 25°C (48 h - 4 days). At least five colonies of different abundant morphotypes were selected for analysis. Strains were purified by repeated streaking of single colonies on respective media with CIP. Additionally, overnight enrichment cultures in BPW (37°C) were spotted on TBX agar with and without CTX and SB agar with and without vancomycin. Plates were incubated as described above.

### Preservation and phylogenetic characterization of CIP resistant strains

Fresh biomass of pure cultures was suspended in new-born calf serum albumin (ThermoFisher Scientific, USA) and stored at -20°C for long-term preservation. For molecular analysis cell lysates were generated from biomass of pure cultures as described by Schauss et al.^57^ Genomic fingerprinting using the primer BOXA1R (Table S2) (BOX-PCR) was used to differentiate all strains at a strain level (genotyping). Analysis was performed according to Bartz *et al.* using BioNumerics version 8 (Applied Maths, Belgium).^58^ Strains with identical BOX-PCR patterns were assigned to one genotype. Few representatives of each genotype were phylogenetically identified by partial 16S rRNA gene sequencing. Amplification of the 16S rRNA gene was done using the primer system EUB9f and EUB1492r (Table S2) as described previously.^59,60^ Purification of PCR products and Sanger sequencing with primer EUB9f were performed by LGC Genomics (Berlin, Germany). Manual corrections of Sanger sequences were conducted using Molecular Evolutionary Genetics Analysis (MEGA) software Version 11 (MEGA 11).^61^ 16S rRNA gene sequence identities to next related type strains were obtained using the Basic Local Alignment Search Tool (BLAST; https://blast.ncbi.nlm.nih.gov/Blast.cgi) and the NCBI RefSeq 16S rRNA gene sequence database. Phylotypes were defined based on a phylogenetic tree and sequence identities. Respective analysis was performed in ARB (http://www.arb-home.de/) as described in detail in Franco *et al*.^62^ Based on the phylotype grouping, genotypes were defined per phylotype using genomic fingerprint data according to Franco *et al.*^62^ All sequences generated in this study were deposited in GenBank under accession numbers PV243452 - PV243571. The 16S rRNA gene sequences of the bacterial strains were used for comparative analysis with amplicon sequence variant (ASV) sequences published by Soufi *et al.* for the same soil incubation experiment as described in our study using MEGA 11.^50,61^

### Genetic diversity analysis of *Achromobacter* spp. strains

*Achromobacter* spp. strains were further genetically differentiated using the 765 bp DNA fragments of the gene *nrdA* coding for the alpha subunit of the ribonucleoside diphosphate reductase 1 as described previously.^63^ Amplification of partial *nrdA* sequences was carried out using primers and PCR condition as described by Spilker *et al.* and Amoureux *et al.*^63,64^ Sanger sequenced partial *nrdA* nucleotide sequences were manually checked by visual control of the chromatograms and shortened to a size of 765 nucleotide positions (5’-AAGAAGCCTACGTG … TGCATCGCAATCGCC-3’) according to the *nrdA* fragment used in the *Achromobacter* PubMLSTdatabase as locus *nrdA*_765 (http://pubmlst.org/achromobacter/).^65–67^ The sequences were processed and compared using MEGA 11. Based on the alignment performed with ClustalW, nucleotide differences per codon position were counted. The *nrdA*_765 nucleotide sequence fragments were deposited in the *Achromobacter* PubMLST database and allele numbers were assigned. Phylogenetic trees based on nucleotides and translated amino acids sequences of the partial *nrdA* sequences were additionally calculated in MEGA 11 using the Maximum likelihood method.^68^

### Antibiotic susceptibility testing (AST) and biocide tolerance tests

*Achromobacter* spp. strains cultivated from cilantro plants in the presence of CIP were used for AST. Testing was performed by the agar diffusion method using MH agar plates incubated at 37°C following the Clinical and Laboratory Standards Institute (CLSI) guidelines (M100-ED34). The following antibiotics were used to test for additional antimicrobial resistances: levofloxacin (LEV, 5 µg), CIP (5 µg), amikacin (AMK, 30 µg), meropenem (MER, 10 µg), imipenem (IMP, 10 µg), doripenem (DOR, 10 µg), piperacillin (PIP, 100 µg), piperacillin/tazobactam (PIT, 100/10 µg), and ceftazidime (CEF, 30 µg) (all Oxoid). *Achromobacter spanius* CCUG 47062^T^, *Pseudomonas aeruginosa* LMG 1242^T^, and *E. coli* ATCC 25922 were used as reference strains. *P. aeruginosa* specific CLSI breakpoints of inhibition zones were used to differentiate antibiotic resistant or susceptible phenotypes. Minimum inhibitory concentration (MIC) values of BAC-C12, DADMAC-C10, and ATMAC-C16 were determined with a broth microdilution assay following the CLSI guidelines (M100-ED34) using methods described previously.^8^ The following concentration ranges were tested, 0, 3.125, 6.25, 12.5, 25, 50, 100, and 200 μg BAC-C12 mL^-1^, 0, 0.3125, 0.625, 1.25, 2.5, 5, 10, and 20 μg DADMAC-C10 mL^-1^, and 0, 1.56, 3.125, 6.25, 12.5, 25, 50, and 100 μg ATMAC-C16 mL^-1^, respectively. The lowest concentration with no visible growth was considered as MIC value for the respective QAACs.

### Genetic characterization of the CIP resistance of *Achromobacter* spp. strains

Quinolone resistance determining regions (QRDRs) of the DNA gyrase (*gyrA*) and topoisomerase IV (*parC*) of CIP resistant *Achromobacter* spp. strains were PCR amplified using primers and PCR conditions as described previously.^69,70^ The PCR products of *gyrA* were Sanger sequenced with forward primers to determine point mutations leading to CIP resistance as reported previously in the Gram-negative bacteria.^69–71^ Amplification results for *parC* were unsatisfactory. The strains were further screened for the presence of plasmid mediated quinolone resistance (PMQR) genes *qnrB* and *qnrS* using primer systems and PCR conditions as described previously.^72,73^ Details of primer systems used are given in Table S2.

### Statistical analysis

Calculations of mean values and standard deviations were done in Microsoft Excel (Office 2016). The relative concentrations (relative abundance) of RB were calculated as the ratio of MH or R2A CFU counts per g or mL obtained from antibiotic or QAAC supplemented agar plates to agar plates without supplements. The Shapiro-Wilk test was applied to test the distribution of CFU counts for normality.^74^ If normal distribution was confirmed, the Brown-Forsythe test for equal variance was performed to check for differences between mean values.^75^ The one-way analysis of variance (ANOVA, parametric) was used to test significant difference between the means of CFU counts in the irrigated soils (incubated for 4 weeks) and soil samples prior to water addition. The Kruskal Wallis ANOVA based on rank was performed for non-parametric data.^76^ Two-way ANOVA was used to test interacting effects of wastewater (influent, effluent) with spiking levels (unspiked, spiked) on means of CFU counts per mL or g, respectively. Effects of wastewater types (influent, effluent), spiking levels (unspiked, spiked), and soil types (Leptosol, Phaeozem, Vertisol) on the relative abundance of RB were tested by a three-way ANOVA. All statistical tests were considered significant at *p*<0.05. SigmaPlot (Version 15, Systat, USA) was used for the described statistics. Principal component analyses (PCAs), permutational multivariate analysis of variance (one-way PERMANOVA), and analysis of similarities (one-way ANOSIM) were performed in PAST4 (https://folk.uio.no/ohammer/past/).

## Results

### Wastewater treatment reduced the abundance of total and resistant heterotrophic bacteria in the effluent compared to influent water, while spiking increased their abundance

The absolute abundance (CFU counts) of total bacteria cultivated on MH or R2A from influent was 5.85 (± 5.1) and 6.91 (± 6.2) log_10_ CFUs mL^-1^ (Table S3). Wastewater treatment significantly reduced the concentrations of both cultivated bacterial groups by 0.58 and 0.86 log_10_ CFUs mL^-1^ in the effluent compared to the influent (*p*<0.05). Spiking of both water types significantly increased the absolute abundance of MH grown bacteria by 0.8 and 0.3 log_10_ CFUs mL^-1^ (*p*<0.05), while bacteria cultivated on R2A showed only a slight but non-significant rise (Table S3).

Antibiotic (CIP, TRIM/SUL, and ERY/CLIN) resistant and BAC-C12 tolerant bacteria (summarized as resistant bacteria, RB) were cultivated from all wastewater types on MH and R2A (Table S3). Compared to the influent, the absolute abundance in the effluent was significantly lower for almost all targeted RB on MH and R2A, with a reduction of 0.27 to 1.54 log_10_ CFUs mL^-1^ (Table S3).

Relative abundances of RB cultivated on MH or R2A (fraction of CFU counts obtained from media with antimicrobials relative to CFU counts for media without antimicrobials) was also lower in the effluent compared to the influent. Only CIP RB cultivated on MH and BAC-C12 RB cultivated on R2A showed increased relative CFU counts in the effluent (Figure 1A).

**Figure 1.**
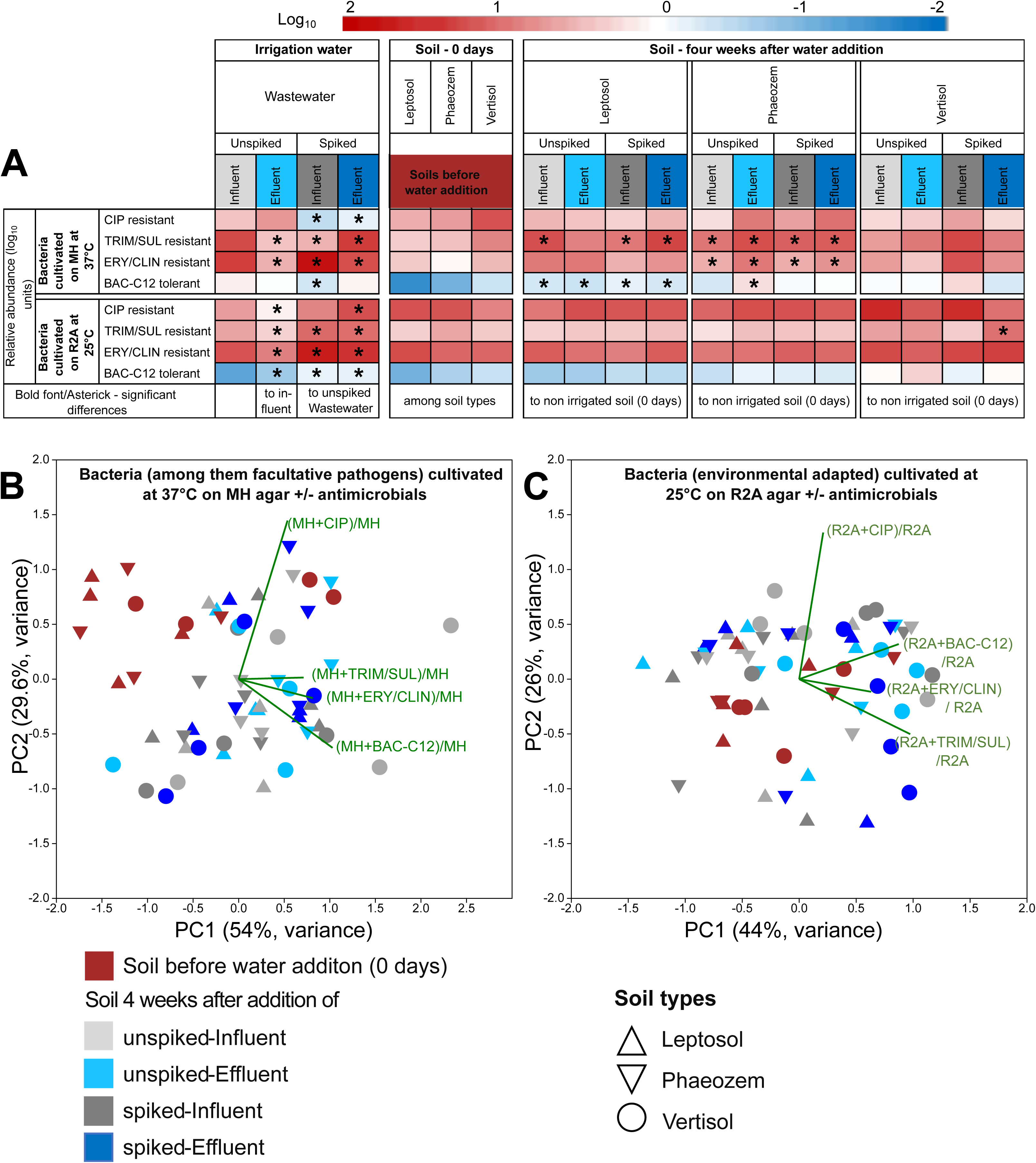
**(A)** Heat map showing mean of relative abundance of RB cultivated on MH (high nutrient conditions at 37°C, among them potential pathogens) and R2A (low nutrient conditions at 25°C, environmentally adapted) from the irrigation water types (unspiked-influent, unspiked-effluent, spiked-influent and spiked-effluent) and soil before (0 days) and after irrigation. Significant differences (*p*<0.05) were indicated with asterisks (*). PCA analysis of relative abundance of RB on **(B)** MH and **(C)** R2A in soil before (0 days) and after four weeks incubation following irrigation with both unspiked and spiked influent or effluent. Mean values of four technical replications (irrigation water types) or four biological replications (four fields per soil type) are presented. Each biological replicate based on four technical replications. For details see Figure S2.

Spiking influenced both, the relative abundance of MH and R2A grown RB, in both water types. In case of RB cultivated on MH, spiking led to a significant decrease in the relative abundance of CIP RB in both water types. Following spiking, the relative abundance of TRIM/SUL RB significantly decreased in the influent but significantly increased in the effluent. While the relative abundance of ERY/CLIN RB increased after spiking of both water types (Figure 1A, Figure S3).

After spiking of both wastewater types, relative abundance of almost all targeted RB cultivated on R2A was significantly higher with a non-significant rise for CIP RB in the spiked influent (Figure 1A, Figure S3).

From none of the water samples, independent of spiking, *E. coli* and enterococci as well as 3GCR *E. coli* and VRE could be cultivated by the spot assay cultivation approach.

### Mexican soils long-term irrigated with untreated wastewater contained cultivable RB

The Mexican soils used for the incubation experiment (before irrigation water addition, 0 days) contained total bacteria cultivated on MH and R2A in a range of 6.35 to 7.05 log_10_ CFUs g^-1^ (Table S4). The highest CFU counts were obtained for Leptosol, followed by Phaeozem and Vertisol samples (Table S4). A significant difference (*p*<0.05, One way ANOVA) was observed only between Leptosol (7.05 ± 0.1 log_10_ CFU g^-1^) and Vertisol (6.35 ± 0.6 log_10_ CFU g^-1^) for MH grown bacteria.

From all soil samples RB could be cultured on MH and R2A without significant differences in the absolute or relative abundance among the soils (Figures 1A, S3, Table S4). The absolute abundance of CIP, TRIM/SUL, and ERY/CLIN RB were in the ranged of 4.87 to 6.06 log_10_ CFU g^-1^, while abundance of BAC-C12 tolerant bacteria were lower (3.48 to 4.04 log_10_ CFU g^-1^) (Table S4). The highest relative abundance of RB on MH was observed in Vertisols (1.36 log_10_ units, CIP RB) and lowest in Leptosols (-1.57 log_10_ units, BAC-C12 RB) (Figure S3). Considering the relative abundance of RB on R2A, both highest and lowest relative abundances were observed in Leptosols (1.41 log_10_ units for ERY/CLIN RB, -1.08 log_10_ units for BAC-C12 RB) (Figure S3).

From none of the soil samples *E. coli* and enterococci as well as 3GCR *E. coli* and VRE could be isolated by the spot assay cultivation approach.

### RB abundance in four-week incubated soils – no difference between the irrigation water types but increased relative abundance compared to soils before water addition

Following the four weeks of soil incubation after water addition, the absolute abundance of total and resistant bacteria (MH and R2A CFU counts per g soil) showed no significant changes compared to soils before water addition (Figure S3, Table S4). Neither the soil type, nor the added water type (effluent versus influent; spiked versus unspiked) showed any effects on the abundance of cultivated bacteria (Three-way ANOVA, *p*>0.05).

Significant differences in the relative abundance of RB were only observed for bacteria cultivated on MH from soil samples after water addition and four weeks of incubation compared to soil samples before water addition (0 days) (Figure 1A). In Leptosol and Phaeozem, TRIM/SUL and ERY/CLIN resistant and BAC-C12 tolerant MH cultivated bacteria occurred at significantly higher relative abundance following water addition and incubation than before (*p*<0.05; Figure 1A, Figure S3). No significant differences were obtained for respective Vertisol samples. Again, water type and spiking showed no significant effects. For R2A cultivated bacteria, no significant effects were observed for relative abundances of RB in Leptosol and Phaeozem, but for TRIM/SUL RB in Vertisol irrigated with the spiked-effluent (Figure 1A, Figure S3).

PCA based on relative abundance data of RB showed a clear separation of soils before (0 days) and 4 weeks after water addition if RB were cultivated on MH at 37°C (Figure 1B). This was supported by one-way PERMANOVA (Euclidean distance, permutations = 9.999, R^2^ = 0.41, F = 2.5, p = 0.0001) and one-way ANOSIM (similarity index= Euclidean, permutations = 9.999, R = 0.137, p = 0.005) which both showed significant differences between soils before versus after water addition for MH cultivated RB.

Similar effects were not observed for the relative abundance of RB cultivated on R2A at 25°C (Figure 1C). Neither one-way PERMANOVA (Euclidean distance, permutations = 9,999, F = 1.20, p = 0.2) nor one-way ANOSIM (similarity index= Euclidean, R = −0.03, p = 0.7) detected significant differences between the two groups (soils before versus after water addition) for R2A cultivated RB.

### Cultivation of diverse RB from leaves/roots of cilantro plants grown in differently irrigated soils via pre-enrichment cultivation method

We aimed to study the transmission of RB present in soils irrigated with the different water types to cilantro plants. Neither from washed cilantro leaves nor from the washed roots samples, *E. coli* or enterococci could be cultivated after the performed non-selective pre-enrichment cultivation. However, RB (CIP, TRIM/SUL, and ERY/CLIN resistant and BAC-C12 tolerant bacteria) for all tested antimicrobials were cultivated from both leaves and root enrichment samples of cilantro plants grown in the different soil types after irrigation with all four water types (Table S5).

Among all targeted RB we decided to study only CIP RB in more detail. A high diversity of CIP RB (both on MH and R2A containing CIP) were cultivated from enrichment of leaves and roots of cilantro plants. A total of 187 colonies, representing 108 strains from leaves (at least 31 from each soil type) and 79 strains from roots (at least 23 from each soil type), were randomly selected for further characterization (Figure 2A). The CIP resistant strains isolated from both leaves and roots belonged to the phyla *Pseudomonadota* (formerly *Proteobacteria*; represented by *Alphaproteobacteria*, *Betaproteobacteria,* and *Gammaproteobacteria*), *Actinomycetota* (formerly *Actinobacteria*; represented by *Actinomycetia*), and *Bacillota* (formerly *Firmicutes*; represented by *Bacilli*).

**Figure 2.**
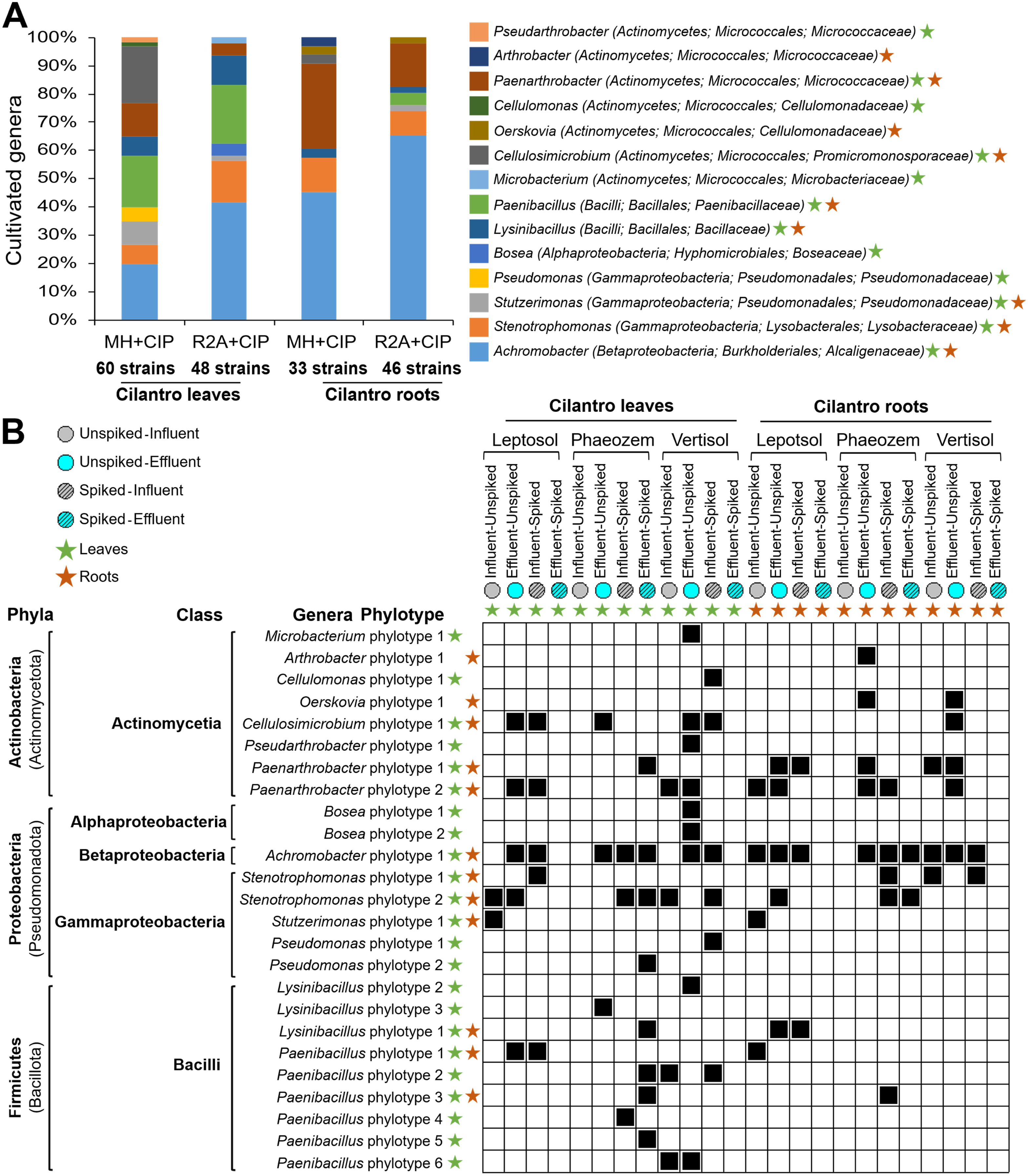
**(A)** Proportion of bacterial genera representing strains cultivated on MH and R2A plates containing CIP (1 and 4 µg mL^-1^) from leaves and roots of cilantro grown on soils irrigated with different wastewaters (influent, effluent, and both spiked). **(B)** Seriation analysis based on an absence-presence (0/1) matrix showing the presence of each phylotype cultivated on MH/R2A with CIP (1 and 4 µg mL^-1^) from cilantro. Cilantro plants were harvested 6 weeks after sowing seeds. The greenhouse experiment comprised 48 pots (3 soil types x 4 wastewater types x 4 replicates).

The CIP resistant strains from leaves were affiliated to 12 different genera, namely, *Achromobacter*, *Bosea*, *Cellulomonas*, *Cellulosimicrobium*, *Lysinibacillus*, *Microbacterium*, *Paenarthrobacter*, *Paenibacillus*, *Pseudarthrobacter*, *Pseudomonas*, *Stenotrophomonas,* and *Stutzerimonas* (Figure 2A, S4 A-F). The largest number of strains was assigned to the genus *Achromobacter* (29.6%, 32/108) followed by *Paenibacillus* (19.4%, 21/108), *Cellulosimicrobium* (11.1%, 12/108), *Stenotrophomonas* (10.2%, 11/108), and *Paenarthrobacter* (8.3%, 9/108). Similarly, CIP resistant strains from roots belonged to nine different genera namely, *Achromobacter*, *Arthrobacter*, *Cellulosimicrobium*, *Lysinibacillus*, *Oerskovia*, *Paenarthrobacter*, *Paenibacillus*, *Stenotrophomonas,* and *Stutzerimonas* (Figure 2B, S4 A-F). The largest numbers of strains were again assigned to the genus *Achromobacter* (57%, 45/79), followed by *Paenarthrobacter* (21.5%; 17/79) and *Stenotrophomonas* (10.1%, 8/79) (Figure 2A). Bacteria of those three genera were isolated from both leaves and roots on MH and R2A supplemented with CIP (MH/R2A with CIP; Figure 2B). *Pseudomonas* spp. strains were isolated only on MH CIP from leaves and *Paenibacillus* spp. and *Cellulosimicrobium* spp. strains were isolated from MH and R2A CIP, mainly from leaves and less frequently from roots. *Borsea* spp. strains were isolated only from R2A CIP from leaves.

### Detailed characterization of CIP resistant *Achromobacter* spp. strains of cilantro leaves and roots, with evidence suggesting soil origin

Due to the frequent isolation of CIP resistant *Achromobacter* spp. strains at both cultivation conditions (MH and R2A) and in both sample types (roots and leaves), *Achromobacter* spp. strains were selected for further characterization. Based on partial 16S rRNA gene sequences (approx. 1000 nt) *Achromobacter* spp. strains were placed in one phylogenetic cluster within the genus *Achromobacter* (Figure S4 D). The cluster contained different *Achromobacter* type strains that shared all 98.9-100% 16S rRNA gene sequence identity with our strains and among each other; among those *Achromobacter spanius*, *Achromobacter kerstersii*, *Achromobacter deleyi*, and *Achromobacter piechaudii*, and *Achromobacter marplatensis* (Figure S4 D). Considering the 16S rRNA gene region used by Soufi *et al*.^50^ for 16S rRNA gene-based amplicon sequencing of soil samples of the same incubation experiment studied here, the isolated *Achromobacter* spp. strains were assigned to two soil-derived *Achromobacter* ASVs (ASV2677 and ASV7979). The two ASVs had one nucleotide difference. ASV2677 was represented by 36 strains and shared 100% sequence identity with the 16S rRNA gene sequences of the type strains of *A. spanius*, *A. kerstersii*, *A. deleyi*, and *A. piechaudii*, while ASV 7979 was represented by 14 strains and was identical to the 16S rRNA gene sequence of *A. marplatensis* (Figures S5, S6). Further phylogenetic analysis based on partial *nrdA* nucleotide and amino acid sequences showed that the isolated *Achromobacter* spp. strains could be separated into three clusters (A to C) (Figure S5). A total of 40 strains were assigned to cluster A. All 40 strains were sequence identical to ASV2677 but represented 21 different *nrdA*_765 alleles. Six of those 21 alleles were previously assigned to strains of the species *A. spanius* (Figure S5). The remaining 18 isolated *Achromobacter* spp. strains were assigned to cluster B (6 strains) and cluster C (12 strains) represented further three and five different *nrdA_*765 alleles (Figure S5).

Interestingly, strains of clusters B and C either represented ASV2677 or ASV7979 (Figure S5). Further analysis based on genomic fingerprinting using BOX-PCR showed that the isolated *Achromobacter* spp. strains even represented a much higher genetic variability (Figures S6, S7). A total of 41 different *Achromobacter* BOX-genotypes were detected (Figure S6). Only two distinct genotypes (5a-1 and 5a-2) were found in both leaves and roots, and higher numbers of distinct genotypes were found specifically in roots (33 out of 41) compared to leaves (7 out of 41) (Figures S6, S7).

For strains of other genera, 16S rRNA gene sequence-based analyses and genomic fingerprinting also showed a high genetic variability among the CIP resistant strains (Figure 2 B; S8). Strains assigned to the genus *Paenarthrobacter* showed the highest genotypic diversity (19 different BOX-genotypes, Figure S8).

### Antibiotic susceptibility and QAAC tolerance of *Achromobacter* spp. strains isolated in the presence of CIP

The 60 *Achromobacter* spp. strains (23 from leaves, 37 from roots) cultivated on MH or R2A with 1 or 4 µg mL^-1^ CIP were further characterized with respect to their susceptibility to antibiotics and QAAC tolerance. All strains were susceptible to CIP and all other tested additional antibiotics (AMK, MER, IMP, DOR, PIP, PIT, and CEF). No nucleotide point mutation leading to changes of amino acids 83 or 87 were detected in the g*yrA* gene sequences of *Achromobacter* spp. strains. The PMQR genes *qnrB* or *qnrS* were not detected in any *Achromobacter* spp. strains. The MIC values for QAACs of the tested *Achromobacter* spp. strains were mostly in the ranges of 25-50 µg BAC-C12 mL^-1^, 5-10 µg DADMAC-C10 mL^-1^, and 50-100 µg ATMAC-C16 mL^-1^ (Figure S6).

## Discussion

Previous studies considering irrigation in agriculture had extensively focused to implications of irrigation with WWTP effluent replacing fresh water such as ground and surface water sources.^1,16,34^ Currently, only few studies have addressed the effects of the transition from irrigation with untreated wastewater to irrigation with WWTP effluent in agricultural soils, a scenario relevant for soils in the Mezquital valley in Mexico.^48–50^ We complemented the previously published cultivation independent microbiome (Soufi *et al*.)^50^ and chemical studies (Heyde *et al*.)^49^ with cultivation-based data.

We and Soufi *et al.*^50^ observed a typical reduction of CFU counts for heterotrophic bacteria between WWTP influent and effluent as reported in various studies of different countries.^77–82^ In the same manner, we found a significant reduction in the CFU counts of RB. While the direct comparison of our results with previous studies was hampered by the use of different media and tested antibiotics (including different antibiotic concentrations), similar reductions of the abundance of ARB, faecal coliforms, and total heterotrophic bacteria in WWTP effluent compared to influent were observed in several previous studies.^77,81–85^ However, while the relative abundance of most targeted RB decreased after wastewater treatment, the relative abundances of CIP RB cultivated on MH at 37°C and BAC-C12 tolerant bacteria cultivated on R2A at 25°C increased. This supported previous discussions that antimicrobial resistance could increase during the wastewater treatment processes.^42,79^

We showed that spiking of both, influent and effluent with antibiotics and QAACs significantly affected the abundance of cultivable bacteria and increased the relative abundance of cultivated RB. The overall rise in CFU counts of cultivated RB after spiking influent and effluent might indicate a metabolic capability of RB to cope with the spiked concentration of antibiotics and QAACs. RB might have grown faster than susceptible bacteria, with their growth being even further augmented by the utilisation of lysed cells of susceptible microbial population.^86–89^

We could not cultivate *E. coli* and enterococci from influent and effluent samples as well as from soil samples. However, at the time when the water samples were used to set up the soil incubation experiment both taxa could be cultured by a direct plating cultivation from influent and effluent waters (see also Suppl. Fig. S4).^50^ Long term storage of the water and soil samples before analysis seemed to have a negative effect on the cultivability of *E. coli* and enterococci in water and irrigated soils. For technical reasons both were stored at 6°C and studied after the irrigation experiment was finished. At the day of water addition, the concentration of cultivable *E. coli* in the influent, the effluent, spiked influent, and spiked effluent were 2.92, 1.86, 2.37, and 2.23 log_10_ CFUs mL^-1^, respectively.^50^ Based on those data the initial amount of cultivable *E. coli* present in the soils mixed with water was 2.0 and 1.0 log_10_ CFUs g^-1^ for influent and effluent water, 1.5 log10 CFU for spiked influent and 1.3 log_10_ CFUs g^-1^ for for spiked effluent. The limit of detection (LOD) for *E. coli* with the spot assay experiment was 2.3 log_10_ CFUs mL^-1^ (for water samples) and 3.3 log_10_ CFUs g^-1^ (soil samples). In addition, the spot assays were performed four months after the soil incubation experiment was set up. The absence of cultivable *E. coli* in our samples therefore aligns with decay rates reported by Murphy *et al.*^90^ for a similar soil greenhouse incubation experiment. This however highlights even more that a large proportion of the cultivated RB is part of the stable microbiome in the long-term irrigated Mexican soils.

Four weeks incubation of soils following irrigation, concentrations of RB were not significantly influenced by the type of water added (influent vs effluent) in any of the soil types, regardless of whether the added water was spiked with antibiotics and disinfectants or not. In line with our cultivation results, Soufi *et al.* also found no significant effect of the water type added (influent vs. effluent) upon the concentrations and patterns of ARGs in soil samples of the same incubation experiment that is addressed here.^50^ However spiking showed a significant effect which was not obtained in our study. An explanation for the lack of differences among influent and effluent irrigation is the fact that the effluent released from the Atotonilco WWTP might contain enough nutrients for the growth of bacteria, particularly on MH (at 37°C), compared to effluents from other conventional WWTPs.^49,50^ The specific response of bacteria cultivable on MH agar at 37°C showed that especially copiotrophic bacteria including potentially pathogenic wastewater derived bacteria may respond to the nutrient input. As pointed out above, a previous study showed that Mexican soils long-term irrigated with untreated wastewater had significant higher abundances of culturable *Gammaproteobacteria*, including potential pathogens, such as *Pseudomonas*, *Stenotrophomonas*, and *Acinetobacter* than rain fed soils.^32^

The lack of effects of spiking on the cultivability of RBs from irrigated soils may also be due to the adaptation of the soil bacterial communities to continuous pollutant exposure caused by the long-term irrigation history of the studied soils (at least 60 years) with untreated wastewater.^49,50^ The abundance of cultivable multidrug resistant gammaproteobacterial strains in long term wastewater irrigated soils may support this hypothesis.^32^ Other factors can also play a role for the observed lack of differences in the relative abundance of RBs. Soufi *et al*. showed lack of pronounced differences in the water-extractable concentrations of the antibiotics (AZI, CIP, ERY; CLIN, and TRIM) in soils between before and after water addition, except for SUL which was present in higher concentrations in soils irrigated with spiked versus unspiked water.^50^ As indicated by the lack of significant differences in the concentrations of antibiotics after irrigation with unspiked and spiked wastewater^50^, different factors (e.g., sorption/desorption) may influence antibiotic concentrations in soil which consequently could influence the abundance of RBs.^31^

Regarding the effect of wastewater irrigation on the abundance of RB in soils, a comparison of wastewater-irrigated soils to non-irrigated soils by Chen *et al*. showed no significant differences except for the relative abundance of sulfadiazine resistant bacteria, despite significantly higher concentrations of antibiotics (sulfamethoxazole, sulfadiazine, oxytetracycline) in wastewater irrigated soils.^91^ Similarly, Negreanu *et al.* also did not find significant difference between relative abundances of RB in soils irrigated with WWTP effluent versus freshwater (including well water).^92^

In contrast, our experiment revealed a significant increase in the relative abundance of RBs cultivable on MH at 37°C 4 weeks after wastewater addition to soils compared to the soils before water addition, especially in Leptosols and Phaeozems. MH agar is a medium used for the cultivation of fast-growing heterotrophs adapted to higher nutrient conditions.^52,53^ At an incubation temperature of 37°C this might include several potential pathogenic bacteria.^93^ In the studies of Chen *et al*.^91^ and Negreanu *et al.*^92^ Luria-Bertani (LB) agar, another nutrient rich agar, was used, but incubation was done at 30°C. Especially differences in media composition or temperature may have resulted the different study outcome. A study of manure-amended soils observed short term survival of manure-derived tetracycline-resistant bacteria in Danish farmland soils.^94^ That study again used LB agar and an incubation temperature of 25°C.^94^ Our finding that effects on the RB abundance in Leptosols and Phaeozems were stronger than in Vertisols is in line with observations that soil types influence the fate of RB and ARGs following the land application of sludge and manure.^95,96^ As shown by Soufi *et al.* the composition of the microbial communities in the studied soils showed clear differences among the soil types, and was less affected by the water applications.^50^ Our findings are in line with Seyoum *et al.* who showed that the occurrence and distribution of ARGs in treated wastewater-irrigated soils was independent of the irrigation water quality, but influenced by soil types, especially the soil clay content.^97^ Marano *et al.* showed that ARGs only correlated to irrigation water types in sandy but not in clay-rich soils, highlighting the significance of soil properties.^98^ While transient effects of wastewater irrigation could occur in the soil microbiomes, native microbial populations of soils are known to be resilient over time.^99^ Krause *et al.* and Obayomi *et al.* indicated that soil properties like the clay and the organic carbon content appeared to shape the bacterial communities, while irrigation water had negligible impact on the community composition and diversity.^100,101^

In this study, we also aimed to demonstrate the transmission of RB from wastewater exposed soils to fresh produce. Despite using a non-selective pre-enrichment approach, *E. coli*, 3GCR *E. coli*, enterococci, and VRE were not cultivated from cilantro samples (leaves and roots). This can be explained by the fact that those faecal indicator bacteria were already not viable in the stored soil samples. This may have also been the case for other wastewater associated bacteria as *Shewanella*, *Rheinheimera*, *Acinetobacter*, *Brevundimonas*, *Acidovorax*, *Cloacibacterium*, or *Arcobacter,* which were cultivated as CIP RB on MH from wastewater samples used to set up the soil incubation experiment (see Fig. S4). However, we showed that the tissue of cilantro plants germinated and cultivated in the soils of the incubation experiment contained a broad range of cultivable RB (CIP, TRIM/SUL and ERY/CLIN resistant, BAC-C12 tolerant bacteria). Those however can also represent members of the native soil microbiome which can have intrinsic or acquired resistances.^102,103^ Whether residues of antibiotics present in the differently irrigated soils or taken up by plants ^15,16,104–109^ affected the abundance of culturable RBs in plants is not clear yet. A comparative experiment with non-irrigated soils was not performed at that time but would have finally been required to answer this question.

Among all targeted RB (CIP, TRIM/SUL, and ERY/CLIN resistant, BAC-C12 tolerant bacteria), only CIP RB were selected for further characterization because fluoroquinolones are among the most prescribed antibiotics in Mexico City, and listed among the highest priority critically important antibiotics by the World Health Organisation.^110,111^ A large proportion of the CIP resistant strains from both leaves and roots represented soil derived *Achromobacter*.^50^ There was no indication for an acquired CIP resistance mechanism in our strains. The presence of plasmid mediated quinolone resistance genes was only scarcely reported for *Achromobacter* spp. strains cultivated from environments impacted by anthropogenic activities.^69^ Active efflux pumps (e.g., AxyABM) were discussed as the quinolone resistance mechanism in the clinical strains of *A. xylosoxidans*.^70,112^ Respective genes were not further analysed for the *Achromobacter* spp. strains obtained in our study. In the EUCAST (https://www.eucast.org/) database, MIC values of CIP for *Achromobacter* spp. strains are in the range of 0.25 to 16 µg mL^-1^. Therefore, it must be considered that the epidemiological cut-off (ECOFF) value of CIP for *Achromobacter* might be above the concentration which we added to MH and R2A for selective growth condition in this study.

*Achromobacter* are ecologically diverse bacteria found in various environments such as soil, water, waste and plants, but also in human clinical specimens.^113–118^ Most *Achromobacter* species are harmless, but few species of the genus *Achromobacter* have recently been considered as emerging pathogens in cystic fibrosis lungs.^119–121^ While *A. xylosoxidans* strains are most often isolated species from cystic fibrosis lungs and respiratory tracts, strains of *A. spanius* and *A. marplatensis* are also reported in cystic fibrosis. ^63,64,67,119, 122–124^ CIP resistance was most common among *Achromobacter* species in community-onset bloodstream infection during 19 years study of an Australian population.^125^ We showed that the cultivated *Achromobacter* spp. strains likely originate from the Mexican soils (identical to detected ASVs, see Soufi *et al.*)^50^ and actively colonized the plants grown in the soil.

Beside *Achromobacter* we cultivated several other CIP resistant strains assigned to the genera *Stenotrophomonas*, *Cellulosimicrobium, Paenibacillus,* and *Lysinibacillus* from both, leaves and roots, or only leaves (*Pseudomonas*) of cilantro plants. *Stenotrophomonas* and *Pseudomonas* were previously already discussed as potential harmful taxa detected in long term irrigated Mexican soils which may colonize plants.^32^ This was also confirmed by our study. Again, according to EUCAST database it must be considered that the *Stenotrophomonas* ECOFF value for CIP (16 µg mL^-1^) was above the applied concentrations used in our study, however for *Pseudomonas* the CIP ECOFF value was lower than the applied concentrations (ECOFF of 0.5 µg mL^-1^ for *P. aeruginosa)*.

All taxa which were detected are known to be plant associated, but also reported in the context of human infections. *Stenotrophomonas* spp. are found in soil and plants, and known for for their beneficial interaction with plants.^126–127^ Currently, only *Stenotrophomonas maltophilia* is known to cause human infections.^126^ *Cellulosimicrobium* spp. are commonly found in soil and sewage.^128^ Strains of the species *Cellulosimicrobium cellulans* and *Cellulosimicrobium funkei* were reported to cause infection to immunocompromised patients.^129,130^ Here, members of the genus *Cellulosimicrobium* were exclusively cultivated on MH containing CIP at 37°C, which are growth conditions for bacteria including potential human pathogens.^93^ However, there are studies which reported *Cellulosimicrobium* spp. strains with plant growth promoting and plant protecting properties.^131,132^ Members of genus *Paenibacillus* have been isolated from variety of environments such as soil, wastewater, and plants, and are known for promoting growth in plants.^133^ *Paenibacillus* infections are opportunistic only against immunocompromised individuals.^133,134^ *Lysinibacillus* spp. strains have been isolated from diverse environments ranging from pristine aquatic environments, soils, plants, food, insects, waste, and indoor air.^135^ Bacteria of this genus are considered non-pathogenic to humans, and have been rarely associated with infections, for instance, *Lysinibacillus fusiformis* linked to bacteremia.^136^ A recent study showed by genome sequencing of a plant growth promoting *Lysinibacillus capsici* strain isolated from leaf of ready-to-eat lettuce plants carried several pathogenicity and ARGs.^137^ If the occurrence of resistance genes in plant growth promoting endophytic bacteria is linked to environmental pollutions e.g., caused by wastewater irrigation needs further evaluation. The CIP RB in cilantro plants cultivated in Mexican soils require further evaluation of the complex mechanisms leading to their CIP resistance and abundance. Pathogenicity of the strains cultivated from cilantro in presence of CIP was not yet studied, but several of the strains cultivated from cilantro belong to genera that contain both plant growth promoting and human pathogenic features. As illustrated by Kleyn *et al.,* plant endophytes can share the different properties.^137^

### Conclusion and Prospective

We investigated the consequences of the transition from long-term irrigation of soils with untreated wastewater to irrigation with WWTP effluent for the abundance of cultivable RB in soil and fresh produce using the Mezquital Valley, Mexico, as model case. Our study showed no significant effects of irrigation water quality (WWTP influent vs effluent) on the abundance of RB in Mexican soils with a long-term history of irrigation with untreated wastewater, contrary to our hypothesis that switching to WWTP effluent would increase RB abundance. Spiking of antibiotics and QAACs increased the abundance of RB in WWTP influent and effluent, but unexpectedly not as hypothesised in the irrigated soils four weeks after irrigation. Inputs of bacteria, nutrients, and organic matter with added WWTP influent and effluent might have increased the relative abundance of RB in Leptosols and Phaeozems, but not in Vertisols. This supported our hypothesis that the soil type modulates the RB abundance. A key finding of our study was that a significant increase of RBs was only observed when a nutrient rich medium and 37°C were used for cultivation. This may indicate that either fast growing bacteria in the soils or wastewater derived RB bacteria were activated. Further studies of isolated bacterial strains from irrigation water and irrigated soils are required to understand this phenomenon. Isolation of soil-derived CIP RBs from cilantro plants showed potential RB selection and their transfer to fresh produce in agriculture system. This suggests that wastewater treatment alone does not replace the monitoring of wastewater irrigation health risks. Long-term field experiments are required in addition to laboratory experiments for assessing environmental risks of RB and ARGs during irrigation transition adopting the “One Health” approach. Our study illustrates that complementary studies using cultivation-independent and dependent studies of soil microbes, and studies of chemical pollutions are the key to understand the resistance spread linked to wastewater irrigation in more detail.

## Acknowledgements

This research was funded by the Deutsche Forschungsgemeinschaft (DFG, German Research Foundation) in the framework of the Research Unit FOR 5095: Pollutant – Antibiotic Resistance – Pathogen Interactions in a Changing Wastewater Irrigation System (project number 431531292) as well as the UNAM-DGAPA PAPIIT program (project number AG101524). The Mexican partners also acknowledge funding by UNAM-DGAPA-PAPIIT project No. AG101221 “Respuesta de las interacciones entre contaminantes, nutrientes y microorganismos al tratamiento del agua en los suelos del Valle del Mezquital y su efecto en la dispersión de multidrogoresistencia”. We further one thanks Dipendra Aryal for his support by the performance of spot assays to cultivated RBs from irrigated Mexican soils.

## References

1. M. F. Jaramillo and I. Restrepo, Wastewater reuse in agriculture: a review about its limitations and benefits, Sustainability, 2017, 9(**10**), 1734.

2. S. Ofori, A. Puškáčová, I. Růžičková and J. Wanner, Treated wastewater reuse for irrigation: pros and cons, Sci. Total Environ., 2021, 760, 143393.

3. M. L. Partyka and R. F. Bond, Wastewater reuse for irrigation of produce: A review of research, regulations, and risks, Sci. Total Environ., 2022, 828, 154385.

4. A. Christou, V. G. Beretsou, I. C. Iakovides, P. Karaolia, C. Michael, T. Benmarhnia, B. Chefetz, E. Donner, B. M. Gawlik, Y. Lee, T. T. Lim, L. Lundy, R. Maffettone, L. Rizzo, E. Topp and D. Fatta-Kassinos, Sustainable wastewater reuse for agriculture, Nat. Rev. Earth & Environ., 2024, 5(**7**), 504–521.

5. T. U. Berendonk, C. M. Manaia, C. Merlin, D. Fatta-Kassinos, E. Cytryn, F. Walsh, H. Bürgmann, H. Sørum, M. Norström, M. N. Pons, N. Kreuzinger, P. Huovinen, S. Stefani, T. Schwartz, V. Kisand, F. Baquero and J. L. Martinez, Tackling antibiotic resistance: the environmental framework, Nat. Rev. Microbiol., 2015, 13, 310–317.

6. A. M. Leiva, B. Piña and G. Vidal, Antibiotic resistance dissemination in wastewater treatment plants: a challenge for the reuse of treated wastewater in agriculture, Rev. Environ. Sci. Biotechnol., 2021, 20, 47–65.

7. S. Li, B. S. Ondon, S. H. Ho, J. Jiang and F. Li, Antibiotic resistant bacteria and genes in wastewater treatment plants: from occurrence to treatment strategies, Sci. Total Environ., 2022, 838, 156544.

8. D. Pulami, P. Kämpfer and S. P. Glaeser, High diversity of the emerging pathogen *Acinetobacter baumannii* and other *Acinetobacter* spp. in raw manure, biogas plants digestates, and rural and urban wastewater treatment plants with system specific antimicrobial resistance profiles, Sci. Total Environ., 2023, 859, 160182.

9. I. Michael, L. Rizzo, C. S. McArdell, C. M. Manaia, C. Merlin, T. Schwartz, C. Dagot and D. Fatta-Kassinos, Urban wastewater treatment plants as hotspots for the release of antibiotics in the environment: a review, Water Res. 2013, 47, 957–995.

10. F. Barancheshme and M. Munir, Strategies to combat antibiotic resistance in the wastewater treatment plants, Front. Microbiol., 2018, 8, 2603.

11. P. Chaturvedi, P. Shukla, B. S. Giri, P. Chowdhary, R. Chandra, P. Gupta and A. Pandey, Prevalence and hazardous impact of pharmaceutical and personal care products and antibiotics in environment: a review on emerging contaminants, Environ. Res., 2021, 194, 110664.

12. 12. L. M. Murray, A. Hayes, J. Snape, B. Kasprzyk-Hordern, W. H. Gaze and A. K. Murray, Co-selection for antibiotic resistance by environmental contaminants, npj Antimicrob. Resist., 2024, **2(1)**, 9.

13. L. J. Carter, B. Adams, T. Berman, N. Cohen, E. Cytryn, F. C. T. Elder, A. L. Garduño-Jiménez, D. Greenwald, B. Kasprzyk-Hordern, H. Korach-Rechtman, E. Lahive, I. Martin, E. ben Mordechay, A. K. Murray, L. M. Murray, J. Nightingale, A. Radian, A. E. Rubin, B. Sallach, D. Sela-Donenfeld, O. Skilbeck, H. Sleight, T. Stanton, I. Zucker and B. Chefetz, Co-contaminant risks in water reuse and biosolids application for agriculture, Environ. Pollut., 2025, 375, 126219.

14. H. M. Zhao, H. B. Huang, H. Du, J. Lin, L. Xiang, Y. W. Li, Q. Y. Cai, H. Li, C. H. Mo, J. S. Liu, M. H. Wong and D. M. Zhou, Intraspecific variability of ciprofloxacin accumulation, tolerance, and metabolism in Chinese flowering cabbage (*Brassica parachinensis*). J Hazard Mater. 2018, 349, 252–261.

15. H. Sidhu, G. O’Connor and J. Kruse, Plant toxicity and accumulation of biosolids-borne ciprofloxacin and azithromycin. Sci. Total Environ., 2019, 648, 1219–1226.

16. S. Lyu, L. Wu, X. Wen, J. Wang and W. Chen, Effects of reclaimed wastewater irrigation on soil-crop systems in China: a review. Sci. Total Environ., 2022, 813, 152531.

17. K. Blau, A. Bettermann, S. Jechalke, E. Fornefeld, Y. Vanrobaeys, T. Stalder, E. M. Top and K. Smalla, The transferable resistome of produce, mBio., 2018, 9(6), 10–1128.

18. K. Smalla, K. Cook, S. P. Djordjevic, U. Klümper and M. Gillings, Environmental dimensions of antibiotic resistance: assessment of basic science gaps, FEMS Microbiol. Ecol., 2018, 94(12), fiy195.

19. C. J. Reid, K. Blau, S. Jechalke, K. Smalla and S. P. Djordjevic, Whole Genome sequencing of *Escherichia coli* from store-bought produce, Front. Microbiol. 2020, 10, 3050.

20. X. van Doorslaer, J. Dewulf, H. van Langenhove and K. Demeestere, Fluoroquinolone antibiotics: an emerging class of environmental micropollutants, Sci. Total Environ., 2014, 500, 250–269

21. M. Conde-Cid, A. Núñez-Delgado, M. J. Fernández-Sanjurjo, E. Álvarez-Rodríguez, D. Fernández-Calviño and M. Arias-Estévez, Tetracycline and sulfonamide antibiotics in soils: presence, fate and environmental risks, Processes, 2020, 8(11), 1479.

22. A. Jurado, A. Margareto, E. Pujades, E. Vázquez-Suñé and M. S. Diaz-Cruz, Fate and risk assessment of sulfonamides and metabolites in urban groundwater. Environ. Pollut., 2020, 267, 115480.

23. W. Duan, H. Cui, X. Jia and X. Huang, Occurrence and ecotoxicity of sulfonamides in the aquatic environment: a review, Sci. Total Environ., 2022, 820, 153178.

24. Q. Yang, Y. Gao, J. Ke, P. L. Show, Y. Ge, Y. Liu, R. Guo and J. Chen, Antibiotics: an overview on the environmental occurrence, toxicity, degradation, and removal methods. Bioengineered, 2021, 12, 2967–3008.

25. V. A. Thai, V. D. Dang, N. T. Thuy, B. Pandit, T. K. Q. Vo and A. P. Khedulkar, Fluoroquinolones: fate, effects on the environment and selected removal methods, J. Clean. Prod., 2023, 418, 137844.

26. C. Yang and T. Wu, A comprehensive review on quinolone contamination in environments: current research progress, Environ. Sci. Pollut. Res., 2023, 30, 51814–51838.

27. C. Siebe and E. Cifuentes, Environmental impact of wastewater irrigation in central Mexico: an overview. Int. J. Environ. Health Res., 1995, 5(2), 161–173.

28. P. Dalkmann, M. Broszat, C. Siebe, E. Willaschek, T. Sakinc, J. Huebner, W. Amelung, E. Grohmann and J. Siemens, Accumulation of pharmaceuticals, enterococcus, and resistance genes in soils irrigated with wastewater for zero to 100 years in central Mexico, PLoS One, 2012, 7(9), e45397

29. B. J. Heyde, A. Anders, C. Siebe, J. Siemens and I. Mulder, Quaternary alkylammonium disinfectant concentrations in soils rise exponentially after long-term wastewater irrigation, Environ. Res. Lett., 2021, 16, 064045.

30. J. Siemens, G. Huschek, C. Siebe and M. Kaupenjohann, Concentrations and mobility of human pharmaceuticals in the world’s largest wastewater irrigation system, Mexico City-Mezquital Valley, Water Res., 2008, 42(8-9), 2124–2134.

31. P. Dalkmann, E. Willaschek, H. Schiedung, L. Bornemann, C. Siebe and J. Siemens, Long-term wastewater irrigation reduces sulfamethoxazole sorption, but not ciprofloxacin binding, in Mexican Soils, J. Environ. Qual., 2014, 43(3), 964–970.

32. M. Broszat, H. Nacke, R. Blasi, C. Siebe, J. Huebner, R. Daniel and E. Grohmann, Wastewater irrigation increases the abundance of potentially harmful Gammaproteobacteria in soils in Mezquital Valley, Mexico, Appl. Environ. Microbiol., 2014, 80(17), 5282–5291.

33. S. Slobodiuk, C. Niven, G. Arthur, S. Thakur and A. Ercumen, Does irrigation with treated and untreated wastewater increase antimicrobial resistance in soil and water: a systematic review, Int. J. Environ. Res. Public Health, 2021, 18, 12159.

34. M. H. Tawfik, H. Al-Zawaidah, J. Hoogesteger, M. Al-Zu’bi, P. Hellegers, J. Mateo-Sagasta and A. Elmahdi, Shifting Waters: The challenges of transitioning from freshwater to treated wastewater irrigation in the northern Jordan Valley, Water, 2023, 15(7), 1315.

35. H. Gao, L. Yang and X. Song, Effect of reclaimed water recharge on bacterial community composition and function in the sediment of the Chaobai River, China, J. Soils Sediments, 2023, 23(1), 526–538.

36. Y. Gao, G. Shao, S. Wu, W. Xiaojun, J. Lu and J. Cui, Changes in soil salinity under treated wastewater irrigation: a meta-analysis, Agric. Water Manag., 2021, 255, 106986.

37. P. Guedes, C. Martins, N. Couto, J. Silva, E. P. Mateus, A. B. Ribeiro and C. S. Pereira, Irrigation of soil with reclaimed wastewater acts as a buffer of microbial taxonomic and functional biodiversity, Sci. Total Environ., 2022, 802, 149671.

38. E. Hasan and A. Abu-Awwad, Impacts of long-term treated wastewater irrigation and rainfall on soil chemical and microbial indicators in semi-arid calcareous soils, Sustainability (Switzerland), 2025, 17(19), 8663.

39. M. Mola, P. G. Kougias, E. Statiris, P. Papadopoulou, S. Malamis and N. Monokrousos, Short-term effect of reclaimed water irrigation on soil health, plant growth and the composition of soil microbial communities, Sci. Total Environ., 2024, 949, 175107.

40. V. Moulia, N. Ait-Mouheb, G. Lesage, J. Hamelin, N. Wéry, V. Bru-Adan, L. Kechichian and M. Heran, Short-term effect of reclaimed wastewater quality gradient on soil microbiome during irrigation, Sci. Total Environ., 2023, 901, 166028.

41. W. Wang, Z. Wang, H. Ling, X. Zheng, C. Chen, J. Wang and Z. Cheng, Effects of reclaimed water irrigation on soil properties and the composition and diversity of microbial communities in northwest China, Sustainability (Switzerland), 2025, 17(1), 308.

42. B. J. Heyde, M. Braun, L. Soufi, K. Lüneberg, S. Gallego, W. Amelung, K. Axtmann, G. Bierbaum, S. P. Glaeser, E. Grohmann, R. Arredondo-Hernández, I. Mulder, D. Pulami, K. Smalla, C. Zarfl, C. Siebe and J. Siemens, Transition from irrigation with untreated wastewater to treated wastewater and associated benefits and risks, npj Clean Water, 2025, 8(1), 6.

43. A. Herre, C. Siebe and M. Kaupenjohann, Effect of irrigation water quality on organic matter, Cd and Cu mobility in soils of central Mexico. Water Sci. Technol., 2004, 50, 109–116.

44. M. Carrillo, G. C. Braun, C. Siebe, W. Amelung and J. Siemens, Desorption of sulfamethoxazole and ciprofloxacin from long-term wastewater-irrigated soils of the Mezquital Valley as affected by water quality, J. Soils Sediments, 2016, 16(3), 966–975.

45. ACCIONA. Atotonilco WWTP (Mexico), the world’s largest wastewater treatment plant, celebrates its first year in operation. [Internet]. 2018 [cited 2020 May 26]. Available from: https://www.acciona-agua.com/pressroom/in-depth/2018/july/atotonilco-wwtp-m%C3%A9xico-the-world-s-largest-wastewater-treatment-plant-celebrates-its-first-year-in-operation/

46. World Bank. Wastewater: From waste to resource-The case of Atotonilco De Tula, Mexico. Washington, DC: World Bank. [Internet]. 2018 [cited 2026 March 5]. Available from: https://openknowledge.worldbank.org/handle/10986/29493.

47. S. Chamizo-Checa, E. Otazo-Sánchez, A. Gordillo-Martínez, J. Suárez-Sánchez, C. González-Ramírez and H. Muñoz-Nava, Megacity wastewater poured into a nearby basin: looking for sustainable scenarios in a case study, Water, 2020, 12(3), 824.

48. F. R. A. Ziegler Rivera, B. Prado Pano, S. Guédron, L. Mora Palomino, C. Ponce de León Hill and C. Siebe Grabach, Impact of the change in irrigation practices from untreated to treated wastewater on the mobility of potentially toxic elements (PTEs) in soil irrigated for decades, J. Soils Sediments, 2023, 23(7), 2726–2743.

49. B. J. Heyde, K. Lüneberg, N. Hahn, M. Braun, J. Siemens and C. Siebe, Nutrients, metals, and carbon in soils irrigated with treated versus untreated wastewater, J. Plant Nutrition Soil Sci., 2026, 189(1), 131–142.

50. L. Soufi, I. D. Kampouris, K. Lüneberg, B. J. Heyde, D. Pulami, S. P. Glaeser, C. Siebe, J. Siemens, K. Smalla, E. Grohmann and S. Gallego, Wastewater-borne pollutants influenced antibiotic resistance genes and mobile genetic elements in the soil without affecting the bacterial community composition in a changing wastewater irrigation system, J Hazard Mater. 2025, 494, 138680.

51. N. Landero-Valenzuela, E. Hernández-Nataren, L. Chávez-Cerón, C. A. Granados-Echegoyen, N. Alonso-Hernández, N. Mayek-Pérez, F. M. Lara-Viveros, B. Ponce-Lira and N. Calderón-Cortés, Influence of Trichoderma species on the reduction of heavy metal levels in bean plants irrigated with wastewater: a case study from the Mezquital Valley, Hidalgo, Mexico, Renew. Agric. Food Syst., 2024, 39, 456–470.

52. CLSI. Clinical Laboratory Standards Institute. Performance Standards for Antimicrobial Susceptibility Testing. 34th ed. CLSI supplement M100. Wayne, PA: Clinical and Laboratory Standards Institute; 2020.

53. C. Fernández-Mazarrasa, O. Mazarrasa, J. Calvo, A. del Arco and L. Martínez-Martínez, High concentrations of manganese in Mueller-Hinton agar increase MICs of tigecycline determined by Etest. J. Clin Microbiol., 2009, 47(3), 827–829.

54. M. J. Allen, S. C. Edberg and D. J. Reasoner, Heterotrophic plate count bacteria - what is their significance in drinking water? Int. J. Food Microbiol., 2004, 92, 265–275.

55. D. J. Reasoner and E. E. Geldreich, A new medium for the enumeration and subculture of bacteria from potable water, Appl. Environ. Microbiol., 1985, 49(1), 1–7.

56. B. Jarvis, C. Wilrich and P. T. Wilrich, Reconsideration of the derivation of Most Probable Numbers, their standard deviations, confidence bounds and rarity values. J. Appl. Microbiol., 2010, 109(5), 1660–1667.

57. T. Schauss, S. P. Glaeser, A. Gütschow, W. Dott and P. Kämpfer, Improved detection of extended spectrum beta-lactamase (ESBL)-producing *Escherichia coli* in input and output samples of German biogas plants by a selective pre-enrichment procedure, PLoS One., 2015, 10(3), e0119791.

58. J. O. Bartz, J. Blom, H. J. Busse, J. B. Mvie, M. Hardt, P. Schubert, T. Wilke, A. Goessmann, G. Wilharm, J. Bender, P. Kämpfer and S. P. Glaeser, *Parendozoicomonas haliclonae* gen. nov. sp. nov. isolated from a marine sponge of the genus *Haliclona* and description of the family *Endozoicomonadaceae* fam. nov. comprising the genera *Endozoicomonas*, *Parendozoicomonas*, and *Kistimonas*, Syst. Appl. Microbiol., 2018, 41(2), 73–84.

59. S. P. Glaeser, O. Sowinsky, J. S. Brunner, W. Dott and P. Kämpfer, Cultivation of vancomycin-resistant enterococci and methicillin-resistant staphylococci from input and output samples of German biogas plants, FEMS Microbiol. Ecol., 2016, 92(3), fiw010.

60. D. Pulami, T. Schauss, T. Eisenberg, J. Blom, O. Schwengers, J. K. Bender, G. Wilharm, P. Kämpfer and S. P. Glaeser, *Acinetobacter stercoris* sp. nov. isolated from output source of a mesophilic German biogas plant with anaerobic operating conditions, Antonie van Leeuwenhoek, International Journal of General and Molecular Microbiology, 2021, 114(3), 489–500.

61. K. Tamura, G. Stecher and S. Kumar, MEGA11: Molecular evolutionary genetics analysis version 11. Mol. Biol. Evol., 2021, 38(7), 3022–3027.

62. A. Franco, C. Rückert, J. Blom, T. Busche, J. Reichert, P. Schubert, A. Goesmann, J. Kalinowski, T. Wilke, P. Kämpfer and S. P. Glaeser, High diversity of *Vibrio* spp. associated with different ecological niches in a marine aquaria system and description of *Vibrio aquimaris* sp. nov., Syst. Appl. Microbiol., 2020, 43(5), 126123.

63. T. Spilker, P. Vandamme and J. J. LiPuma, Identification and distribution of *Achromobacter* species in cystic fibrosis, J. Cyst. Fibros., 2013, 12, 298–301.

64. L. Amoureux, J. Bador, F. Bounoua Zouak, A. Chapuis, C. de Curraize and C. Neuwirth, Distribution of the species of *Achromobacter* in a French Cystic Fibrosis Centre and multilocus sequence typing analysis reveal the predominance of *A. xylosoxidans* and clonal relationships between some clinical and environmental isolates, J. Cyst. Fibros., 2016, 15(4), 486–494.

65. K. A. Jolley and M. C. J. Maiden, BIGSdb: Scalable analysis of bacterial genome variation at the population level, BMC Bioinformatics, 2010, 11(1), 595.

66. T. Spilker, P. Vandamme and J. J. LiPuma, A multilocus sequence typing scheme implies population structure and reveals several putative novel *Achromobacter* species, J. Clin. Microbiol., 2012, 50(9), 3010–3015.

67. S. S. Gade, N. Nørskov-Lauritsen and W. Ridderberg, Prevalence and species distribution of *Achromobacter* sp. cultured from cystic fibrosis patients attending the Aarhus centre in Denmark, J. Med. Microbiol., 2017, 66(5), 686–9.

68. J. Felsenstein, Evolutionary trees from DNA sequences: a maximum likelihood approach, J. Mol. Evol., 1981, 17(6), 368–376.

69. J. P. R. Furlan, D. G. Sanchez, I. F. L. Gallo and E. G. Stehling, Replicon typing of plasmids in environmental *Achromobacter* sp. producing quinolone-resistant determinants, APMIS, 2018, 126(11), 864–869.

70. A. Magallon, M. Roussel, C. Neuwirth, J. Tetu, A. C. Cheiakh, B. Boulet, V. Varin, V. Urbain, J. Bador and L. Amoureux, Fluoroquinolone resistance in *Achromobacter* spp.: substitutions in QRDRs of GyrA, GyrB, ParC and ParE and implication of the RND efflux system AxyEF-OprN, J. Antimicrob. Chemother., 2021, 76, 297–304.

71. D. C. Hooper and G. A. Jacoby, Mechanisms of drug resistance: quinolone resistance, Ann N Y Acad. Sci., 2015, 1354(1), 12–31.

72. V. Cattoir, L. Poirel, V. Rotimi, C. J. Soussy and P. Nordmann, Multiplex PCR for detection of plasmid-mediated quinolone resistance *qnr* genes in ESBL-producing enterobacterial isolates, J. Antimicrob. Chemother., 2007, 60(2), 394–397.

73. S. Alipour, M. Owrang, M. Rajabnia, M. Olfatifar, H. Kazemian and H. Houri, Prevalence of plasmid-mediated quinolone resistance genes in *Escherichia coli* isolates from colonic biopsies of Iranian patients with inflammatory bowel diseases: a cross-sectional study, Health Sci Rep. 2024, 7(12), e70204.

74. S. S. Shapiro and M. B. Wilk, An analysis of variance test for normality (complete samples), Biometrika, 1965, 52(3-4), 591–611.

75. M. B. Brown and A. B. Forsythe, Robust tests for the equality of variances, J. Am. Stat. Assoc., 1974, 69(346), 364–367.

76. W. H. Kruskal and W. A. Wallis, Use of ranks in one-criterion variance analysis, J. Am. Stat. Assoc., 1952, 47(260), 583–621.

77. L. Guardabassi, D. M. A. lo Fo Wong and A. Dalsgaard, The effects of tertiary wastewater treatment on the prevalence of antimicrobial resistant bacteria. Water Res., 2002, 36(8), 1955–1964.

78. M. Ferreira Da Silva, I. Vaz-Moreira, M. Gonzalez-Pajuelo, O. C. Nunes and C. M. Manaia, Antimicrobial resistance patterns in *Enterobacteriaceae* isolated from an urban wastewater treatment plant, FEMS Microbiol. Ecol., 2007, 60(1), 166–176.

79. Y. Zhang, C. F. Marrs, C. Simon and C. Xi, Wastewater treatment contributes to selective increase of antibiotic resistance among *Acinetobacter* spp., Sci. Total Environ., 2009, 407(12), 3702–3706.

80. A. Novo and C. M. Manaia, Factors influencing antibiotic resistance burden in municipal wastewater treatment plants, Appl. Microbiol. Biotechnol., 2010, 87(3), 1157–1166.

81. P. Gao, M. Munir and I. Xagoraraki, Correlation of tetracycline and sulfonamide antibiotics with corresponding resistance genes and resistant bacteria in a conventional municipal wastewater treatment plant, Sci. Total Environ., 2012, 421, 173–183.

82. S. Zhang, B. Han, J. Gu, C. Wang, P. Wang, Y. Ma, J. Cao and Z. He, Fate of antibiotic resistant cultivable heterotrophic bacteria and antibiotic resistance genes in wastewater treatment processes, Chemosphere, 2015, 135, 9–17

83. M. Munir, K. Wong and I. Xagoraraki, Release of antibiotic resistant bacteria and genes in the effluent and biosolids of five wastewater utilities in Michigan, Water Res., 2011, 45(2), 681–693.

84. L. Rizzo, C. Manaia, C. Merlin, T. Schwartz, C. Dagot, M. C. Ploy, I. Michael and D. Fatta-Kassinos, Urban wastewater treatment plants as hotspots for antibiotic resistant bacteria and genes spread into the environment: a review, Sci. Total Environ., 2013, 447, 345–360.

85. J. Wang, L. Chu, L. Wojnárovits and E. Takács, Occurrence and fate of antibiotics, antibiotic resistant genes (ARGs) and antibiotic resistant bacteria (ARB) in municipal wastewater treatment plant: an overview, Sci. Total Environ., 2020, 744, 140997.

86. G. Dantas, M. O. A. Sommer, R. D. Oluwasegun and G. M. Church, Bacteria subsisting on antibiotics, Science, 2008, 320(5872), 100-103.

87. E. Ortega Morente, M. A. Fernández-Fuentes, M. J. Grande Burgos, H. Abriouel, R. Pérez Pulido and A. Gálvez, Biocide tolerance in bacteria, Int. J. Food Microbiol., 2013, 162, 13–25.

88. O. Seungdae, K. Zohre, T. Despina, R. W. Michael, Minjae Kim, K. H. Janet, T. Madan, G. P. Spyros, C. S. Jim and T. K. Konstantinos, Microbial community degradation of widely used quaternary ammonium disinfectants, Appl. Environ. Microbiol., 2014, 80(19), 5892–5900.

89. B. Brycki, M. Waligórska and A. Szulc, The biodegradation of monomeric and dimeric alkylammonium surfactants, J. Hazard. Mater., 2014, 280, 627–635.

90. C. M. Murphy, D. L. Weller, C. A. Bardsley, D. T. Ingram, Y. Chen, D. Oryang, S. L. Rideout and L. K. Strawn, Survival of twelve pathogenic and generic *Escherichia coli* strains in agricultural soils as influenced by strain, soil type, irrigation regimen, and soil amendment, J. Food Prot., 2024, 87(10), 100343.

91. C. Chen, J. Li, P. Chen, R. Ding, P. Zhang and X. Li, Occurrence of antibiotics and antibiotic resistances in soils from wastewater irrigation areas in Beijing and Tianjin, China, Environ. Pollut., 2014, 193, 257–264.

92. Y. Negreanu, Z. Pasternak, E. Jurkevitch and E. Cytryn, Impact of treated wastewater irrigation on antibiotic resistance in agricultural soils, Environ. Sci. Technol., 2012, 46(9), 4800–4808.

93. E. Tahiri Vela, R. M, Gecaj, D. Pulami, A. Mehmeti, P. Kämpfer and S. P. Glaeser, Third generation cephalosporin-resistant (3GCR) *Escherichia coli* and biocide-tolerant heterotrophic bacteria in irrigation water used in *Capsicum annuum* cultivation areas in Kosovo, Sci. Rep., 2026.

94. G. Sengeløv, Y. Agersø, B. Halling-Sørensen, S. B. Baloda, J. S. Andersen and L. B. Jensen, Bacterial antibiotic resistance levels in Danish farmland as a result of treatment with pig manure slurry, Environ. Int., 2003, 28, 587–595.

95. J. Zhang, Q. Sui, J. Tong, H. Zhong, Y. Wang, M. Chen and Y. Wei, Soil types influence the fate of antibiotic-resistant bacteria and antibiotic resistance genes following the land application of sludge composts, Environ. Int., 2018, 118, 34–43.

96. Y. Zhang, D. Cheng, Y. Zhang, J. Xie, H. Xiong, Y. Wan, Y. Zhang, X. Chen and X. Shi, Soil type shapes the antibiotic resistome profiles of long-term manured soil, Sci. Total Environ., 2021, 786, 147361.

97. M. M. Seyoum, O. Obayomi, N. Bernstein, C. F. Williams and O. Gillor, Occurrence and distribution of antibiotics and corresponding antibiotic resistance genes in different soil types irrigated with treated wastewater, Sci. Total Environ., 2021, 782, 146835.

98. R. B. M. Marano, A. Zolti, E. Jurkevitch and E. Cytryn, Antibiotic resistance and class 1 integron gene dynamics along effluent, reclaimed wastewater irrigated soil, crop continua: elucidating potential risks and ecological constraints, Water Res., 2019, 164, 114906.

99. S. Frenk, Y. Hadar and D. Minz, Resilience of soil bacterial community to irrigation with water of different qualities under Mediterranean climate, Environ. Microbiol., 2014, 16(2), 559–569.

100. S. M. B. Krause, A. B. Dohrmann, O. Gillor, B. T. Christensen, I. Merbach and C. C. Tebbe, Soil properties and habitats determine the response of bacterial communities to agricultural wastewater irrigation, Pedosphere, 2020, 30(1), 146–158.

101. O. Obayomi, M. M. Seyoum, L. Ghazaryan, C. C. Tebbe, J. Murase, N. Bernstein and O. Gillor, Soil texture and properties rather than irrigation water type shape the diversity and composition of soil microbial communities, Appl. Soil Ecol., 2021, 161, 103867.

102. E. Harrelson, Q. Zeng, M. Gao, M. Toro and R. A. Blaustein, Multidrug resistance in bacteria associated with leafy greens and soil in urban agriculture systems, Front. Plant Sci., 2025, 16, 1664284.

103. D. Kiplimo, R. Mwirichia, W. A. Wicaksono, G. Berg and A. Abdelfattah, Intrinsic and acquired antimicrobial resistomes in plant microbiomes: implications for agriculture and public Health, J. Sustain. Agric. Environ., 2025, 4, 45–59.

104. P. A. Herklotz, P. Gurung, B. vanden Heuvel and C. A. Kinney, Uptake of human pharmaceuticals by plants grown under hydroponic conditions, Chemosphere, 2010, 78(11), 1416–1421.

105. T. Eggen, T. N. Asp, K. Grave and V. Hormazabal, Uptake and translocation of metformin, ciprofloxacin and narasin in forage- and crop plants, Chemosphere. 2011, 85(1), 26–33.

106. D. H. Kang, S. Gupta, C. Rosen, V. Fritz, A. Singh, Y. Chander, H. Murray and C. Rohwer, Antibiotic uptake by vegetable crops from manure-applied soils, J. Agric. Food Chem., 2013, 61(42), 9992–10001.

107. X. W. Li, Y. F. Xie, C. L. Li, H. N. Zhao, H. Zhao, N. Wang and J. F. Wang, Investigation of residual fluoroquinolones in a soil-vegetable system in an intensive vegetable cultivation area in Northern China, Sci. Total Environ., 2014, 468, 258–264.

108. M. B. M. Ahmed, A. U. Rajapaksha, J. E. Lim, N. T. Vu, I. S. Kim, H. M. Kang, S. S. Lee and Y. S. Ok, Distribution and accumulative pattern of tetracyclines and sulfonamides in edible vegetables of cucumber, tomato, and lettuce, J. Agric. Food Chem., 2015, 63(2), 398–405.

109. S. Hussain, M. Naeem, M. N. Chaudhry and M. A. Iqbal, Accumulation of residual antibiotics in the vegetables irrigated by pharmaceutical wastewater, Exposure and Health, 2016, 8(1), 107–115.

110. R. Sánchez-Huesca, A. Lerma, R. M. E. Guzmán-Saldaña and C. Lerma, Prevalence of antibiotics prescription and assessment of prescribed daily dose in outpatients from Mexico city, Antibiotics. 2020, 9(1), 38.

111. WHO. WHO list of critically important antimicrobials for human medicine (WHO CIA list). World Health Organization. 2019.

112. J. Bador, L. Amoureux, J. M. Duez, A. Drabowicz, E. Siebor, C. Llanes and C. Neuwirth, First description of an RND-type multidrug efflux pump in *Achromobacter xylosoxidans*, AxyABM, Antimicrob Agents Chemother., 2011, 55(10), 4912–4914.

113. T. Coenye, M. Vancanneyt, E. Falsen, J. Swings and P. Vandamme, *Achromobacter insolitus* sp. nov. and *Achromobacter spanius* sp. nov., from human clinical samples, Int. J. Syst. Evol. Microbiol., 2003, 53(6), 1819–1824.

114. G. Garrity, D. J. Brenner, N. R. Krieg and J. T. Staley, Bergey’s Manual of Systematic Bacteriology - Vol 2: The Proteobacteria, Part C - The Alpha-, Beta-, Delta- and Epsilonproteobacteria, Springer-Verlag New York Inc. 2005.

115. M. Gomila, L. Tvrzová, A. Teshim, I. Sedĺček, N. GonzáLez-Escalona, Z. ZdráHal, O. Šedo, F. GonzáLez, A. Bennasar, E. R. B. Moore, J. Lalucat and S. E. Murialdo, *Achromobacter marplatensis* sp. nov., isolated from a pentachlorophenol-contaminated soil, Int. J. Syst. Evol. Microbiol., 2011, 61(9), 2231–2237.

116. N. Kuncharoen, Y. Muramatsu, C. Shibata, Y. Kamakura, Y. Nakagawa and S. Tanasupawat, *Achromobacter aloeverae* sp. nov., isolated from the root of *Aloe vera* (L.) Burm.f., Int. J. Syst. Evol. Microbiol., 2017, 67(1), 37–41.

117. Busse H.J. and Auling G., Achromobacter. In Bergey’s manual of systematics of Archaea and Bacteria, Williams & Wilkins, Baltimore, MD., 2015, 1–14 p.

118. E. M. Rashad, D. M. Shaheen, A. A. Al-Askar, K. M. Ghoneem, A. A. Arishi, E. S. A. Hassan and W. E. I. A. Saber, Seed endophytic *Achromobacter* sp. F23KW as a promising growth promoter and biocontrol of rhizoctonia root rot of fenugreek, Molecules, 2022, 27(17), 5546.

119. B. Isler, T. J. Kidd, A. G. Stewart, P. Harris and D. L. Paterson, *Achromobacter* infections and treatment options, *Antimicrob. Agents Chemother*., 2020, 64, e01224–20.

120. M. Papalia, R. Figueroa-Espinosa, C. Steffanowski, C. Barberis, M. Almuzara, R. Barrios, C. Vay, G. Gutkind, J. di Conza and M. Radice, Expansion and improvement of MALDI-TOF MS databases for accurate identification of *Achromobacter* species, J. Microbiol. Methods., 2020, 172, 105889.

121. N. Bhowmik and R. M. Stubbendieck, *Achromobacter* spp.: Emerging pathogens in the cystic fibrosis lung, PLoS Pathog., 2025; 21(4), e1013067.

122. A. Coward, D. T. D. Kenna, C. Perry, K. Martin, M. Doumith and J. F. Turton, Use of *nrdA* gene sequence clustering to estimate the prevalence of different *Achromobacter* species among cystic fibrosis patients in the UK, J. Cyst. Fibros., 2016, 15, 478–485.

123. B. D. Edwards, J. Greysson-Wong, R. Somayaji, B. Waddell, F. J. Whelan, D. G. Storey, H. R. Rabin, M. G. Surette and M. D. Parkinsa, Prevalence and outcomes of *Achromobacter* species infections in adults with cystic fibrosis: a North American cohort study, J. Clin. Microbiol., 2017, 55(7), 2074–2085.

124. L. Veschetti, M. Boaretti, G. M. Saitta, R. Passarelli Mantovani, M. M. Lleò, A. Sandri and G. Malerba, *Achromobacter* spp. prevalence and adaptation in cystic fibrosis lung infection, Microbiol. Res., 2022, 263, 127177.

125. B. Isler, D. L. Paterson, P. N. A. Harris, W. Ling, F. Edwards, C. M. Rickard, T. J. Kidd, I. Gassiep and K. B. Laupland, *Achromobacter* species: an emerging cause of community-onset bloodstream infections, Microorganisms, 2022, 10(7), 1449.

126. 126. R. Ghosh, S. Chatterjee and N. C. Mandal, Stenotrophomonas, beneficial microbes in agro-ecology, 2020, Academic Press.

127. R. P. Ryan, S. Monchy, M. Cardinale, S. Taghavi, L. Crossman, M. B. Avison, G. Berg, D. van der Lelie and J. M. Dow, The versatility and adaptation of bacteria from the genus Stenotrophomonas, Nat. Rev. Microbiol., 2009, 7, 399–411.

128. P. Schumann and E. Stackebrandt, Cellulosimicrobium, in: Bergey’s manual of systematics of *Archaea* and *Bacteria*, 2015.

129. H. Petkar, A. Li, N. Bunce, K. Duffy, H. Malnick and J. J. Shah, *Cellulosimicrobium funkei*: First report of infection in a nonimmunocompromised patient and useful phenotypic tests for differentiation from *Cellulosimicrobium cellulans* and *Cellulosimicrobium terreum*, J. Clin. Microbiol., 2011, 49(3), 1175–1178.

130. H. Zhang, C. He, R. Tian and R. Wang, A case report of the differential diagnosis of *Cellulosimicrobium cellulans*-infected endocarditis combined with intracranial infection by conventional blood culture and second-generation sequencing, BMC Infect. Dis., 2020, 20(1), 963.

131. A. A. Eida, S. Bougouffa, I. Alam, M. M. Saad and H. Hirt, Complete genome sequence of the endophytic bacterium *Cellulosimicrobium* sp. JZ28 isolated from the root endosphere of the perennial desert tussock grass Panicum turgidum, Arch. Microbiol., 2020, 202(6), 1563-1569.

132. S. Thokchom, S. Mukherjee and D. S. Ningthoujam, Draft genome sequence of plant growth-promoting bacterial endophyte *Cellulosimicrobium cellulans* strain MUKR 5 isolated from Iris laevigata fisch from Manipur, India, Microbiol. Resour. Announc., 2025, e00892–25.

133. E. N. Grady, J. MacDonald, L. Liu, A. Richman and Z. C. Yuan, Current knowledge and perspectives of *Paenibacillus*: a review, Microbial Cell Factories, 2016, 15(1), 203.

134. 134. F. G. Priest, Paenibacillus, in: Bergey’s manual of systematics of Archaea and Bacteria, Wiley; 2015. p. 1–40.

135. 135. K. C. Failor, L. Tian, C. L. Monteil and B. A. Vinatzer, *Lysinibacillus*, in: Bergey’s manual of systematics of Archaea and Bacteria, Wiley; 2019. p. 1–62.

136. H. Morioka, K. Oka, Y. Yamada, Y. Nakane, H. Komiya, C. Murase, M. Iguchi and T. Yagi, *Lysinibacillus fusiformis* bacteremia: case report and literature review, J. Infect. Chemother., 2022, 28, 230–235.

137. M. S. Kleyn, M. O. Akinyemi, C. Bezuidenhout and R. Adeleke, Draft genome sequence of *Lysinibacillus capsici* NAVL5D with potential for plant growth promotion, 3 Biotech, 2025, **15**(10), 1-9.

