## Supporting information for "Antimicrobial resistant bacteria in wastewater-irrigated Mexican soils and transfer of resistant bacteria from irrigated soils to cilantro plants"

**to**

**Table S1**. Physiochemical wastewater parameters and concentrations of antibiotic and biocidal compound present in or spiked to irrigation water types (according to Soufi *et al.,* 2025)

| Parameters | WWTP influent | WWTP effluent | Spiked concentrations |
|---|---|---|---|
| pH | 7.4 | 7.4 | |
| EC | 1300 | 1300 | |
| BOD [mg/L] | 50 | 12 | |
| COD [mg/L] | 320 | 110 | |
| TOC [mg/L] | 65 | 34 | |
| Azithromycin [ng/L] | 0-1390 | 0-1330 | 695000 |
| Ciprofloxacin [ng/L] | 2350-3390 | 1940-2270 | 1695000 |
| Clindamycin [ng/L] | < DL | < DL | 60000*** |
| (Anhydro)Erythromycin [ng/L]* | 210 (<QL) -370 | 170-250 (<QL) | 185000 |
| Sulfamethoxazole [ng/L] | 3190-3990 | 2820-3780 | 1995000 |
| Trimethoprim [ng/L] | 1100-1300 | 1200-1400 | 650000 |
| Sum ATMACs [ng/L] | 170 | 98 | 84546 |
| ATMAC-C8 | 12 | 12 | 5770 |
| ATMAC-C10 | 11 | 10 | 5326 |
| ATMAC-C12 | 24 | 16 | 11766 |
| ATMAC-C14 | 13 | 11 | 6493 |
| ATMAC-C16 | 110 | 49 | 55191 |
| Sum BACs [ng/L] | 524 | 200 | 261650 |
| BAC-C8 | 9 | 9 | 4322 |
| BAC-C10 | 12 | 12 | 6115 |
| BAC-C12 | 318 | 95 | 158921 |
| BAC-C14 | 141 | 53 | 70559 |
| BAC-C16 | 19 | 14 | 9287 |
| BAC-C18 | 25 | 17 | 12446 |
| Sum DADMACs [ng/L] | 208 | 53 | 156452 |
| DADMAC-C8 | 0 (21)** | 0 (21)** | 10527 |
| DADMAC-C10 | 0 (21)** | 0 (21)** | 10527 |
| DADMAC-C12 | 0 (21)** | 0 (21)** | 10527 |
| DADMAC-C14 | 0 (21)** | 0 (21)** | 10527 |
| DADMAC-C16 | 0 (21)** | 0 (21)** | 10527 |
| DADMAC-C18 | 208 | 53 | 103817 |

*Anhydroerythromycin = degradation product of erythromycin

** concentration <DL, therefore the median of all QAACs was used as concentration

*** Clindamycin was not detected. The spiked concentration was calculated using the concentration found in Siemens *et al.,* (2008) (120 [ng/L]).

EC: electrical conductivity; BOD: biochemical oxygen demand; COD: chemical oxygen demand; TOC: total organic carbon; $N_{tot}$: total nitrogen; $NO_3^-$-N: Nitrate content; $NH_4^+$-N: ammonium content; ATMACs: alkyltrimethyl ammonium compounds; BACs: benzylalkyldimethyl ammonium compounds; DADMAC: dialkyldimethyl ammonium compounds, QL: quantification limit; DL: detection limit

**Table S2** Details of primer systems used in this study.

| Target | Primers | 5'-3' sequences | Primer concentration (µM) | Annealing temperature | Product size | Reference |
|---|---|---|---|---|---|---|
| Repetitive extragenic palindromic (REP) sequence | -BOXA1R | CTACGGCAAGGCGACGCTGACG | 1 µM | 53°C | Strain specific genomic fingerprints | Versalovic et al. (1994)<br>Glaeser et al. (2013) |
| 16S rRNA gene | EUB9F<br>EUB1492R | GAGTTTGATCMTGGCTCAG<br>CGGTTACCTTGTTACGACTT | 0.2 µM<br>0.2 µM | 54°C | ~1470 bp | Lane (1991) |
| *nrdA* | nrdA-F<br>nrdA-R | GAACTGGATTCCCGACCTGTTC<br>TTCGATTTGACGTACAAGTTCTGG | 0.2 µM<br>0.2 µM | 56°C | 954 bp | Spilker et al. (2012) |
| *gyrA* | gyrA-F1<br>gyrA-R1 | CCGGTATCGCTGGAAGAAGAGA<br>CCTGCTCGCTGCCGTCGTA | 0.2 µM<br>0.2 µM | 57°C | 436 bp | Magallon et al. (2021) |
| *parC* | parC-F<br>parC-R | ATCGGCGACGGCCTGAAGCC<br>CGGGATTCGGTATAACGCAT | 0.2 µM | 55°C | 273 bp | Furlan et al. (2018) |
| *qnrB* | qnrBm-F<br>qnrBm-R | GGMATHGAAATTCGCCACTG<br>TTTGCYGYYCGCCAGTCGAA | 0.2 µM<br>0.2 µM | 54°C | 264 bp | Cattoir et al. (2007);<br>Alipour et al. (2024) |
| *qnrS* | qnrSm-F<br>qnrSm-R | GCAAGTTCATTGAACAGGGT<br>TCTAAACCGTCGAGTTCGGCG | 0.2 µM<br>0.2 µM | 54°C | 428 bp | Cattoir et al. (2007);<br>Alipour et al. (2024) |

**Table S3**. Absolute abundance ($log_{10}$ CFU $mL^{-1}$) of bacteria cultivated on MH and R2A from wastewater used for incubation experiment. The data represented the mean CFU $mL^{-1}$ obtained after spotting four independent dilution series of each wastewater types (influent, effluent, and both spiked) on respective media plates (four technical replications, see Fig. S2). Asterisk (*) showed significant difference (Two-way ANOVA; p<0.05).

| Targeted heterotrophic culturable bacteria | Concentration in $log_{10}$ CFU $mL^{-1}$ [Mean (±Standard deviation)] | | | |
|---|---|---|---|---|
| | Unspiked | | Spiked | |
| | Influent | Effluent | Influent | Effluent |
| **Bacteria cultivated on MH (-/+ supplements) 37°C, 24 h** | | | | |
| Bacteria on MH | 5.85 (± 5.1) | 5.27 (± 4.6)* | 6.65 (± 6.0)* | 5.56 (± 0.0)* |
| CIP resistant bacteria (MH+CIP) | 4.36 (± 4.1) | 4.09 (± 3.5) | 4.18 (± 3.8) | 3.37 (± 3.2)* |
| TRI/SUL resistant bacteria (MH+TRI/SUL) | 5.28 (± 4.3) | 3.74 (± 3.3)* | 5.26 (± 0.0) | 5.09 (± 4.7)* |
| ERY/CLI resistant bacteria (MH+ERY/CLI) | 5.37 (± 4.7) | 3.90 (± 3.5)* | 6.53 (± 6.1)* | 4.95 (± 4.6)* |
| BAC-C12 tolerant bacteria (MH+BAC-C12) | 3.99 (± 3.3) | 3.26 (± 2.9)* | 4.30 (± 3.6)* | 3.64 (± 3.0)* |
| **Bacteria cultivated on R2A (-/+ supplements) 25°C, 48 h** | | | | |
| Bacteria on R2A | 6.91 (± 6.2) | 6.05 (± 5.3)* | 6.91 (± 6.2) | 6.19 (± 5.4) |
| CIP resistant bacteria (R2A+CIP) | 5.68 (± 5.4) | 4.16 (± 3.8)* | 5.87 (± 5.3) | 5.57 (± 5.2)* |
| TRI/SUL resistant bacteria (R2A+TRI/SUL) | 5.62 (± 5.4) | 4.41 (± 3.4)* | 6.08 (± 5.7)* | 5.33 (± 4.9)* |
| ERY/CLI resistant bacteria (R2A+ERY/CLI) | 6.26 (± 5.6) | 5.04 (± 5.0)* | 6.69 (± 5.9)* | 5.80 (± 5.0)* |
| BAC-C12 tolerant bacteria (R2A+BAC-C12) | 3.64 (± 3.0) | 3.29 (± 2.4)* | 4.73 (± 0.0)* | 4.00 (± 3.6)* |
| **Significance test to** | | to Influent | to Unspiked-Influent | to Unspiked-Effluent |

**Table S4.** Absolute abundance ($log_{10}$ CFU $g^{-1}$) of bacteria cultivated on MH and R2A from soils before irrigation (0 days) and after four weeks following irrigation with unspiked-influent, unspiked-effluent, spiked-influent and spiked-effluent, respectively. The data represented the mean CFU $g^{-1}$ obtained after spotting four independent dilution series of each composite soil sample obtained from four different points from respective soil types (Leptosol, Phaeozem, and Vertisol). Standard deviations are given in brackets. Significant differences are marked in bold ($p<0.05$, One-way ANOVA). Mean values of four biological replications (four fields per soil type). Each biological replicate based on four technical replications. For details see Figure S2.

| | **Soils - 0 days (before water addition)** | | | **Soils - four weeks after water addition** | | | | | | | | | | | |
|---|---|---|---|---|---|---|---|---|---|---|---|---|---|---|---|
| **Targeted cultivable heterotrophic bacteria** | **Absolute abundance in $log_{10}$ CFU $g^{-1}$ [Mean (±Standard deviation)]** | | | | | | | | | | | | | | |
| **Soil types** | **Leptosol** | **Phaeozem** | **Vertisol** | **Leptosol** | | | | **Phaeozem** | | | | **Vertisol** | | | |
| | | | | Unspiked | | Spiked | | Unspiked | | Spiked | | Unspiked | | Spiked | |
| | | | | Influent | Effluent | Influent | Effluent | Influent | Effluent | Influent | Effluent | Influent | Effluent | Influent | Effluent |
| **Bacteria cultivated on MH (-/+ supplements) 37°C, 24 h** | | | | | | | | | | | | | | | |
| Bacteria on MH | **7.05 (± 0.1)** | 6.81 (± 0.1) | **6.35 (± 0.6)** | 6.5 (± 0.3) | 6.5 (± 0.2) | 6.6 (± 0.3) | 6.6 (± 0.2) | 6.5 (± 0.4) | 6.0 (± 0.4) | 6.3 (± 0.3) | 6.3 (± 0.4) | 6.7 (± 0.3) | 6.7 (± 0.4) | 6.4 (± 0.6) | 6.7 (± 0.3) |
| CIP resistant bacteria (MH+CIP) | 5.69 (± 0.3) | 5.58 (± 0.4) | 5.71 (± 0.2) | 5.1 (± 0.6) | 5.0 (±0.6) | 5.0 (± 0.7) | 5.1 (± 0.6) | 4.8 (± 0.4) | 5.1 (± 0.9) | 5.1 (± 0.8) | 5.4 (± 1.0) | 5.0 (± 0.7) | 5.1 (± 0.7) | 5.1 (± 0.8) | 5.0 (± 0.8) |
| TRI/SUL resistant bacteria (MH+TRI/SUL) | 5.44 (± 0.2) | 5.32 (± 0.1) | 5.32 (± 0.3) | 5.9 (± 0.1) | 5.5 (± 0.4) | 5.7 (± 0.3) | 6.0 (± 0.2) | 5.6 (± 0.6) | 5.4 (± 0.5) | 5.6 (± 0.4) | 5.6 (± 0.6) | 5.5 (± 0.5) | 5.7 (± 0.2) | 5.8 (± 0.1) | 5.7 (± 0.1) |
| ERY/CLI resistant bacteria (MH+ERY/CLI) | 5.47 (± 0.4) | 4.87 (± 0.2) | 4.90 (± 0.3) | 5.6 (±0.5) | 5.6 (± 0.2) | 5.2 (± 0.2) | 5.4 (± 0.3) | 5.2 (± 0.6) | 5.1 (± 0.6) | 5.0 (± 0.2) | 5.1 (± 0.6) | 5.5 (± 0.4) | 5.1 (± 0.6) | 5.7 (± 0.2) | 5.6 (± 0.2) |
| BAC-C12 tolerant bacteria (MH+BAC-C12) | 3.48 (± 0.6) | 3.71 (± 0.5) | 3.95 (± 0.2) | 4.2 (±0.1) | 4.1 (± 0.2) | 4.4 (± 0.7) | 4.3 (± 0.1) | 4.2 (± 0.5) | 4.3 (± 0.6) | 4.3 (± 0.5) | 4.3 (± 0.4) | 4.6 (± 0.4) | 4.6 (± 0.6) | 4.7 (± 0.4) | 4.4 (± 0.4) |
| **Bacteria cultivated on R2A (-/+ supplements) 25°C, 48 h** | | | | | | | | | | | | | | | |
| Bacteria on R2A | 6.64 (± 0.3) | 6.35 (± 0.5) | 6.46 (± 0.4) | 6.7 (± 0.5) | 6.6 (± 0.5) | 6.6 (± 0.5) | 6.5 (± 0.5) | 6.4 (± 0.4) | 6.1 (± 0.6) | 6.2 (± 0.4) | 6.2 (± 0.2) | 6.2 (± 0.7) | 6.4 (± 0.7) | 6.2 (± 0.6) | 6.3 (± 0.6) |
| CIP resistant bacteria (R2A+CIP) | 6.01 (± 0.5) | 5.69 (± 0.3) | 5.44 (± 0.5) | 5.8 (± 1.2) | 6.0 (± 1.0) | 5.9 (± 1.2) | 5.8 (± 1.0) | 5.6 (± 0.8) | 5.5 (± 0.9) | 5.5 (± 0.7) | 5.5 (± 0.9) | 5.9 (± 0.8) | 5.8 (± 0.9) | 5.9 (± 0.9) | 5.4 (± 1.0) |
| TRI/SUL resistant bacteria (R2A+TRI/SUL) | 5.02 (± 0.1) | 5.05 (± 0.1) | 4.88 (± 0.2) | 5.1 (± 0.5) | 5.1 (± 0.4) | 5.0 (± 0.5) | 5.0 (± 0.3) | 4.6 (± 0.7) | 4.6 (± 0.8) | 4.7 (± 0.6) | 4.7 (± 0.6) | 5.0 (± 0.3) | 5.3 (± 0.6) | 4.6 (± 0.4) | 5.4 (± 0.6) |
| ERY/CLI resistant bacteria (R2A+ERY/CLI) | 6.06 (± 0.6) | 5.54 (± 0.5) | 5.65 (± 0.6) | 6.0 (± 0.5) | 5.7 (± 0.5) | 5.6 (± 0.5) | 5.8 (± 0.6) | 5.4 (± 0.5) | 5.2 (± 0.7) | 5.4 (± 0.5) | 5.4 (± 0.8) | 5.6 (± 0.7) | 5.7 (± 0.5) | 5.7 (± 0.6) | 5.8 (± 0.3) |
| BAC-C12 tolerant bacteria (R2A+BAC-C12) | 3.56 (± 0.4) | 3.72 (± 0.4) | 4.04 (± 0.1) | 3.9 (± 0.4) | 3.9 (± 0.4) | 4.1 (± 0.9) | 4.0 (± 0.5) | 3.9 (± 0.4) | 3.9 (± 0.5) | 3.9 (± 0.7) | 4.0 (± 0.5) | 4.3 (± 0.3) | 4.8 (± 0.9) | 4.1 (± 0.3) | 4.3 (± 0.7) |

**Table S5**: Phylogenetic overview of bacterial strains cultivated on MH/R2A plates containing CIP from leaves and roots of cilantro grown in soil types (Leptosol, Phaeozem, Vertisol) irrigated with unspiked-influent, unspiked-effluent, spiked-influent, and spiked-effluent wastewater. Similarity values represented ranges of pairwise sequence similarities of phylotype strains to next related type strain placed within a phylotype cluster. Number inside the bracket represented the number of strains sharing identical DNA fingerprint pattern (BOX-PCR).

| | | **Cilantro tissue** | **Cilantro leaves** | | | | | | | | | | | | **Cilantro roots** | | | | | | | | | | | |
|---|---|---|---|---|---|---|---|---|---|---|---|---|---|---|---|---|---|---|---|---|---|---|---|---|---|---|
| | | **Soil types** | **Leptosol** | | | | **Phaeozem** | | | | **Vertisol** | | | | **Leptosol** | | | | **Phaeozem** | | | | **Vertisol** | | | |
| | | **Spiking of wastewaters** | Unspiked | | Spiked | | Unspiked | | Spiked | | Unspiked | | Spiked | | Unspiked | | Spiked | | Unspiked | | Spiked | | Unspiked | | Spiked | |
| **Genus** | **Phylotype** | **Pairwise 16S rRNA gene similarity to type strains of given species** | Influent | Effluent | Influent | Effluent | Influent | Effluent | Influent | Effluent | Influent | Effluent | Influent | Effluent | Influent | Effluent | Influent | Effluent | Influent | Effluent | Influent | Effluent | Influent | Effluent | Influent | Effluent |
| *Achromobacter* | 1 | 99–100%, *A. spanius/A. marplatensis/A. deleyi/A. kerstersii/A. piechaudii* | | 6 (4) | 2 (1) | | | 7 (5) | 6 (4) | 1 | | 9 (7) | 1 | | 2 | 3 | 10 (2) | | | 8 (1) | 3 | 3 | 6 (1) | 3 | 7 (2) | |
| *Bosea* | 2 | 98.8%, B. lathyri | | | | | | | | | | 1 | | | | | | | | | | | | | | |
| | 1 | 99.2-99.4%, *B. robiniae/B. thiooxidans/B. spartocytisi* | | | | | | | | | | 1 | | | | | | | | | | | | | | |
| *Cellulomonas* | 1 | 99.1%, *C. pakistanensis* | | | | | | | | | | | 1 | | | | | | | | | | | | | |
| *Cellulosimicrobium* | 1 | 98.5-99.7%, *C. funkei* | | 3 (2) | 4 (3) | | | 1 | | | | 2 (1) | 2 (2) | | | | | | | | | | | 1 | | |
| *Lysinibacillus* | 1 | 99.5-100%, *L. fusiformis* | | | | | | | | 5 (4) | | | | | | 1 | 1 | | | | | | | | | |
| | 2 | 99.5%, *L. xylanilyticus* | | | | | | | | | | 2(1) | | | | | | | | | | | | | | |
| | 3 | 99.8-100%, *L. capsici* | | | | | | 2 | | | | | | | | | | | | | | | | | | |
| *Microbacterium* | 1 | 99.3%, *M. lacus* | | | | | | | | | | 1 | | | | | | | | | | | | | | |
| *Paenarthrobacter* | 1 | 99.2-100%, *P. nicotinovorans* | | | | | | | | 2 (1) | | | | | | 3 (1) | 2 | | | 1 | | | 1 | 1 | | |
| | 2 | 98.6-100%, *P. aurescens/P. nitroguajacolicus* | | 1 | 2 (2) | | | | | | 2 | 2 (1) | | | 2 | 1 | | | | 3 | 1 | | | 2 (1) | | |
| *Paenibacillus* | 1 | 98.2-100%, *P. etheri* | | 1 | 1 | | | | | | | | | | 1 | | | | | | | | | | | |
| | 2 | 97%, *P. auburnensis/P. typhae* | | | | | | | | 1 | 1 | | 2 | | | | | | | | | | | | | |
| | 3 | 99.4-99.5%, *P. taohuashanense* | | | | | | | | 1 | | | | | | | | | | | 1 | | | | | |
| | 4 | 99.5%, *P. cucumis/P. polysaccharolyticus* | | | | | | | 1 | | | | | | | | | | | | | | | | | |
| | 5 | 99.4-99.7%, *P. xylanexedens/P. amylolyticus* | | | | | | | | 1 | | | | | | | | | | | | | | | | |
| | 6 | 99.2-99.6%, *P. ottowii/P. brasilensis* | | | | | | | | | 11 (8) | 1 | | | | | | | | | | | | | | |
| *Pseudarthrobacter* | 1 | 99.2%, *P. defluvii/P. niigatensis* | | | | | | | | | | 1 | | | | | | | | | | | | | | |

| | | | | | | | | | | | | | | | | | | | | | | | | | |
|---|---|---|---|---|---|---|---|---|---|---|---|---|---|---|---|---|---|---|---|---|---|---|---|---|---|
| *Pseudomonas* | 2 | 100%, *P. atacamensis/P. iranenesis/P. koreensis* | | | | | | | | 2 (1) | | | | | | | | | | | | | | | |
| | 1 | 99.7%, *P. citrulli* | | | | | | | | | | | 1 | | | | | | | | | | | | |
| *Stenotrophomonas* | 1 | 99.6-100%, *S. lactitubi* | | | 2 (1) | | | | | | | | | | | | | | | | **1** | | 1 | | 1 |
| | 2 | 99.6-100%, *S. rhizophila/S. nematodicola* | 1 (1) | 2 (1) | | | | | 2 | 2 | 1 | | 1 (1) | | | 2 (1) | | | | | 2 (1) | 1 | | | |
| *Stutzerimonas* | 1 | 99-100%, *S. chloritidismutans/S. kunmingensis* | 6 (4) | | | | | | | | | | | | 1 | | | | | | | | | | |
| *Arthrobacter* | 1 | 97.6%, *A. crystallopoietes* | | | | | | | | | | | | | | | | | | 1 | | | | | |
| *Oerskovia* | 1 | 99.9-100%, *O. jenensis* | | | | | | | | | | | | | | | | | | 1 | | | | 1 | |
| **Total strains** | | | **108** | | | | | | | | | | | | **79** | | | | | | | | | | |

### Supplementary figures

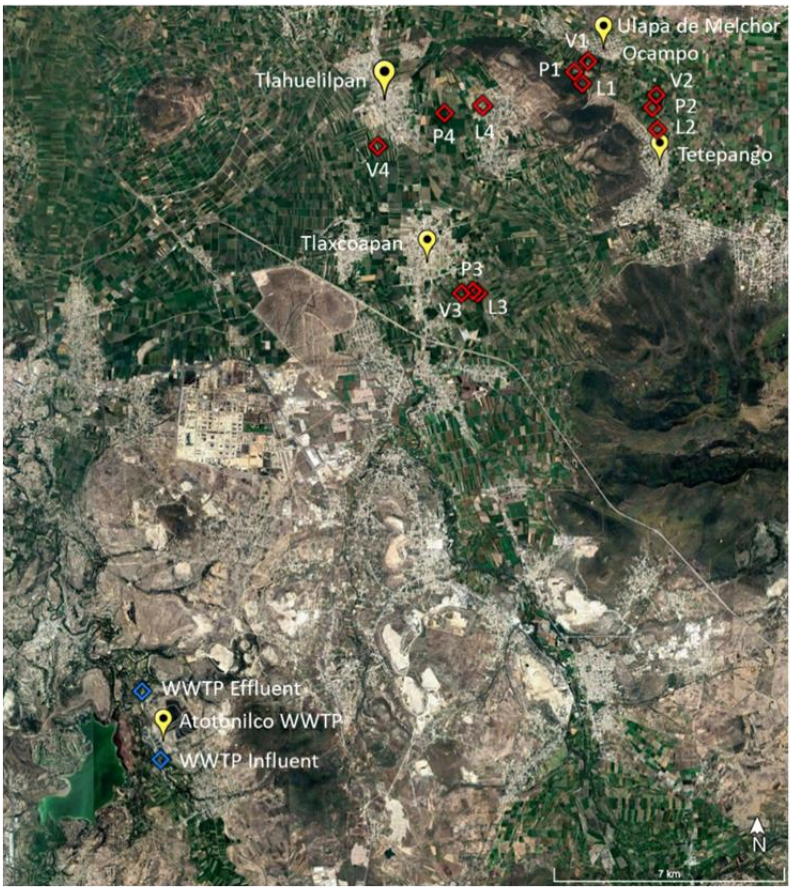

| Location | Years of UWW irrigation | Soil type | Coordinates | |
|---|---|---|---|---|
| **Ulapa de Melchor Ocampo** | 111 | Leptosol (L1) | 20° 8' 9.5" N | 99° 10' 31.3" W |
| | | Phaeozem (P1) | 20° 8' 13.6" N | 99° 10' 23.5" W |
| | | Vertisol (V1) | 20° 8' 30.9" N | 99° 10' 25.3" W |
| **Between Tetepango and Ulapa** | 60 | Leptosol (L2) | 20° 7' 24.5" N | 99° 9' 13.3" W |
| | | Phaeozem (P2) | 20° 7' 45.7" N | 99° 9' 18.4" W |
| | | Vertisol (V2) | 20° 7' 58" N | 99° 9' 14.3" W |
| **Tlaxcoapan (Bojayito Chico)** | 92 | Leptosol (L3) | 20° 4' 44.9" N | 99° 12' 20.1" W |
| | | Phaeozem (P3) | 20° 4' 47" N | 99° 12' 25" W |
| | | Vertisol (V3) | 20° 4' 45" N | 99° 12' 37" W |
| **Tlahuelilpan** | 101 | Leptosol (L4) | 20° 7' 28.4" N | 99° 12' 36.3" W |
| | | Phaeozem (P4) | 20° 7' 40.4" N | 99° 12' 54.5" W |
| | | Vertisol (V4) | 20° 7' 8.24" N | 99° 14' 3.78" W |

**Figure S1**. Geographical area, location, coordinates and years of untreated wastewater (UWW) irrigation of the agricultural fields used for the study (L1: Leptosol field 1; L2: Leptosol Field 2; L3: Leptosol Field 3; L4: Leptosol Field 4; P1: Phaeozem field 1; P2: Phaeozem Field 2; P3: Phaeozem Field 3; P4: Phaeozem Field4; V1: Vertisol field 1; V2: Vertisol Field 2; V3: Vertisol Field 3; V4: Vertisol Field 4). Adapted from Soufi *et al.,* (2025)

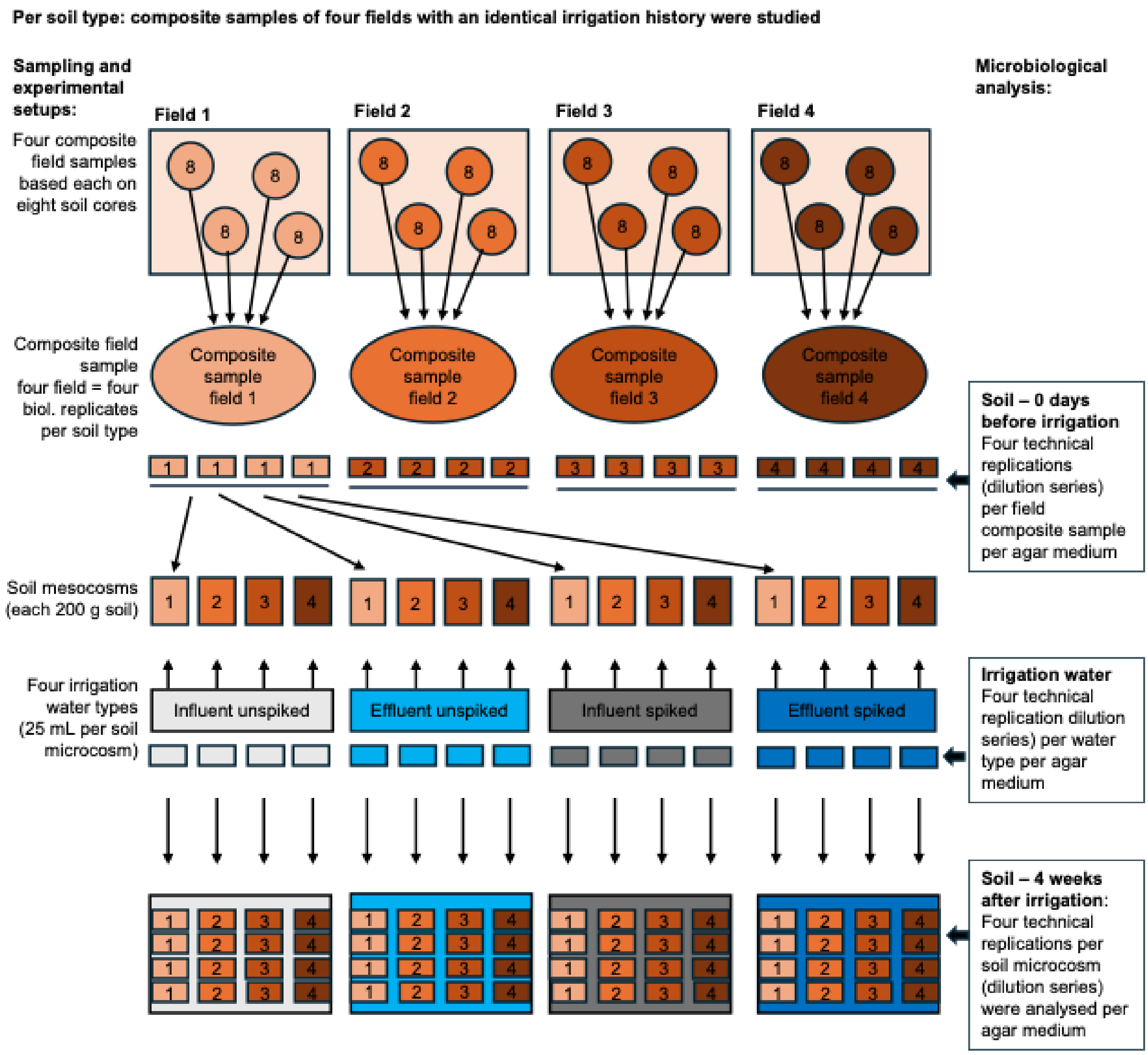

**Figure S2** Overview of sample collection in Mexican fields, set up of the incubation experiment, and performed microbiological analysis

| | | | Irrigation water | | | | Soil - 0 days | | |
|---|---|---|---|---|---|---|---|---|---|
| | | | Wastewater | | | | Leptosol | Phaeozem | Vertisol |
| | | | Unspiked | | Spiked | | | | |
| | | | Influent | Effluent | Influent | Effluent | Non-irrgated | | |
| A Absolute abundance (log10 CFU per mL wastewater or g soil) | Fecal indicator bacteria | *E. coli* | n.g. | n.g. | n.g. | n.g. | n.g. | n.g. | n.g. |
| | | 3GCR *E. coli* | n.g. | n.g. | n.g. | n.g. | n.g. | n.g. | n.g. |
| | | Enterococci | n.g. | n.g. | n.g. | n.g. | n.g. | n.g. | n.g. |
| | | VRE | n.g. | n.g. | n.g. | n.g. | n.g. | n.g. | n.g. |
| | Bacteria cultivated on MH at 37°C | Total | 5.85 | **5.27*** | **6.65*** | **5.56*** | **7.05*** | 6.81 | **6.35*** |
| | | CIP resistant | 4.33 | 4.09 | 4.18 | **3.37*** | 5.69 | 5.58 | 5.71 |
| | | TRIM/SUL resistant | 5.28 | **3.74*** | 5.26 | **5.09*** | 5.44 | 5.32 | 5.32 |
| | | ERY/CLIN resistant | 5.37 | **3.90*** | **6.53*** | **4.95*** | 5.47 | 4.87 | 4.90 |
| | | BAC-C12 tolerant | 3.99 | **3.26*** | **4.30*** | **3.64*** | 3.48 | 3.71 | 3.95 |
| | Bacteria cultivated on R2A at 25°C | Total | 6.91 | **6.05*** | 6.91 | 6.19 | 6.64 | 6.35 | 6.46 |
| | | CIP resistant | 5.68 | **4.16*** | 5.87 | **5.57*** | 6.01 | 5.69 | 5.44 |
| | | TRIM/SUL resistant | 5.62 | **4.41*** | **6.08*** | **5.33*** | 5.02 | 5.05 | 4.88 |
| | | ERY/CLIN resistant | 6.26 | **5.04*** | **6.69*** | **5.80*** | 6.06 | 5.54 | 5.65 |
| | | BAC-C12 tolerant | 3.64 | **3.29*** | **4.73*** | **4.00*** | 3.56 | 3.72 | 4.04 |
| B Relative abundance (log10 units) | Bacteria cultivated on MH at 37°C | CIP resistant | 0.48 | 0.82 | **-0.46*** | **-0.18*** | 0.65 | 0.77 | 1.36 |
| | | TRIM/SUL resistant | 1.43 | **0.48*** | **0.61*** | **1.53*** | 0.40 | 0.50 | 0.97 |
| | | ERY/CLIN resistant | 1.52 | **0.63*** | **1.88*** | **1.39*** | 0.42 | 0.06 | 0.55 |
| | | BAC-C12 tolerant | 0.14 | -0.01 | **-0.35*** | 0.09 | -1.57 | -1.08 | -0.40 |
| | Bacteria cultivated on R2A at 25°C | CIP resistant | 0.77 | **0.11*** | 0.96 | **1.38*** | 1.36 | 1.34 | 0.98 |
| | | TRIM/SUL resistant | 0.72 | **0.36*** | **1.18*** | **1.14*** | 0.38 | 0.70 | 0.42 |
| | | ERY/CLIN resistant | 1.35 | **0.99*** | **1.79*** | **1.60*** | 1.41 | 1.19 | 1.19 |
| | | BAC-C12 tolerant | -1.26 | **-0.75*** | **-0.17*** | **-0.19*** | -1.08 | -0.63 | -0.43 |
| Bold font/Asterick - significant differences | | | | to in-fluent | to unspiked Wastewater | | among soil types | | |

| | | | Soil - four weeks after irrigation | | | | | | | | | | | |
|---|---|---|---|---|---|---|---|---|---|---|---|---|---|---|
| | | | Leptosol | | | | Phaeozem | | | | Vertisol | | | |
| | | | Unspiked | | Spiked | | Unspiked | | Spiked | | Unspiked | | Spiked | |
| | | | Influent | Effluent | Influent | Effluent | Influent | Effluent | Influent | Effluent | Influent | Effluent | Influent | Effluent |
| A Absolute abundance (log10 CFU per mL wastewater or g soil) | Fecal indicator bacteria | *E. coli* | n.g. | n.g. | n.g. | n.g. | n.g. | n.g. | n.g. | n.g. | n.g. | n.g. | n.g. | n.g. |
| | | 3GCR *E. coli* | n.g. | n.g. | n.g. | n.g. | n.g. | n.g. | n.g. | n.g. | n.g. | n.g. | n.g. | n.g. |
| | | Enterococci | n.g. | n.g. | n.g. | n.g. | n.g. | n.g. | n.g. | n.g. | n.g. | n.g. | n.g. | n.g. |
| | | VRE | n.g. | n.g. | n.g. | n.g. | n.g. | n.g. | n.g. | n.g. | n.g. | n.g. | n.g. | n.g. |
| | Bacteria cultivated on MH at 37°C | Total | 6.57 | 6.59 | 6.67 | 6.62 | 6.54 | 6.06 | 6.37 | 6.32 | 6.72 | 6.72 | 6.43 | 6.78 |
| | | CIP resistant | 5.13 | 5.08 | 5.04 | 5.18 | 4.80 | 5.11 | 5.11 | 5.45 | 5.03 | 5.11 | 5.17 | 5.08 |
| | | TRIM/SUL resistant | 5.95 | 5.58 | 5.73 | 6.07 | 5.63 | 5.45 | 5.62 | 5.60 | 5.52 | 5.78 | 5.87 | 5.78 |
| | | ERY/CLIN resistant | 5.61 | 5.65 | 5.29 | 5.47 | 5.20 | 5.12 | 5.02 | 5.17 | 5.51 | 5.16 | 5.79 | 5.64 |
| | | BAC-C12 tolerant | 4.24 | 4.19 | 4.47 | 4.30 | 4.22 | 4.31 | 4.31 | 4.31 | 4.68 | 4.69 | 4.74 | 4.48 |
| | Bacteria cultivated on R2A at 25°C | Total | 6.74 | 6.66 | 6.63 | 6.53 | 6.42 | 6.14 | 6.21 | 6.29 | 6.26 | 6.49 | 6.28 | 6.35 |
| | | CIP resistant | 5.81 | 6.04 | 5.95 | 5.85 | 5.62 | 5.53 | 5.58 | 5.57 | 5.94 | 5.80 | 5.94 | 5.47 |
| | | TRIM/SUL resistant | 5.11 | 5.10 | 5.05 | 5.03 | 4.66 | 4.66 | 4.70 | 4.73 | 5.00 | 5.31 | 4.67 | 5.48 |
| | | ERY/CLIN resistant | 6.02 | 5.78 | 5.63 | 5.81 | 5.47 | 5.24 | 5.47 | 5.45 | 5.68 | 5.75 | 5.71 | 5.85 |
| | | BAC-C12 tolerant | 3.99 | 3.96 | 4.15 | 4.00 | 3.98 | 3.96 | 3.97 | 4.01 | 4.30 | 4.85 | 4.10 | 4.39 |
| B Relative abundance (log10 units) | Bacteria cultivated on MH at 37°C | CIP resistant | 0.56 | 0.48 | 0.37 | 0.56 | 0.26 | 1.05 | 0.74 | 1.13 | 0.31 | 0.39 | 0.74 | 0.30 |
| | | TRIM/SUL resistant | **1.38*** | 0.98 | **1.06*** | **1.45*** | **1.09*** | **1.39*** | **1.26*** | **1.27*** | 0.79 | 1.06 | 1.44 | 1.00 |
| | | ERY/CLIN resistant | 1.04 | 1.06 | 0.62 | 0.85 | **0.66*** | **1.06*** | **0.65*** | **0.84*** | 0.79 | 0.44 | 1.36 | 0.87 |
| | | BAC-C12 tolerant | **-0.33*** | **-0.40*** | **-0.20*** | **-0.32*** | -0.32 | **0.26*** | -0.05 | -0.01 | -0.05 | -0.03 | 0.31 | -0.30 |
| | Bacteria cultivated on R2A at 25°C | CIP resistant | 1.07 | 1.37 | 1.33 | 1.32 | 1.21 | 1.39 | 1.36 | 1.28 | 1.68 | 1.31 | 1.66 | 1.12 |
| | | TRIM/SUL resistant | 0.37 | 0.44 | 0.42 | 0.50 | 0.24 | 0.52 | 0.48 | 0.44 | 0.74 | 0.83 | 0.39 | **1.12*** |
| | | ERY/CLIN resistant | 1.28 | 1.12 | 1.00 | 1.28 | 1.05 | 1.10 | 1.25 | 1.16 | 1.42 | 1.26 | 1.44 | 1.50 |
| | | BAC-C12 tolerant | -0.75 | -0.71 | -0.47 | -0.53 | -0.44 | -0.18 | -0.24 | -0.28 | 0.03 | 0.36 | -0.17 | 0.04 |
| Bold font/Asterick - significant differences | | | to non irrigated soil (0 days) | | | | to non irrigated soil (0 days) | | | | to non irrigated soil (0 days) | | | |

**Figure S3.** Heat map showing mean **(A)** absolute abundance (Log10 CFU $mL^{-1}$ or $g^{-1}$) of total and antimicrobial (CIP, TRIM/SUL, ERY/CLIN, and BAC-C12) resistant bacteria cultivable on MH and R2A and **(B)** their relative abundances in WWTP influent and effluent and soil before (0 days) and after irrigation with both unspiked and spiked influent or effluent. Significant differences ($p$<0.05) are indicated with bold font and asterisks (*). No growth: n.g. Mean values of four technical replications (irrigation water types) or four biological replications (four fields per soil type) are presented. Each biological replicate based on four technical replications. For details see Figure S2.

A

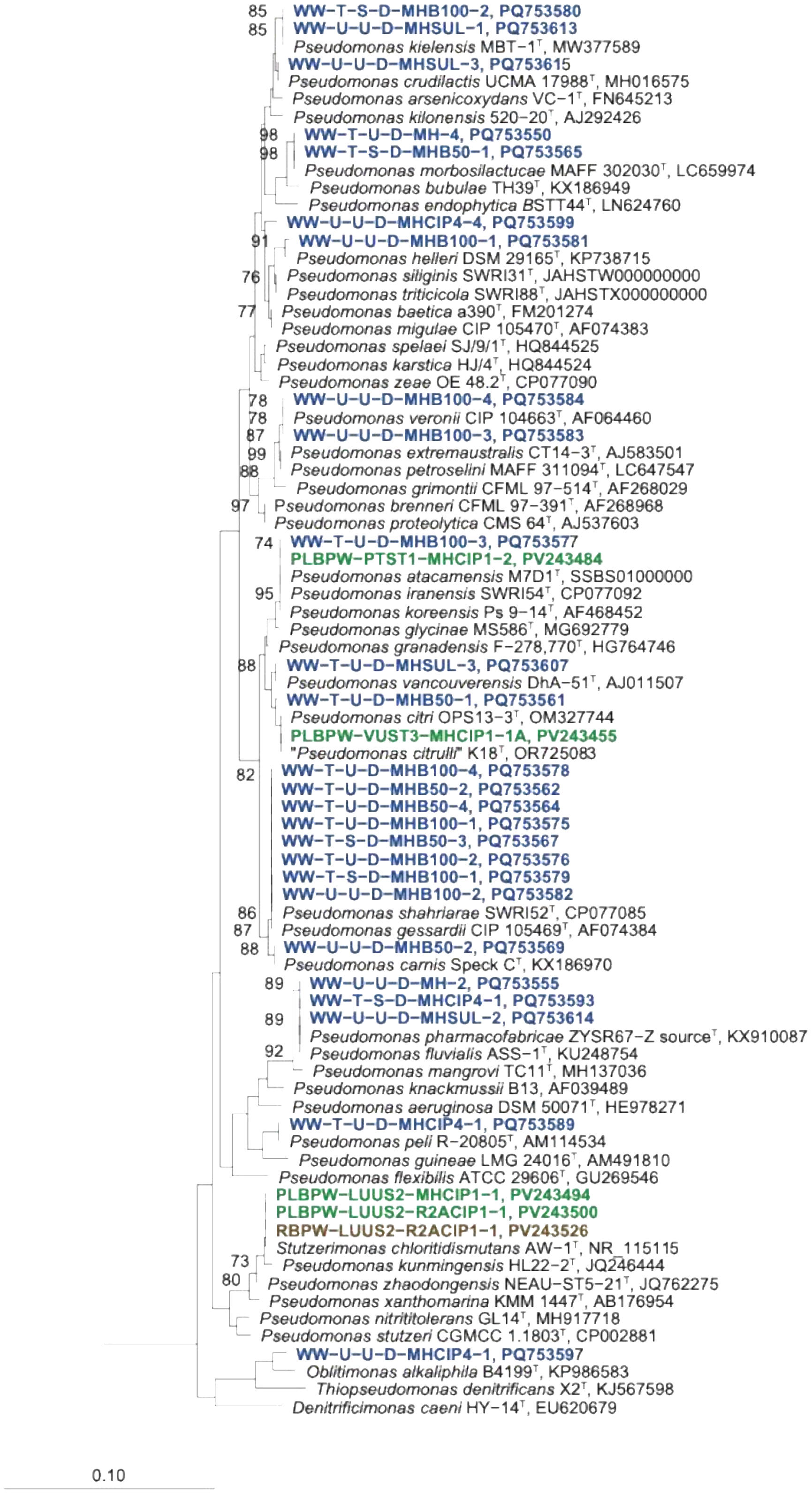

85 WW-T-S-D-MHB100-2, PQ753580
85 WW-U-U-D-MHSUL-1, PQ753613
Pseudomonas kielensis MBT-1$^{T}$, MW377589
WW-U-U-D-MHSUL-3, PQ753615
Pseudomonas crudilactis UCMA 17988$^{T}$, MH016575
Pseudomonas arsenicoxydans VC-1$^{T}$, FN645213
Pseudomonas kilonensis 520-20$^{T}$, AJ292426
98 WW-T-U-D-MH-4, PQ753550
98 WW-T-S-D-MHB50-1, PQ753565
Pseudomonas morbosilactucae MAFF 302030$^{T}$, LC659974
Pseudomonas bubulae TH39$^{T}$, KX186949
Pseudomonas endophytica BSTT44$^{T}$, LN624760
WW-U-U-D-MHCIP4-4, PQ753599
91 WW-U-U-D-MHB100-1, PQ753581
Pseudomonas helleri DSM 29165$^{T}$, KP738715
76 Pseudomonas siliginis SWRI31$^{T}$, JAHSTW000000000
Pseudomonas triticicola SWRI88$^{T}$, JAHSTX000000000
77 Pseudomonas baetica a390$^{T}$, FM201274
Pseudomonas migulae CIP 105470$^{T}$, AF074383
Pseudomonas spelaei SJ/9/1$^{T}$, HQ844525
Pseudomonas karstica HJ/4$^{T}$, HQ844524
Pseudomonas zeae OE 48.2$^{T}$, CP077090
78 WW-U-U-D-MHB100-4, PQ753584
78 Pseudomonas veronii CIP 104663$^{T}$, AF064460
87 WW-U-U-D-MHB100-3, PQ753583
99 Pseudomonas extremaustralis CT14-3$^{T}$, AJ583501
88 Pseudomonas petroselini MAFF 311094$^{T}$, LC647547
Pseudomonas grimontii CFML 97-514$^{T}$, AF268029
97 Pseudomonas brenneri CFML 97-391$^{T}$, AF268968
Pseudomonas proteolytica CMS 64$^{T}$, AJ537603
74 WW-T-U-D-MHB100-3, PQ753577
PLBPW-PTST1-MHCIP1-2, PV243484
Pseudomonas atacamensis M7D1$^{T}$, SSBS01000000
95 Pseudomonas iranensis SWRI54$^{T}$, CP077092
Pseudomonas koreensis Ps 9-14$^{T}$, AF468452
Pseudomonas glycinae MS586$^{T}$, MG692779
Pseudomonas granadensis F-278,770$^{T}$, HG764746
88 WW-T-U-D-MHSUL-3, PQ753607
Pseudomonas vancouverensis DhA-51$^{T}$, AJ011507
WW-T-U-D-MHB50-1, PQ753561
Pseudomonas citri OPS13-3$^{T}$, OM327744
PLBPW-VUST3-MHCIP1-1A, PV243455
"Pseudomonas citrulli" K18$^{T}$, OR725083
82 WW-T-U-D-MHB100-4, PQ753578
WW-T-U-D-MHB50-2, PQ753562
WW-T-U-D-MHB50-4, PQ753564
WW-T-U-D-MHB100-1, PQ753575
WW-T-S-D-MHB50-3, PQ753567
WW-T-U-D-MHB100-2, PQ753576
WW-T-S-D-MHB100-1, PQ753579
WW-U-U-D-MHB100-2, PQ753582
86 Pseudomonas shahriarae SWRI52$^{T}$, CP077085
87 Pseudomonas gessardii CIP 105469$^{T}$, AF074384
88 WW-U-U-D-MHB50-2, PQ753569
Pseudomonas carnis Speck C$^{T}$, KX186970
89 WW-U-U-D-MH-2, PQ753555
WW-T-S-D-MHCIP4-1, PQ753593
89 WW-U-U-D-MHSUL-2, PQ753614
Pseudomonas pharmacofabricae ZYSR67-Z source$^{T}$, KX910087
92 Pseudomonas fluvialis ASS-1$^{T}$, KU248754
Pseudomonas mangrovi TC11$^{T}$, MH137036
Pseudomonas knackmussii B13, AF039489
Pseudomonas aeruginosa DSM 50071$^{T}$, HE978271
WW-T-U-D-MHCIP4-1, PQ753589
Pseudomonas peli R-20805$^{T}$, AM114534
Pseudomonas guineae LMG 24016$^{T}$, AM491810
Pseudomonas flexibilis ATCC 29606$^{T}$, GU269546
PLBPW-LUUS2-MHCIP1-1, PV243494
PLBPW-LUUS2-R2ACIP1-1, PV243500
RBPW-LUUS2-R2ACIP1-1, PV243526
Stutzerimonas chloritidismutans AW-1$^{T}$, NR_115115
73 Pseudomonas kunmingensis HL22-2$^{T}$, JQ246444
80 Pseudomonas zhaodongensis NEAU-ST5-21$^{T}$, JQ762275
Pseudomonas xanthomarina KMM 1447$^{T}$, AB176954
Pseudomonas nitritireducens GL14$^{T}$, MH917718
Pseudomonas stutzeri CGMCC 1.1803$^{T}$, CP002881
WW-U-U-D-MHCIP4-1, PQ753597
Oblitimonas alkaliphila B4199$^{T}$, KP986583
Thiopseudomonas denitrificans X2$^{T}$, KJ567598
Denitrificimonas caeni HY-14$^{T}$, EU620679
0.10

**B**

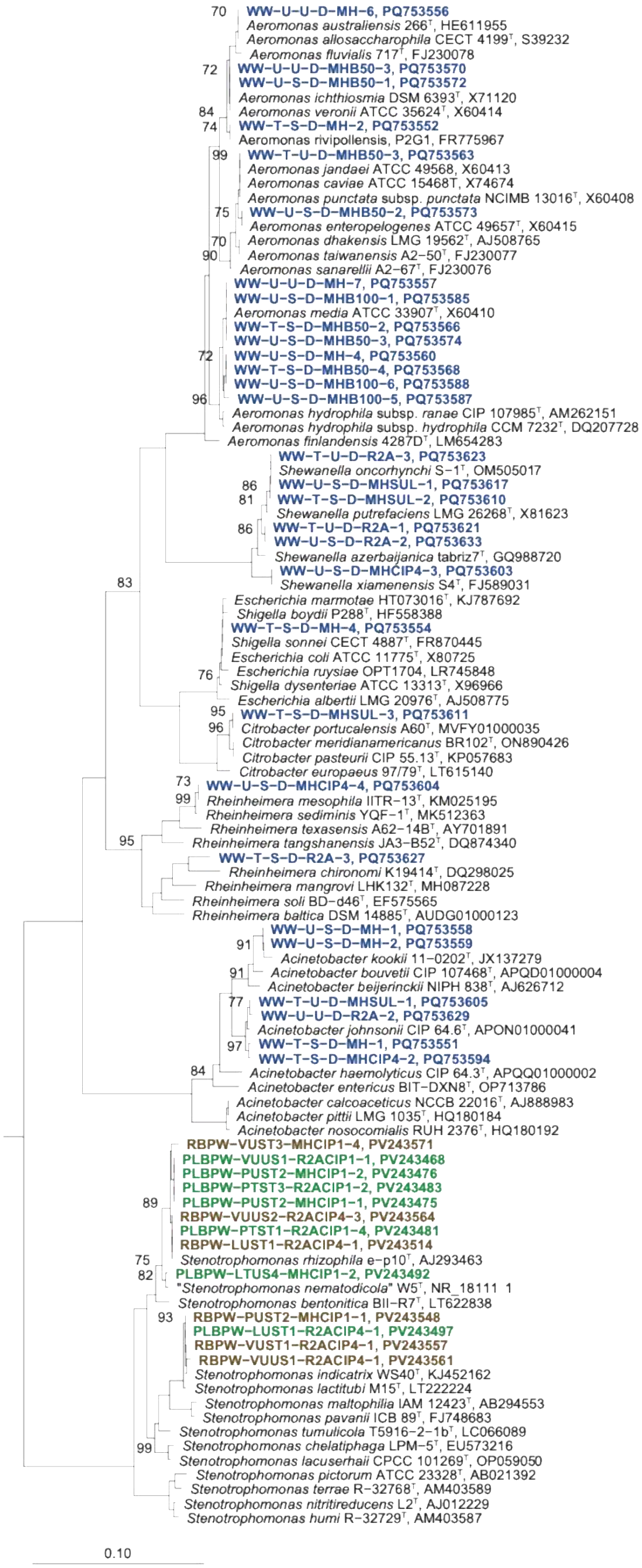

WW-U-U-D-MH-6, PQ753556
Aeromonas australiensis 266T, HE611955
Aeromonas allosaccharophila CECT 4199T, S39232
Aeromonas fluvialis 717T, FJ230078
WW-U-U-D-MHB50-3, PQ753570
WW-U-S-D-MHB50-1, PQ753572
Aeromonas ichthiosmia DSM 6393T, X71120
Aeromonas veronii ATCC 35624T, X60414
WW-T-S-D-MH-2, PQ753552
Aeromonas rivipollensis, P2G1, FR775967
WW-T-U-D-MHB50-3, PQ753563
Aeromonas jandaei ATCC 49568, X60413
Aeromonas caviae ATCC 15468T, X74674
Aeromonas punctata subsp. punctata NCIMB 13016T, X60408
WW-U-S-D-MHB50-2, PQ753573
Aeromonas enteropelogenes ATCC 49657T, X60415
Aeromonas dhakensis LMG 19562T, AJ508765
Aeromonas taiwanensis A2-50T, FJ230077
Aeromonas sanarellii A2-67T, FJ230076
WW-U-U-D-MH-7, PQ753557
WW-U-S-D-MHB100-1, PQ753585
Aeromonas media ATCC 33907T, X60410
WW-T-S-D-MHB50-2, PQ753566
WW-U-S-D-MHB50-3, PQ753574
WW-U-S-D-MH-4, PQ753560
WW-T-S-D-MHB50-4, PQ753568
WW-U-S-D-MHB100-6, PQ753588
WW-U-S-D-MHB100-5, PQ753587
Aeromonas hydrophila subsp. ranae CIP 107985T, AM262151
Aeromonas hydrophila subsp. hydrophila CCM 7232T, DQ207728
Aeromonas finlandensis 4287DT, LM654283
WW-T-U-D-R2A-3, PQ753623
Shewanella oncorhynchi S-1T, OM505017
WW-U-S-D-MHSUL-1, PQ753617
WW-T-S-D-MHSUL-2, PQ753610
Shewanella putrefaciens LMG 26268T, X81623
WW-T-U-D-R2A-1, PQ753621
WW-U-S-D-R2A-2, PQ753633
Shewanella azerbaijanica tabriz7T, GQ988720
WW-U-S-D-MHCIP4-3, PQ753603
Shewanella xiamenensis S4T, FJ589031
Escherichia marmotae HT073016T, KJ787692
Shigella boydii P288T, HF558388
WW-T-S-D-MH-4, PQ753554
Shigella sonnei CECT 4887T, FR870445
Escherichia coli ATCC 11775T, X80725
Escherichia ruysiae OPT1704, LR745848
Shigella dysenteriae ATCC 13313T, X96966
Escherichia albertii LMG 20976T, AJ508775
WW-T-S-D-MHSUL-3, PQ753611
Citrobacter portucalensis A60T, MVFY01000035
Citrobacter meridianamericanus BR102T, ON890426
Citrobacter pasteurii CIP 55.13T, KP057683
Citrobacter europaeus 97/79T, LT615140
WW-U-S-D-MHCIP4-4, PQ753604
Rheinheimera mesophila IITR-13T, KM025195
Rheinheimera sediminis YQF-1T, MK512363
Rheinheimera texasensis A62-14BT, AY701891
Rheinheimera tangshanensis JA3-B52T, DQ874340
WW-T-S-D-R2A-3, PQ753627
Rheinheimera chironomi K19414T, DQ298025
Rheinheimera mangrovi LHK132T, MH087228
Rheinheimera soli BD-d46T, EF575565
Rheinheimera baltica DSM 14885T, AUDG01000123
WW-U-S-D-MH-1, PQ753558
WW-U-S-D-MH-2, PQ753559
Acinetobacter kookii 11-0202T, JX137279
Acinetobacter bouvetii CIP 107468T, APQD01000004
Acinetobacter beijerinckii NIPH 838T, AJ626712
WW-T-U-D-MHSUL-1, PQ753605
WW-U-U-D-R2A-2, PQ753629
Acinetobacter johnsonii CIP 64.6T, APON01000041
WW-T-S-D-MH-1, PQ753551
WW-T-S-D-MHCIP4-2, PQ753594
Acinetobacter haemolyticus CIP 64.3T, APQQ01000002
Acinetobacter entericus BIT-DXN8T, OP713786
Acinetobacter calcoaceticus NCCB 22016T, AJ888983
Acinetobacter pittii LMG 1035T, HQ180184
Acinetobacter nosocomialis RUH 2376T, HQ180192
RBPW-VUST3-MHCIP1-4, PV243571
PLBPW-VUUS1-R2ACIP1-1, PV243468
PLBPW-PUST2-MHCIP1-2, PV243476
PLBPW-PTST3-R2ACIP1-2, PV243483
PLBPW-PUST2-MHCIP1-1, PV243475
RBPW-VUUS2-R2ACIP4-3, PV243564
PLBPW-PTST1-R2ACIP1-4, PV243481
RBPW-LUST1-R2ACIP4-1, PV243514
Stenotrophomonas rhizophila e-p10T, AJ293463
PLBPW-LTUS4-MHCIP1-2, PV243492
"Stenotrophomonas nematodicola" W5T, NR_18111 1
Stenotrophomonas bentonitica BII-R7T, LT622838
RBPW-PUST2-MHCIP1-1, PV243548
PLBPW-LUST1-R2ACIP4-1, PV243497
RBPW-VUST1-R2ACIP4-1, PV243557
RBPW-VUUS1-R2ACIP4-1, PV243561
Stenotrophomonas indicatrix WS40T, KJ452162
Stenotrophomonas lactitubi M15T, LT222224
Stenotrophomonas maltophilia IAM 12423T, AB294553
Stenotrophomonas pavanii ICB 89T, FJ748683
Stenotrophomonas tumulicola T5916-2-1bT, LC066089
Stenotrophomonas chelatiphaga LPM-5T, EU573216
Stenotrophomonas lacuserhaii CPCC 101269T, OP059050
Stenotrophomonas pictorum ATCC 23328T, AB021392
Stenotrophomonas terrae R-32768T, AM403589
Stenotrophomonas nitritireducens L2T, AJ012229
Stenotrophomonas humi R-32729T, AM403587
0.10

C

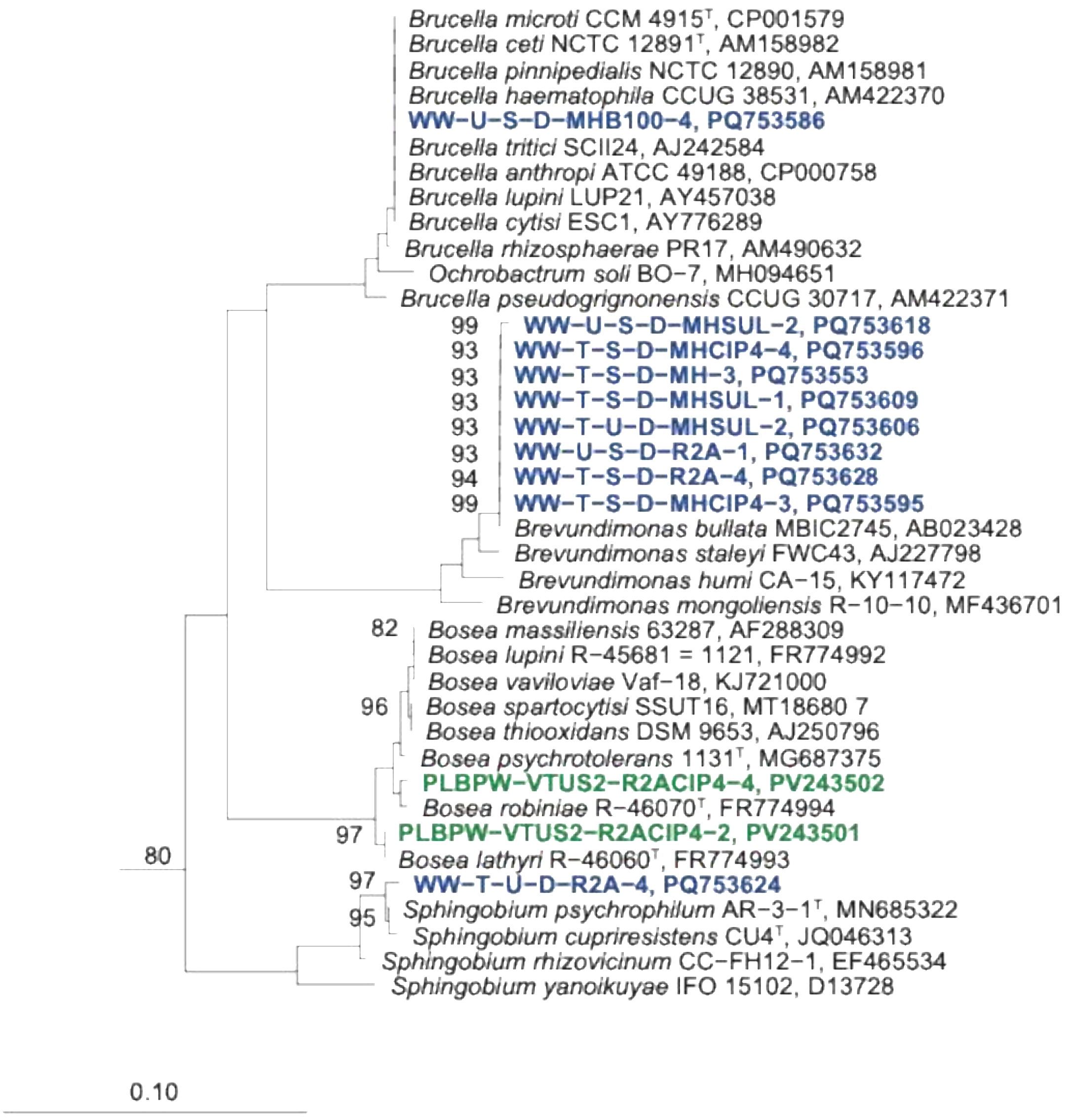
Brucella microti CCM 4915T, CP001579
Brucella ceti NCTC 12891T, AM158982
Brucella pinnipedialis NCTC 12890, AM158981
Brucella haematophila CCUG 38531, AM422370
WW-U-S-D-MHB100-4, PQ753586
Brucella tritici SCII24, AJ242584
Brucella anthropi ATCC 49188, CP000758
Brucella lupini LUP21, AY457038
Brucella cytisi ESC1, AY776289
Brucella rhizosphaerae PR17, AM490632
Ochrobactrum soli BO-7, MH094651
Brucella pseudogrignonensis CCUG 30717, AM422371
99
WW-U-S-D-MHSUL-2, PQ753618
93
WW-T-S-D-MHCIP4-4, PQ753596
93
WW-T-S-D-MH-3, PQ753553
93
WW-T-S-D-MHSUL-1, PQ753609
93
WW-T-U-D-MHSUL-2, PQ753606
93
WW-U-S-D-R2A-1, PQ753632
94
WW-T-S-D-R2A-4, PQ753628
99
WW-T-S-D-MHCIP4-3, PQ753595
Brevundimonas bullata MBIC2745, AB023428
Brevundimonas staleyi FWC43, AJ227798
Brevundimonas humi CA-15, KY117472
Brevundimonas mongoliensis R-10-10, MF436701
82
Bosea massiliensis 63287, AF288309
Bosea lupini R-45681 = 1121, FR774992
Bosea vaviloviae Vaf-18, KJ721000
96
Bosea spartocytisi SSUT16, MT18680 7
Bosea thiooxidans DSM 9653, AJ250796
Bosea psychrotolerans 1131T, MG687375
PLBPW-VTUS2-R2ACIP4-4, PV243502
Bosea robiniae R-46070T, FR774994
97
PLBPW-VTUS2-R2ACIP4-2, PV243501
Bosea lathyri R-46060T, FR774993
80
97
WW-T-U-D-R2A-4, PQ753624
95
Sphingobium psychrophilum AR-3-1T, MN685322
Sphingobium cupriresistens CU4T, JQ046313
Sphingobium rhizovicinum CC-FH12-1, EF465534
Sphingobium yanoikuyae IFO 15102, D13728
0.10

**D**

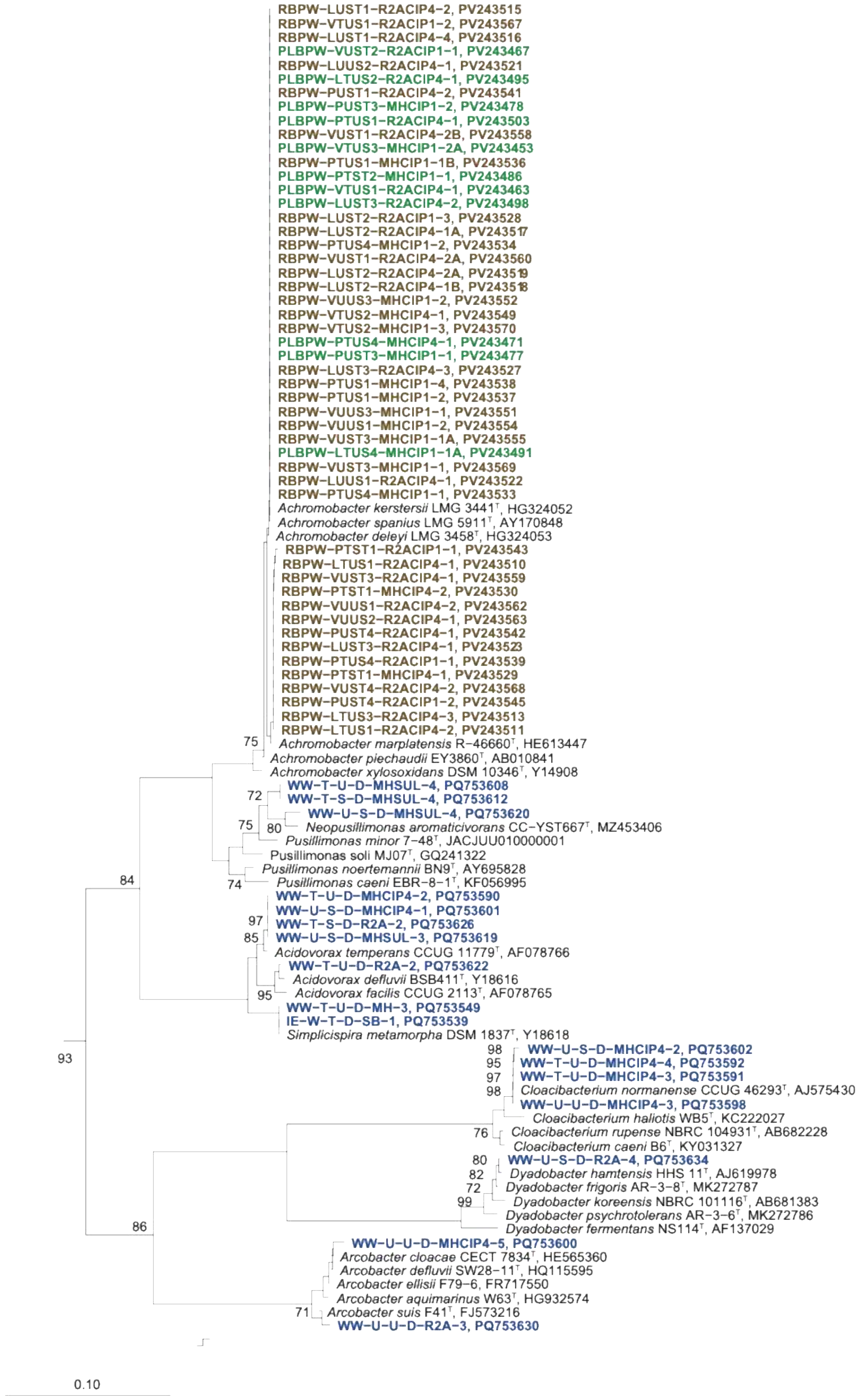

RBPW-LUST1-R2ACIP4-2, PV243515
RBPW-VTUS1-R2ACIP1-2, PV243567
RBPW-LUST1-R2ACIP4-4, PV243516
PLBPW-VUST2-R2ACIP1-1, PV243467
RBPW-LUUS2-R2ACIP4-1, PV243521
PLBPW-LTUS2-R2ACIP4-1, PV243495
RBPW-PUST1-R2ACIP4-2, PV243541
PLBPW-PUST3-MHCIP1-2, PV243478
PLBPW-PTUS1-R2ACIP4-1, PV243503
RBPW-VUST1-R2ACIP4-2B, PV243558
PLBPW-VTUS3-MHCIP1-2A, PV243453
RBPW-PTUS1-MHCIP1-1B, PV243536
PLBPW-PTST2-MHCIP1-1, PV243486
PLBPW-VTUS1-R2ACIP4-1, PV243463
PLBPW-LUST3-R2ACIP4-2, PV243498
RBPW-LUST2-R2ACIP1-3, PV243528
RBPW-LUST2-R2ACIP4-1A, PV243517
RBPW-PTUS4-MHCIP1-2, PV243534
RBPW-VUST1-R2ACIP4-2A, PV243560
RBPW-LUST2-R2ACIP4-2A, PV243519
RBPW-LUST2-R2ACIP4-1B, PV243518
RBPW-VUUS3-MHCIP1-2, PV243552
RBPW-VTUS2-MHCIP4-1, PV243549
RBPW-VTUS2-MHCIP1-3, PV243570
PLBPW-PTUS4-MHCIP4-1, PV243471
PLBPW-PUST3-MHCIP1-1, PV243477
RBPW-LUST3-R2ACIP4-3, PV243527
RBPW-PTUS1-MHCIP1-4, PV243538
RBPW-PTUS1-MHCIP1-2, PV243537
RBPW-VUUS3-MHCIP1-1, PV243551
RBPW-VUUS1-MHCIP1-2, PV243554
RBPW-VUST3-MHCIP1-1A, PV243555
PLBPW-LTUS4-MHCIP1-1A, PV243491
RBPW-VUST3-MHCIP1-1, PV243569
RBPW-LUUS1-R2ACIP4-1, PV243522
RBPW-PTUS4-MHCIP1-1, PV243533
Achromobacter kerstersii LMG 3441$^{T}$, HG324052
Achromobacter spanius LMG 5911$^{T}$, AY170848
Achromobacter deleyi LMG 3458$^{T}$, HG324053
RBPW-PTST1-R2ACIP1-1, PV243543
RBPW-LTUS1-R2ACIP4-1, PV243510
RBPW-VUST3-R2ACIP4-1, PV243559
RBPW-PTST1-MHCIP4-2, PV243530
RBPW-VUUS1-R2ACIP4-2, PV243562
RBPW-VUUS2-R2ACIP4-1, PV243563
RBPW-PUST4-R2ACIP4-1, PV243542
RBPW-LUST3-R2ACIP4-1, PV243523
RBPW-PTUS4-R2ACIP1-1, PV243539
RBPW-PTST1-MHCIP4-1, PV243529
RBPW-VUST4-R2ACIP4-2, PV243568
RBPW-PUST4-R2ACIP1-2, PV243545
RBPW-LTUS3-R2ACIP4-3, PV243513
RBPW-LTUS1-R2ACIP4-2, PV243511
75
Achromobacter marplatensis R-46660$^{T}$, HE613447
Achromobacter piechaudii EY3860$^{T}$, AB010841
Achromobacter xylosoxidans DSM 10346$^{T}$, Y14908
WW-T-U-D-MHSUL-4, PQ753608
72
WW-T-S-D-MHSUL-4, PQ753612
WW-U-S-D-MHSUL-4, PQ753620
75
80
Neopusillimonas aromaticivorans CC-YST667$^{T}$, MZ453406
Pusillimonas minor 7-48$^{T}$, JACJUU010000001
Pusillimonas soli MJ07$^{T}$, GQ241322
84
Pusillimonas noertemannii BN9$^{T}$, AY695828
74
Pusillimonas caeni EBR-8-1$^{T}$, KF056995
WW-T-U-D-MHCIP4-2, PQ753590
WW-U-S-D-MHCIP4-1, PQ753601
97
WW-T-S-D-R2A-2, PQ753626
85
WW-U-S-D-MHSUL-3, PQ753619
Acidovorax temperans CCUG 11779$^{T}$, AF078766
WW-T-U-D-R2A-2, PQ753622
Acidovorax defluvii BSB411$^{T}$, Y18616
95
Acidovorax facilis CCUG 2113$^{T}$, AF078765
WW-T-U-D-MH-3, PQ753549
IE-W-T-D-SB-1, PQ753539
Simplicispira metamorpha DSM 1837$^{T}$, Y18618
93
98
WW-U-S-D-MHCIP4-2, PQ753602
95
WW-T-U-D-MHCIP4-4, PQ753592
97
WW-T-U-D-MHCIP4-3, PQ753591
98
Cloacibacterium normanense CCUG 46293$^{T}$, AJ575430
WW-U-U-D-MHCIP4-3, PQ753598
Cloacibacterium haliotis WB5$^{T}$, KC222027
76
Cloacibacterium rupense NBRC 104931$^{T}$, AB682228
Cloacibacterium caeni B6$^{T}$, KY031327
80
WW-U-S-D-R2A-4, PQ753634
82
Dyadobacter hamtensis HHS 11$^{T}$, AJ619978
72
Dyadobacter frigoris AR-3-8$^{T}$, MK272787
99
Dyadobacter koreensis NBRC 101116$^{T}$, AB681383
Dyadobacter psychrotolerans AR-3-6$^{T}$, MK272786
86
Dyadobacter fermentans NS114$^{T}$, AF137029
WW-U-U-D-MHCIP4-5, PQ753600
Arcobacter cloacae CECT 7834$^{T}$, HE565360
Arcobacter defluvii SW28-11$^{T}$, HQ115595
Arcobacter ellisii F79-6, FR717550
Arcobacter aquimarinus W63$^{T}$, HG932574
71
Arcobacter suis F41$^{T}$, FJ573216
WW-U-U-D-R2A-3, PQ753630
0.10

**E**

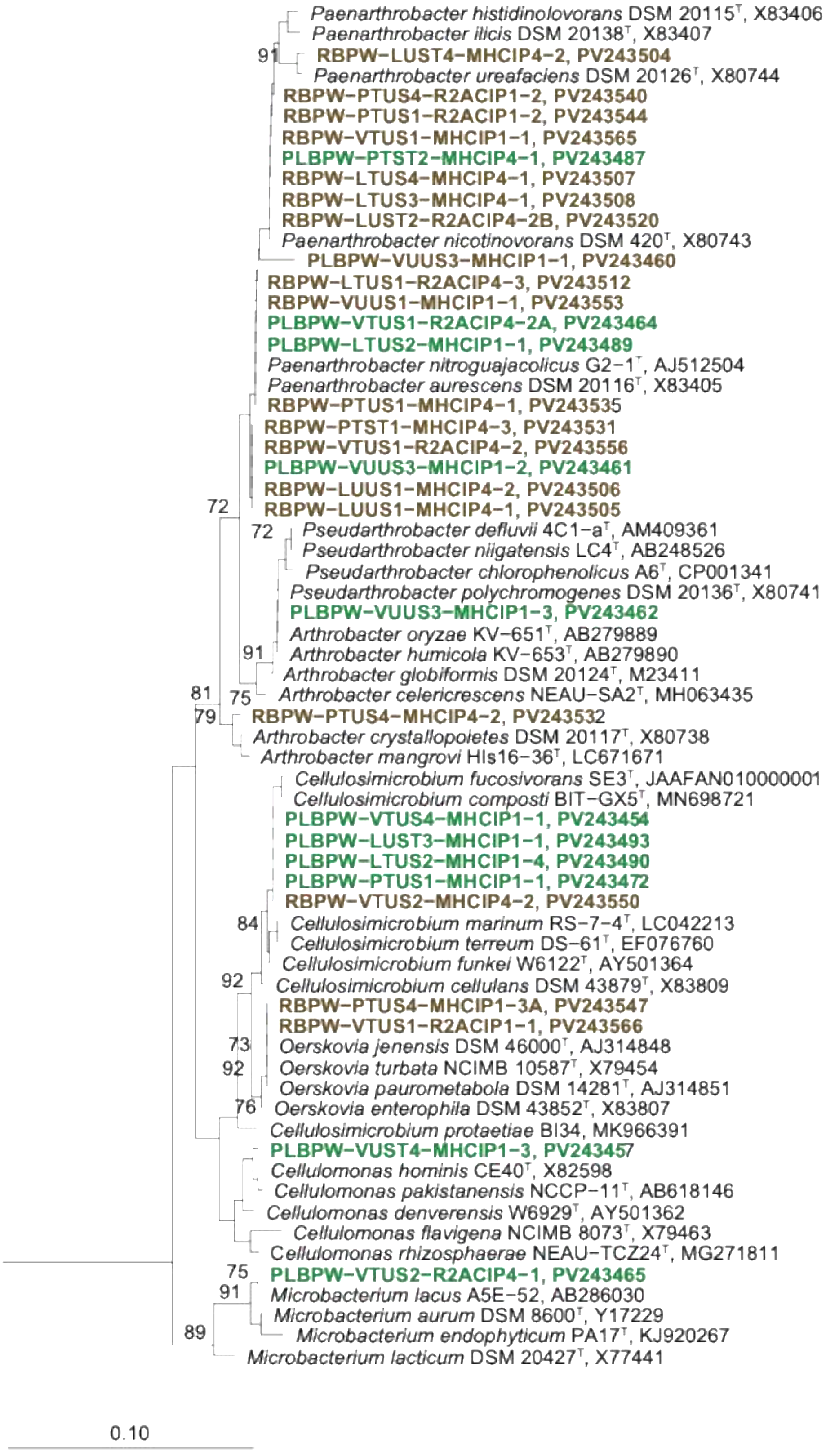
Paenarthrobacter histidinolovorans DSM 20115$^{T}$, X83406
Paenarthrobacter ilicis DSM 20138$^{T}$, X83407
91
RBPW-LUST4-MHCIP4-2, PV243504
Paenarthrobacter ureafaciens DSM 20126$^{T}$, X80744
RBPW-PTUS4-R2ACIP1-2, PV243540
RBPW-PTUS1-R2ACIP1-2, PV243544
RBPW-VTUS1-MHCIP1-1, PV243565
PLBPW-PTST2-MHCIP4-1, PV243487
RBPW-LTUS4-MHCIP4-1, PV243507
RBPW-LTUS3-MHCIP4-1, PV243508
RBPW-LUST2-R2ACIP4-2B, PV243520
Paenarthrobacter nicotinovorans DSM 420$^{T}$, X80743
PLBPW-VUUS3-MHCIP1-1, PV243460
RBPW-LTUS1-R2ACIP4-3, PV243512
RBPW-VUUS1-MHCIP1-1, PV243553
PLBPW-VTUS1-R2ACIP4-2A, PV243464
PLBPW-LTUS2-MHCIP1-1, PV243489
Paenarthrobacter nitroguajacolicus G2-1$^{T}$, AJ512504
Paenarthrobacter aurescens DSM 20116$^{T}$, X83405
RBPW-PTUS1-MHCIP4-1, PV243535
RBPW-PTST1-MHCIP4-3, PV243531
RBPW-VTUS1-R2ACIP4-2, PV243556
PLBPW-VUUS3-MHCIP1-2, PV243461
RBPW-LUUS1-MHCIP4-2, PV243506
RBPW-LUUS1-MHCIP4-1, PV243505
72
72
Pseudarthrobacter defluvii 4C1-a$^{T}$, AM409361
Pseudarthrobacter niigatensis LC4$^{T}$, AB248526
Pseudarthrobacter chlorophenolicus A6$^{T}$, CP001341
Pseudarthrobacter polychromogenes DSM 20136$^{T}$, X80741
PLBPW-VUUS3-MHCIP1-3, PV243462
Arthrobacter oryzae KV-651$^{T}$, AB279889
91
Arthrobacter humicola KV-653$^{T}$, AB279890
Arthrobacter globiformis DSM 20124$^{T}$, M23411
81
75
Arthrobacter celericrescens NEAU-SA2$^{T}$, MH063435
79
RBPW-PTUS4-MHCIP4-2, PV243532
Arthrobacter crystallopoietes DSM 20117$^{T}$, X80738
Arthrobacter mangrovi HIs16-36$^{T}$, LC671671
Cellulosimicrobium fucosivorans SE3$^{T}$, JAAFAN010000001
Cellulosimicrobium composti BIT-GX5$^{T}$, MN698721
PLBPW-VTUS4-MHCIP1-1, PV243454
PLBPW-LUST3-MHCIP1-1, PV243493
PLBPW-LTUS2-MHCIP1-4, PV243490
PLBPW-PTUS1-MHCIP1-1, PV243472
RBPW-VTUS2-MHCIP4-2, PV243550
84
Cellulosimicrobium marinum RS-7-4$^{T}$, LC042213
Cellulosimicrobium terreum DS-61$^{T}$, EF076760
Cellulosimicrobium funkei W6122$^{T}$, AY501364
92
Cellulosimicrobium cellulans DSM 43879$^{T}$, X83809
RBPW-PTUS4-MHCIP1-3A, PV243547
RBPW-VTUS1-R2ACIP1-1, PV243566
73
Oerskovia jenensis DSM 46000$^{T}$, AJ314848
92
Oerskovia turbata NCIMB 10587$^{T}$, X79454
Oerskovia paurometabola DSM 14281$^{T}$, AJ314851
76
Oerskovia enterophila DSM 43852$^{T}$, X83807
Cellulosimicrobium protaetiae BI34, MK966391
PLBPW-VUST4-MHCIP1-3, PV243457
Cellulomonas hominis CE40$^{T}$, X82598
Cellulomonas pakistanensis NCCP-11$^{T}$, AB618146
Cellulomonas denverensis W6929$^{T}$, AY501362
Cellulomonas flavigena NCIMB 8073$^{T}$, X79463
Cellulomonas rhizosphaerae NEAU-TCZ24$^{T}$, MG271811
75
PLBPW-VTUS2-R2ACIP4-1, PV243465
91
Microbacterium lacus A5E-52, AB286030
Microbacterium aurum DSM 8600$^{T}$, Y17229
89
Microbacterium endophyticum PA17$^{T}$, KJ920267
Microbacterium lacticum DSM 20427$^{T}$, X77441
0.10

F

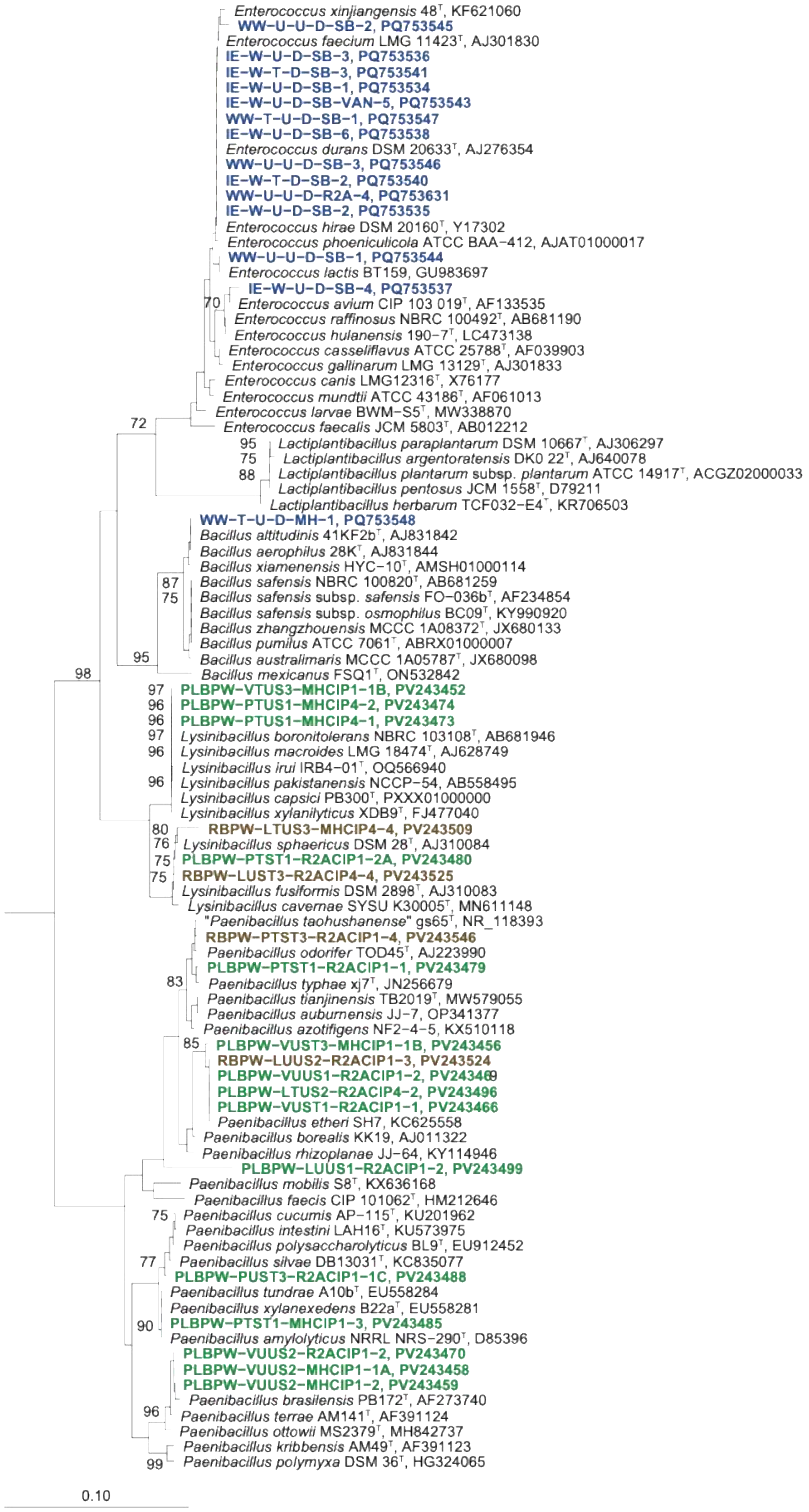
Enterococcus xinjiangensis 48$^{T}$, KF621060
WW-U-U-D-SB-2, PQ753545
Enterococcus faecium LMG 11423$^{T}$, AJ301830
IE-W-U-D-SB-3, PQ753536
IE-W-T-D-SB-3, PQ753541
IE-W-U-D-SB-1, PQ753534
IE-W-U-D-SB-VAN-5, PQ753543
WW-T-U-D-SB-1, PQ753547
IE-W-U-D-SB-6, PQ753538
Enterococcus durans DSM 20633$^{T}$, AJ276354
WW-U-U-D-SB-3, PQ753546
IE-W-T-D-SB-2, PQ753540
WW-U-U-D-R2A-4, PQ753631
IE-W-U-D-SB-2, PQ753535
Enterococcus hirae DSM 20160$^{T}$, Y17302
Enterococcus phoeniculicola ATCC BAA-412, AJAT01000017
WW-U-U-D-SB-1, PQ753544
Enterococcus lactis BT159, GU983697
IE-W-U-D-SB-4, PQ753537
70 Enterococcus avium CIP 103 019$^{T}$, AF133535
Enterococcus raffinosus NBRC 100492$^{T}$, AB681190
Enterococcus hulanensis 190-7$^{T}$, LC473138
Enterococcus casseliflavus ATCC 25788$^{T}$, AF039903
Enterococcus gallinarum LMG 13129$^{T}$, AJ301833
Enterococcus canis LMG12316$^{T}$, X76177
Enterococcus mundtii ATCC 43186$^{T}$, AF061013
Enterococcus larvae BWM-S5$^{T}$, MW338870
72 Enterococcus faecalis JCM 5803$^{T}$, AB012212
95 Lactiplantibacillus paraplantarum DSM 10667$^{T}$, AJ306297
75 Lactiplantibacillus argentoratensis DK0 22$^{T}$, AJ640078
88 Lactiplantibacillus plantarum subsp. plantarum ATCC 14917$^{T}$, ACGZ02000033
Lactiplantibacillus pentosus JCM 1558$^{T}$, D79211
Lactiplantibacillus herbarum TCF032-E4$^{T}$, KR706503
WW-T-U-D-MH-1, PQ753548
Bacillus altitudinis 41KF2b$^{T}$, AJ831842
Bacillus aerophilus 28K$^{T}$, AJ831844
Bacillus xiamenensis HYC-10$^{T}$, AMSH01000114
87 Bacillus safensis NBRC 100820$^{T}$, AB681259
75 Bacillus safensis subsp. safensis FO-036b$^{T}$, AF234854
Bacillus safensis subsp. osmophilus BC09$^{T}$, KY990920
Bacillus zhangzhouensis MCCC 1A08372$^{T}$, JX680133
Bacillus pumilus ATCC 7061$^{T}$, ABRX01000007
95 Bacillus australimaris MCCC 1A05787$^{T}$, JX680098
98 Bacillus mexicanus FSQ1$^{T}$, ON532842
97 PLBPW-VTUS3-MHCIP1-1B, PV243452
96 PLBPW-PTUS1-MHCIP4-2, PV243474
96 PLBPW-PTUS1-MHCIP4-1, PV243473
97 Lysinibacillus boronitolerans NBRC 103108$^{T}$, AB681946
96 Lysinibacillus macroides LMG 18474$^{T}$, AJ628749
Lysinibacillus irui IRB4-01$^{T}$, OQ566940
96 Lysinibacillus pakistanensis NCCP-54, AB558495
Lysinibacillus capsici PB300$^{T}$, PXXX01000000
Lysinibacillus xylanilyticus XDB9$^{T}$, FJ477040
80 RBPW-LTUS3-MHCIP4-4, PV243509
76 Lysinibacillus sphaericus DSM 28$^{T}$, AJ310084
75 PLBPW-PTST1-R2ACIP1-2A, PV243480
75 RBPW-LUST3-R2ACIP4-4, PV243525
Lysinibacillus fusiformis DSM 2898$^{T}$, AJ310083
Lysinibacillus cavernae SYSU K30005$^{T}$, MN611148
"Paenibacillus taohushanense" gs65$^{T}$, NR_118393
RBPW-PTST3-R2ACIP1-4, PV243546
Paenibacillus odorifer TOD45$^{T}$, AJ223990
PLBPW-PTST1-R2ACIP1-1, PV243479
83 Paenibacillus typhae xj7$^{T}$, JN256679
Paenibacillus tianjinensis TB2019$^{T}$, MW579055
Paenibacillus auburnensis JJ-7, OP341377
Paenibacillus azotifigens NF2-4-5, KX510118
85 PLBPW-VUST3-MHCIP1-1B, PV243456
RBPW-LUUS2-R2ACIP1-3, PV243524
PLBPW-VUUS1-R2ACIP1-2, PV243469
PLBPW-LTUS2-R2ACIP4-2, PV243496
PLBPW-VUST1-R2ACIP1-1, PV243466
Paenibacillus etheri SH7, KC625558
Paenibacillus borealis KK19, AJ011322
Paenibacillus rhizoplanae JJ-64, KY114946
PLBPW-LUUS1-R2ACIP1-2, PV243499
Paenibacillus mobilis S8$^{T}$, KX636168
Paenibacillus faecis CIP 101062$^{T}$, HM212646
75 Paenibacillus cucumis AP-115$^{T}$, KU201962
Paenibacillus intestini LAH16$^{T}$, KU573975
Paenibacillus polysaccharolyticus BL9$^{T}$, EU912452
77 Paenibacillus silvae DB13031$^{T}$, KC835077
PLBPW-PUST3-R2ACIP1-1C, PV243488
Paenibacillus tundrae A10b$^{T}$, EU558284
Paenibacillus xylanexedens B22a$^{T}$, EU558281
90 PLBPW-PTST1-MHCIP1-3, PV243485
Paenibacillus amylolyticus NRRL NRS-290$^{T}$, D85396
PLBPW-VUUS2-R2ACIP1-2, PV243470
PLBPW-VUUS2-MHCIP1-1A, PV243458
PLBPW-VUUS2-MHCIP1-2, PV243459
Paenibacillus brasilensis PB172$^{T}$, AF273740
96 Paenibacillus terrae AM141$^{T}$, AF391124
Paenibacillus ottowii MS2379$^{T}$, MH842737
Paenibacillus kribbensis AM49$^{T}$, AF391123
99 Paenibacillus polymyxa DSM 36$^{T}$, HG324065
0.10

**Figure S4.** Phylogenetic placement of cultivated bacterial strains from this study based on partial 16S rRNA gene sequences. The phylogenetic tree was calculated in ARB in the LTP type strain database using the Neighbour joining method. Due to a better overview the calculated tree was spitted in subtrees. A/B: *Pseudomonadota* (*Gammaproteobacteria*); C: *Pseudomonadota* (*Alphaproteobacteria*); D: *Pseudomonadota* (*Betaproteobacteria*), *Campylobacterota; Bacteroidota*; E: *Actinomycetota*; F: *Bacillota*. Numbers at nodes: bootstrap values of 70% and larger. Scale bars: 0.1 substitutions per nucleotide position. Bold: sequences of strains cultivated from surface washed cilantro leaves (green coloured) and roots (brown colours) in this study or by irrigation water types (blue coloured) as described by Soufi *et al.* (2025).

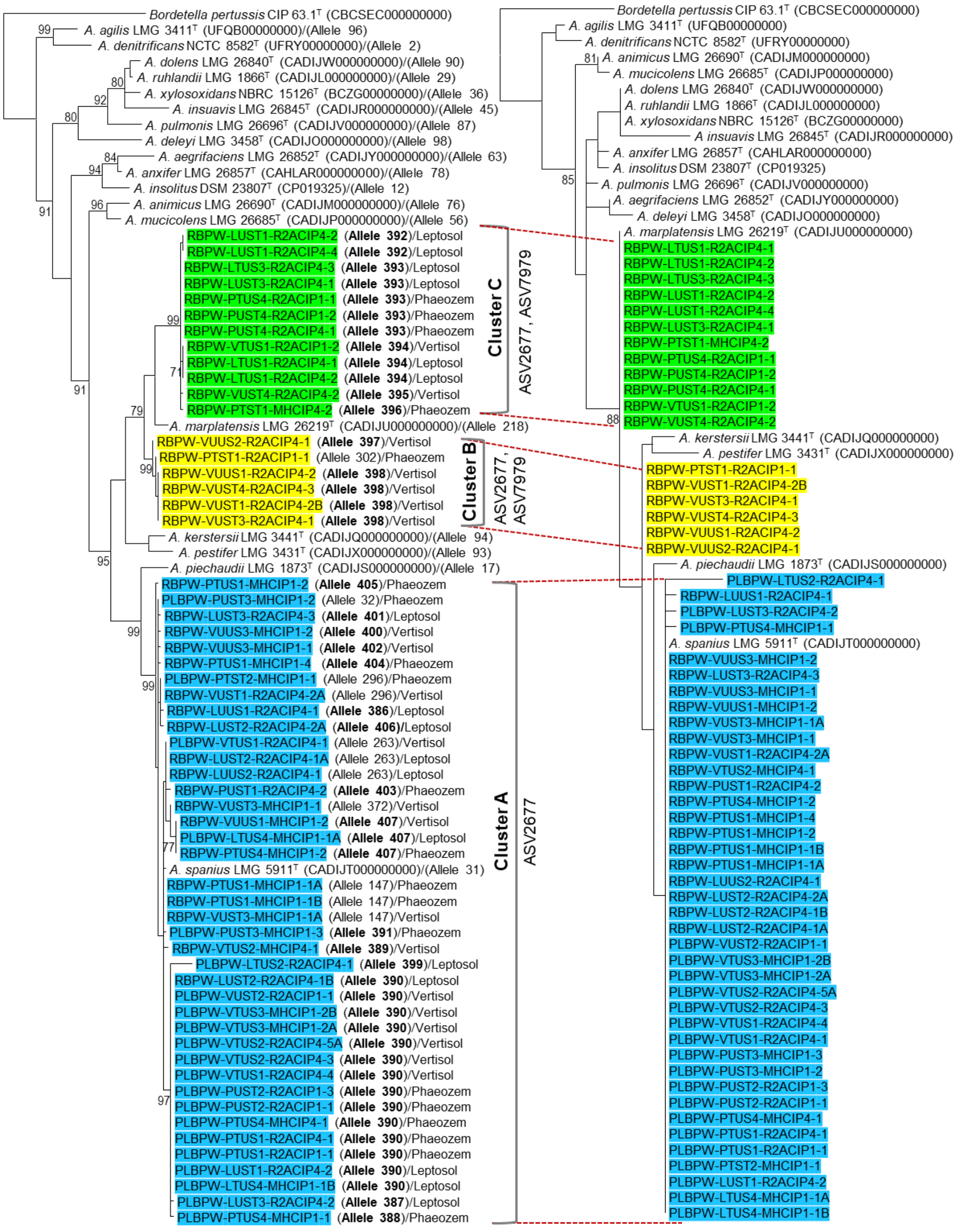

**(A) *nrdA*** **(B) NrdA**

**Figure S5.** Phylogenetic placement of *Achromobacter* spp. strains within the genus *Achromobacter*. Phylogenetic analyses were performed using Maximum Likelihood method (Felsenstein 1981). Nucleotide based analysis was carried out by the General Time Reversible model (GTR; Nei and Kumar 2000). Amino acid-based analysis was done by the JTT matrix-based model (Jones, Taylor and Thornton. 1992). A discrete Gamma distribution was used to model evolutionary rate differences among sites [5 categories (+G)] assuming some sites to be evolutionarily invariable (+I). The trees of (A) *nrdA* and (B) NrdA were based on total of 765 nucleotides and 255 amino acids positions in the final datasets. All positions containing gaps and missing data were eliminated. Bootstrap values (≥70%) after 100 resamplings are indicated at branch nodes; bar, number of substitutions per site. Evolutionary analyses were conducted in MEGA11 (Tamura *et al*., 2021). Novel alleles from this study were given in bold font.

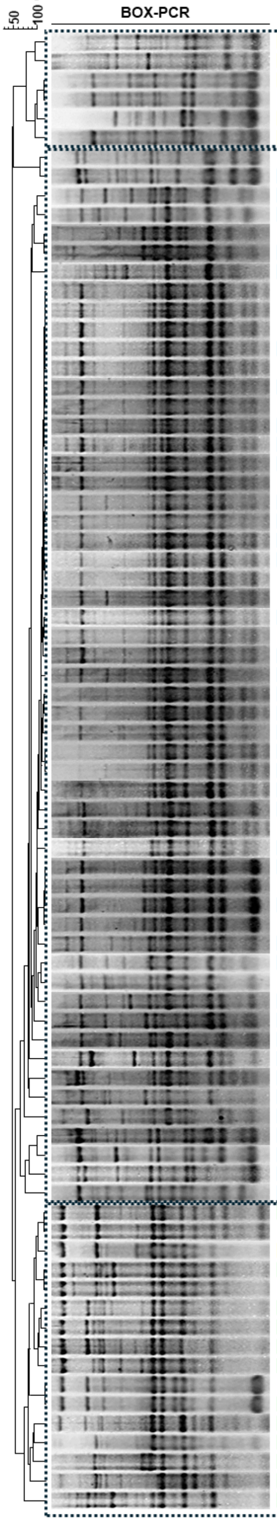

Legend: Unspiked-Influent (grey circle); Spiked-Influent (hatched grey circle); Unspiked-Effluent (cyan circle); Spiked-Effluent (hatched cyan circle); Leaves (green star); Roots (orange star).

QACs (μg mL^-1^): 1.BAC-C12, 2.DADMAC-C10, 3.ATMAC-C16

| Strains | Source | Plant part | Soil types | Genotype | ASV (16S rRNA gene similarity) | *nrdA* (765nt) allele | *nrdA* (765 nt) based spp. assignment | QACs 1 | QACs 2 | QACs 3 | FQs resistance | Cluster |
|---|---|---|---|---|---|---|---|---|---|---|---|---|
| **RBPW-VUST1-R2ACIP4-2B** | Spiked-Influent | Roots | Vertisol | 1a | ASV2677 (100%) | **398** | *Achromobacter* spp. | 50 | 10 | 100 | Susceptible | Cluster B |
| **RBPW-VUST3-R2ACIP4-1** | Spiked-Influent | Roots | Vertisol | 1b | ASV7979 (100%) | **398** | *Achromobacter* spp. | 50 | 10 | 50 | Susceptible | Cluster B |
| **RBPW-VUUS2-R2ACIP4-1** | Unspiked-Influent | Roots | Vertisol | 1c | ASV7979 (100%) | **397** | *Achromobacter* spp. | 25 | 10 | 50 | Susceptible | Cluster B |
| RBPW-VUUS2-R2ACIP4-2 | Spiked-Influent | Roots | Vertisol | 1c | n.d. | n.d. | n.d. | n.d. | n.d. | n.d. | n.d. | Cluster B |
| **RBPW-VUUS1-R2ACIP4-2** | Unspiked-Influent | Roots | Vertisol | 1d | ASV7979 (100%) | **398** | *Achromobacter* spp. | 25 | 10 | 50 | Susceptible | Cluster B |
| **RBPW-PTST1-R2ACIP1-1** | Spiked-Effluent | Roots | Phaeozem | 1e | ASV7979 (100%) | 302 | *Achromobacter* spp. | 50 | 10 | 50 | Susceptible | Cluster B |
| **RBPW-VTUS2-MHCIP1-3** | Spiked-Effluent | Roots | Vertisol | 2 | ASV2677 (100%) | n.d. | n.d. | n.d. | n.d. | n.d. | n.d. | Cluster A |
| **RBPW-VTUS2-MHCIP4-1** | Unspiked-Effluent | Roots | Vertisol | 2 | ASV2677 (100%) | **389** | *Achromobacter spanius* | 50 | 10 | 50 | CIP, LEV | Cluster A |
| RBPW-VUST1-R2ACIP4-2A | Spiked-Influent | Roots | Vertisol | 3a | ASV2677 (100%) | 296 | *Achromobacter spanius* | 25 | 10 | 50 | Susceptible | Cluster A |
| **RBPW-VUUS3-MHCIP1-1** | Unspiked-Influent | Roots | Vertisol | 3b | ASV2677 (100%) | **402** | *Achromobacter spanius* | 50 | 10 | 50 | Susceptible | Cluster A |
| **PLBPW-PUST3-MHCIP1-1** | Spiked-Influent | Leaves | Phaeozem | 3c | ASV2677 (100%) | n.d. | n.d. | n.d. | n.d. | n.d. | n.d. | Cluster A |
| PLBPW-PUST3-MHCIP1-3 | Spiked-Influent | Leaves | Phaeozem | 3c | n.d. | **391** | *Achromobacter spanius* | 50 | 10 | 50 | Susceptible | Cluster A |
| **PLBPW-VUST2-R2ACIP1-1** | Spiked-Influent | Leaves | Vertisol | 4 | ASV2677 (100%) | **390** | *Achromobacter spanius* | 25 | 10 | 50 | CIP | Cluster A |
| PLBPW-PTUS4-R2ACIP4-1 | Unspiked-Effluent | Leaves | Phaeozem | 5a-1 | n.d. | n.d. | n.d. | 25 | 10 | 50 | CIP, LEV | Cluster A |
| PLBPW-PTUS4-R2ACIP1-1 | Unspiked-Effluent | Leaves | Phaeozem | 5a-1 | n.d. | n.d. | n.d. | n.d. | n.d. | n.d. | n.d. | Cluster A |
| PLBPW-LTUS2-R2ACIP4-3 | Unspiked-Effluent | Leaves | Leptosol | 5a-1 | n.d. | n.d. | n.d. | n.d. | n.d. | n.d. | n.d. | Cluster A |
| **PLBPW-LTUS2-R2ACIP4-1** | Unspiked-Effluent | Leaves | Leptosol | 5a-1 | ASV2677 (100%) | **399** | *Achromobacter spanius* | 25 | 10 | 50 | CIP, LEV | Cluster A |
| PLBPW-LTUS4-R2ACIP4-1 | Unspiked-Effluent | Leaves | Leptosol | 5a-1 | n.d. | n.d. | n.d. | n.d. | n.d. | n.d. | n.d. | Cluster A |
| PLBPW-VTUS3-MHCIP1-2B | Unspiked-Effluent | Leaves | Vertisol | 5a-1 | n.d. | **390** | *Achromobacter spanius* | 50 | 10 | 50 | CIP, LEV | Cluster A |
| PLBPW-VTUS3-MHCIP1-4 | Unspiked-Effluent | Leaves | Vertisol | 5a-1 | n.d. | n.d. | n.d. | n.d. | n.d. | n.d. | n.d. | Cluster A |
| **RBPW-LUST2-R2ACIP4-1B** | Spiked-Influent | Roots | Leptosol | 5a-1 | ASV2677 (100%) | **390** | *Achromobacter spanius* | 25 | 5 | 50 | CIP, LEV | Cluster A |
| PLBPW-PTUS1-R2ACIP1-1 | Unspiked-Effluent | Leaves | Phaeozem | 5a-1 | n.d. | **390** | *Achromobacter spanius* | 50 | 10 | 100 | Susceptible | Cluster A |
| PLBPW-PTUS4-MHCIP1-1 | Unspiked-Effluent | Leaves | Phaeozem | 5a-1 | n.d. | **388** | *Achromobacter spanius* | 25 | 10 | 50 | Susceptible | Cluster A |
| PLBPW-PTUS4-MHCIP1-2 | Unspiked-Effluent | Leaves | Phaeozem | 5a-1 | n.d. | n.d. | n.d. | n.d. | n.d. | n.d. | n.d. | Cluster A |
| PLBPW-VTUS2-R2ACIP4-5A | Unspiked-Effluent | Leaves | Vertisol | 5a-1 | n.d. | **390** | *Achromobacter spanius* | 25 | 5 | 50 | CIP, LEV | Cluster A |
| PLBPW-VTUS1-R2ACIP4-4 | Unspiked-Effluent | Leaves | Vertisol | 5a-1 | n.d. | **390** | *Achromobacter spanius* | 25 | 10 | 50 | CIP, LEV | Cluster A |
| PLBPW-PUST2-R2ACIP1-3 | Spiked-Influent | Leaves | Phaeozem | 5a-1 | n.d. | **390** | *Achromobacter spanius* | 25 | 10 | 100 | Susceptible | Cluster A |
| **PLBPW-PTUS1-R2ACIP4-1** | Unspiked-Effluent | Leaves | Phaeozem | 5a-1 | ASV2677 (100%) | **390** | *Achromobacter spanius* | 50 | 10 | 50 | CIP, LEV | Cluster A |
| PLBPW-PUST2-R2ACIP1-2 | Spiked-Influent | Leaves | Phaeozem | 5a-1 | n.d. | n.d. | n.d. | n.d. | n.d. | n.d. | n.d. | Cluster A |
| PLBPW-LTUS2-R2ACIP4-4 | Unspiked-Effluent | Leaves | Leptosol | 5a-1 | n.d. | n.d. | n.d. | n.d. | n.d. | n.d. | n.d. | Cluster A |
| PLBPW-LUST1-R2ACIP4-2 | Spiked-Influent | Leaves | Leptosol | 5a-1 | n.d. | **390** | *Achromobacter spanius* | 50 | 10 | 50 | CIP, LEV | Cluster A |
| PLBPW-VTUS1-R2ACIP4-5 | Unspiked-Effluent | Leaves | Vertisol | 5a-1 | n.d. | n.d. | n.d. | n.d. | n.d. | n.d. | n.d. | Cluster A |
| **PLBPW-LUST3-R2ACIP4-2** | Spiked-Influent | Leaves | Leptosol | 5a-1 | ASV2677 (100%) | **387** | *Achromobacter spanius* | 50 | 10 | 50 | CIP | Cluster A |
| **PLBPW-PTST2-MHCIP1-1** | Spiked-Effluent | Leaves | Phaeozem | 5a-1 | ASV2677 (100%) | 296 | *Achromobacter spanius* | 50 | 10 | 50 | CIP, LEV | Cluster A |
| PLBPW-LTUS4-MHCIP1-1B | Unspiked-Effluent | Leaves | Leptosol | 5a-2 | n.d. | **390** | *Achromobacter spanius* | 50 | 10 | 50 | CIP, LEV | Cluster A |
| **PLBPW-PTUS4-MHCIP4-1** | Unspiked-Effluent | Leaves | Phaeozem | 5a-2 | ASV2677 (100%) | **390** | *Achromobacter spanius* | 50 | 10 | 50 | CIP, LEV | Cluster A |
| **RBPW-PTUS1-MHCIP1-4** | Unspiked-Effluent | Roots | Phaeozem | 5a-2 | ASV2677 (100%) | **404** | *Achromobacter spanius* | 50 | 10 | 50 | Susceptible | Cluster A |
| **PLBPW-VTUS3-MHCIP1-2A** | Unspiked-Effluent | Leaves | Vertisol | 5a-3 | ASV2677 (100%) | **390** | *Achromobacter spanius* | 50 | 10 | 50 | Susceptible | Cluster A |
| PLBPW-VTUS2-R2ACIP4-3 | Unspiked-Effluent | Leaves | Vertisol | 5a-3 | n.d. | **390** | *Achromobacter spanius* | 25 | 10 | 50 | CIP, LEV | Cluster A |
| PLBPW-PUST2-R2ACIP1-1 | Spiked-Influent | Leaves | Phaeozem | 5a-3 | n.d. | **390** | *Achromobacter spanius* | 50 | 10 | 50 | Susceptible | Cluster A |
| **RBPW-LUST2-R2ACIP4-2A** | Spiked-Influent | Roots | Leptosol | 5a-4 | ASV2677 (100%) | **406** | *Achromobacter spanius* | 50 | 10 | 50 | CIP, LEV | Cluster A |
| PLBPW-VTUS1-R2ACIP4-3 | Unspiked-Effluent | Leaves | Vertisol | 5a-5 | n.d. | n.d. | n.d. | 50 | 10 | 50 | CIP, LEV | Cluster A |
| **RBPW-PTUS1-MHCIP1-1B** | Unspiked-Effluent | Roots | Phaeozem | 5b | ASV2677 (100%) | 147 | *Achromobacter spanius* | 50 | 10 | 50 | Susceptible | Cluster A |
| RBPW-LUUS2-R2ACIP4-1 | Unspiked-Influent | Roots | Leptosol | 6a | ASV2677 (100%) | 263 | *Achromobacter spanius* | 25 | 10 | 50 | Susceptible | Cluster A |
| **RBPW-LUST2-R2ACIP1-3** | Spiked-Influent | Roots | Leptosol | 6a | ASV2677 (100%) | n.d. | n.d. | n.d. | n.d. | n.d. | n.d. | Cluster A |
| **RBPW-LUST2-R2ACIP4-1A** | Spiked-Influent | Roots | Leptosol | 6a | ASV2677 (100%) | 263 | *Achromobacter spanius* | 25 | 10 | 50 | CIP | Cluster A |
| RBPW-LUST2-R2ACIP1-4 | Spiked-Influent | Roots | Leptosol | 6a | n.d. | n.d. | n.d. | n.d. | n.d. | n.d. | n.d. | Cluster A |
| **RBPW-LUST3-R2ACIP4-3** | Spiked-Influent | Roots | Leptosol | 6b | ASV2677 (100%) | **401** | *Achromobacter spanius* | 25 | 10 | 50 | CIP, LEV | Cluster A |
| RBPW-PTUS1-MHCIP1-1A | Unspiked-Effluent | Roots | Phaeozem | 7 | n.d. | 147 | *Achromobacter spanius* | 50 | 10 | 50 | Susceptible | Cluster A |
| **RBPW-PTUS4-MHCIP1-1B** | Unspiked-Effluent | Roots | Phaeozem | 7 | ASV2677 (100%) | n.d. | n.d. | n.d. | n.d. | n.d. | n.d. | Cluster A |
| RBPW-PTUS4-MHCIP1-2 | Unspiked-Effluent | Roots | Phaeozem | 8 | ASV2677 (100%) | **407** | *Achromobacter spanius* | 50 | 10 | 50 | Susceptible | Cluster A |
| **RBPW-PUST1-R2ACIP4-2** | Spiked-Influent | Roots | Phaeozem | 9 | ASV2677 (100%) | **403** | *Achromobacter spanius* | 25 | 10 | 50 | CIP, LEV | Cluster A |
| **PLBPW-VTUS1-R2ACIP4-1** | Unspiked-Effluent | Leaves | Vertisol | 10 | ASV2677 (100%) | 263 | *Achromobacter spanius* | 50 | 10 | 50 | CIP, LEV | Cluster A |
| **PLBPW-PUST3-MHCIP1-2** | Spiked-Influent | Leaves | Phaeozem | 11 | ASV2677 (100%) | 32 | *Achromobacter spanius* | 50 | 10 | 100 | Susceptible | Cluster A |
| **RBPW-LUUS1-R2ACIP4-1** | Unspiked-Influent | Roots | Leptosol | 12 | ASV2677 (100%) | **386** | *Achromobacter spanius* | 25 | 10 | 50 | CIP, LEV | Cluster A |
| RBPW-VUUS3-MHCIP1-2 | Unspiked-Influent | Roots | Vertisol | 13 | ASV2677 (100%) | **400** | *Achromobacter spanius* | 50 | 10 | 50 | Susceptible | Cluster A |
| **RBPW-PTUS1-MHCIP1-2** | Unspiked-Effluent | Roots | Phaeozem | 14 | ASV2677 (100%) | **405** | *Achromobacter spanius* | 50 | 10 | 50 | Susceptible | Cluster A |
| **PLBPW-LTUS4-MHCIP1-1A** | Unspiked-Effluent | Leaves | Leptosol | 15a | ASV2677 (100%) | **407** | *Achromobacter spanius* | 50 | 10 | 50 | Susceptible | Cluster A |
| **RBPW-VUUS1-MHCIP1-2** | Unspiked-Influent | Roots | Vertisol | 15b | ASV2677 (100%) | **407** | *Achromobacter spanius* | 25 | 5 | 50 | Susceptible | Cluster A |
| **RBPW-VUST3-MHCIP1-1A** | Spiked-Influent | Roots | Vertisol | 16 | ASV2677 (100%) | 147 | *Achromobacter spanius* | 50 | 10 | 50 | CIP, LEV | Cluster A |
| RBPW-VUST3-MHCIP1-1 | Spiked-Influent | Roots | Vertisol | 17 | ASV2677 (100%) | **372** | *Achromobacter spanius* | 50 | 10 | 50 | CIP, LEV | Cluster A |
| **RBPW-PTST1-MHCIP4-2** | Spiked-Effluent | Roots | Phaeozem | 18 | ASV7979 (100%) | **396** | *Achromobacter* spp. | 50 | 10 | 50 | CIP, LEV | Cluster C |
| **RBPW-PTST1-MHCIP4-1** | Spiked-Effluent | Roots | Phaeozem | 18 | ASV7979 (100%) | n.d. | n.d. | n.d. | n.d. | n.d. | n.d. | Cluster C |
| RBPW-VUST4-R2ACIP4-3 | Spiked-Influent | Roots | Vertisol | 19 | n.d. | **398** | *Achromobacter* spp. | 50 | 10 | 50 | Susceptible | Cluster C |
| **RBPW-LUST1-R2ACIP4-2** | Spiked-Influent | Roots | Leptosol | 20 | ASV7979 (100%) | **392** | *Achromobacter* spp. | 50 | 10 | 50 | Susceptible | Cluster C |
| RBPW-LUST1-R2ACIP4-3 | Spiked-Influent | Roots | Leptosol | 20 | n.d. | n.d. | n.d. | n.d. | n.d. | n.d. | n.d. | Cluster C |
| **RBPW-LTUS1-R2ACIP4-2** | Unspiked-Effluent | Roots | Leptosol | 21a | ASV7979 (100%) | **394** | *Achromobacter* spp. | 25 | 10 | 50 | Susceptible | Cluster C |
| **RBPW-PUST4-R2ACIP1-2** | Spiked-Influent | Roots | Phaeozem | 21a | ASV7979 (100%) | **393** | *Achromobacter* spp. | 50 | 10 | 50 | Susceptible | Cluster C |
| **RBPW-VTUS1-R2ACIP1-2** | Unspiked-Effluent | Roots | Vertisol | 21b | ASV2677 (100%) | **394** | *Achromobacter* spp. | 25 | 10 | 50 | CIP, LEV | Cluster C |
| **RBPW-LTUS3-R2ACIP4-3** | Unspiked-Effluent | Roots | Leptosol | 21c | ASV7979 (100%) | **393** | *Achromobacter* spp. | 12.5 | 2.5 | 6.25 | Susceptible | Cluster C |
| RBPW-PTUS4-R2ACIP1-3 | Unspiked-Effluent | Roots | Phaeozem | 22 | n.d. | n.d. | n.d. | n.d. | n.d. | n.d. | n.d. | Cluster C |
| **RBPW-PTUS4-R2ACIP1-1** | Unspiked-Effluent | Roots | Phaeozem | 22 | ASV7979 (100%) | **393** | *Achromobacter* spp. | 50 | 10 | 50 | Susceptible | Cluster C |
| **RBPW-PUST4-R2ACIP4-1** | Spiked-Influent | Roots | Phaeozem | 23 | ASV7979 (100%) | **393** | *Achromobacter* spp. | 50 | 10 | 50 | Susceptible | Cluster C |
| **RBPW-VUST4-R2ACIP4-2** | Spiked-Influent | Roots | Vertisol | 24 | ASV7979 (100%) | **395** | *Achromobacter* spp. | 50 | 10 | 50 | Susceptible | Cluster C |
| **RBPW-LUST3-R2ACIP4-1** | Spiked-Influent | Roots | Leptosol | 25 | ASV7979 (100%) | **393** | *Achromobacter* spp. | 25 | 10 | 50 | Susceptible | Cluster C |
| **RBPW-LUST1-R2ACIP4-4** | Spiked-Influent | Roots | Leptosol | 26 | ASV7979 (100%) | **392** | *Achromobacter* spp. | 50 | 10 | 50 | Susceptible | Cluster C |
| **RBPW-LTUS1-R2ACIP4-1** | Unspiked-Effluent | Roots | Leptosol | 27 | ASV7979 (100%) | **394** | *Achromobacter* spp. | 50 | 10 | 50 | Susceptible | Cluster C |

*Achromobacter* **Phylotype A-1** (99–100%, pairwise 16S rRNA gene similarities to ***A. spanius***, ***A. marplatensis***, ***A. deleyi***, ***A. kerstersii*** and ***A. piechaudii***)

**Figure S6.** Genotyping of all *Achromobacter* spp. strains based on fingerprint patterns obtained by genomic fingerprinting using primers targeting BOX repetitive elements. Cluster analysis was performed in BioNumerics version 8 (Applied Maths, Belgium) using UPGMA clustering, based on a dissimilarity matrix generated by the Pearson correlation coefficient. Strains in bold font were identified by partial 16S rRNA gene sequencing. Species assignment as *Achromobacter spanius* was based on *nrdA* gene, and species assignment provided inside the brackets were based on identical BOX pattern (same genotype). Allele numbers with bold font represented novel alleles from this study.

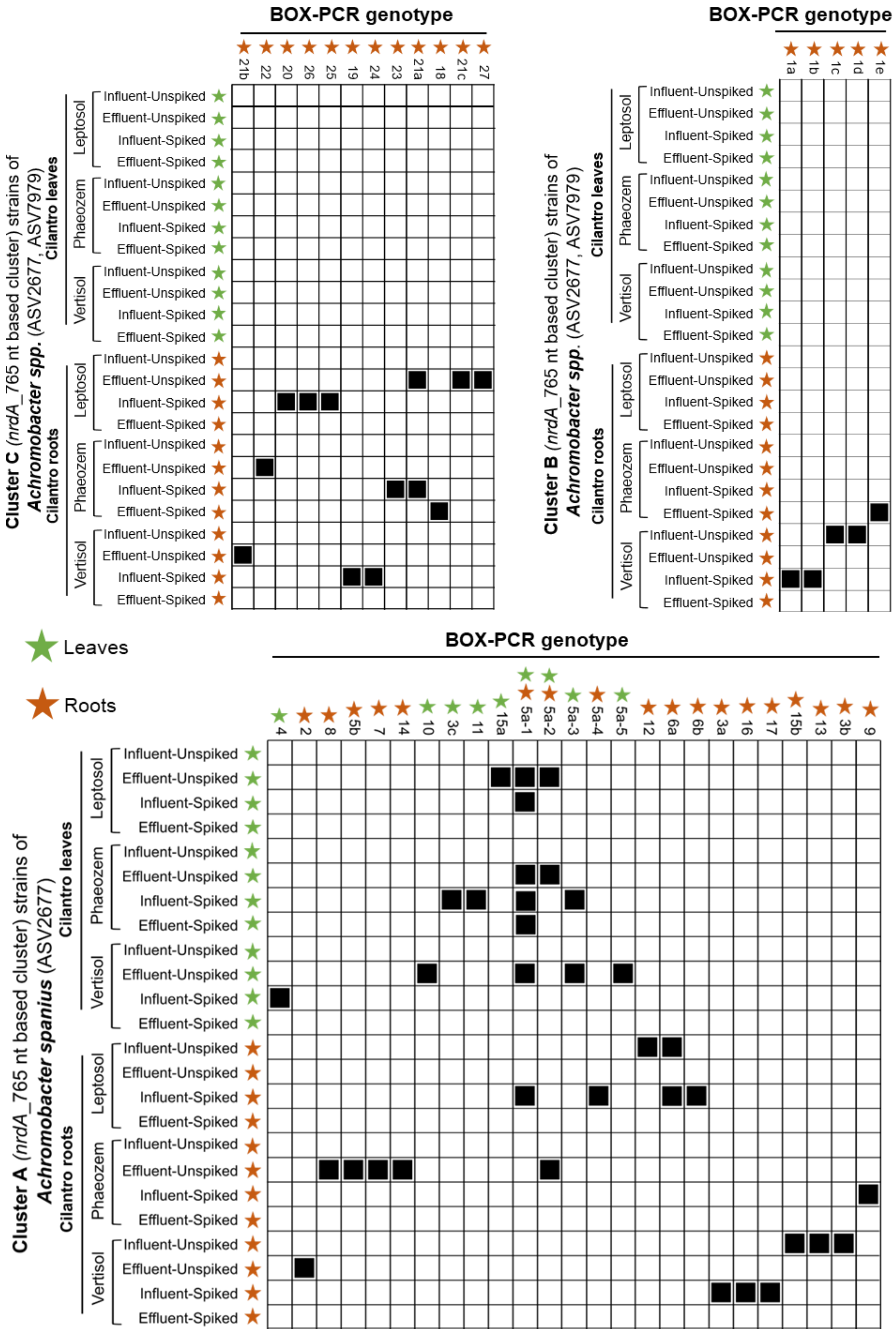

**Figure S7.** Seriation analysis based on an absence-presence (0/1) matrix illustrating the presence of *Achromobacter* genotypes in the leaves and roots of cilantro grown on different soil types irrigated with different wastewater types.

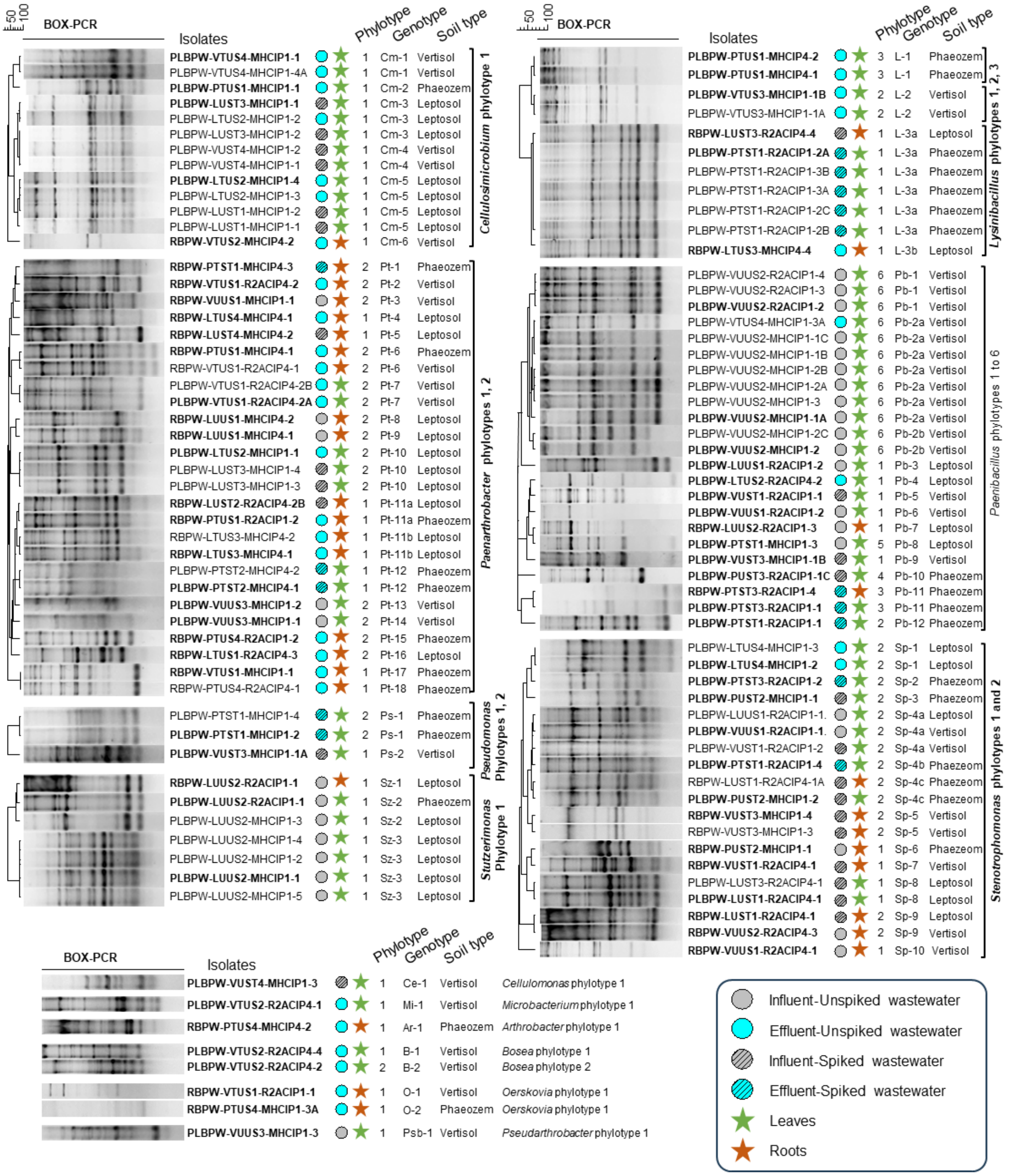

**Figure S8.** Genotyping of all bacterial phylotypes beside *Achromobacter* based on fingerprint patterns obtained by genomic fingerprinting using primer BOXA1R. Cluster analysis was performed in BioNumerics version 8 (Applied Maths, Belgium) using UPGMA clustering, based on a dissimilarity matrix generated by the Pearson correlation coefficient. Strains in bold font were identified by partial 16S rRNA gene sequencing.

**Supplementary reference**

1. L. Soufi, I. D. Kampouris, K. Lüneberg, B. J. Heyde, D. Pulami, S. P. Glaeser, C. Siebe, J. Siemens, K. Smalla, E. Grohmann and S. Gallego, Wastewater-borne pollutants influenced antibiotic resistance genes and mobile genetic elements in the soil without affecting the bacterial community composition in a changing wastewater irrigation system. *J Hazard Mater*. 2025, **494**, 138680.
2. J. Siemens, G. Huschek, C. Siebe, M Kaupenjohann. Concentrations and mobility of human pharmaceuticals in the world's largest wastewater irrigation system, Mexico City-Mezquital Valley. *Water Res.,* 2008, ***42***, 2124–2134.
3. J. Versalovic, M. Schneider, F. J. de Bruijn and J. R. Lupski, Genomic fingerprinting of bacteria using repetitive sequence-based polymerase chain reaction, *Methods Mol. Cell Biol*., 1994, ***5***, 25-40.
4. S. P. Glaeser, H. Galatis, K. Martin and P. Kämpfer, *Niabella hirudinis* and *Niabella drilacis* sp. nov., isolated from the medicinal leech Hirudo verbana, *Int. J. Syst. Evol. Microbiol*., 2013, ***63***, 3487-3493.
5. D. J. Lane, 16S/23S rRNA Sequencing, *Nucleic acid techniques in bacterial systematics*, 1991.
6. T. Spilker, P. Vandamme and J. J. LiPuma, A multilocus sequence typing scheme implies population structure and reveals several putative novel *Achromobacter* species, *J. Clin. Microbiol*., 2012, **50**(9), 3010-3015.
7. A. Magallon, M. Roussel, C. Neuwirth, J. Tetu, A. C. Cheiakh, B. Boulet, V. Varin, V. Urbain, J. Bador and L. Amoureux, Fluoroquinolone resistance in *Achromobacter* spp.: substitutions in QRDRs of GyrA, GyrB, ParC and ParE and implication of the RND efflux system AxyEF-OprN. J. *Antimicrob. Chemother.*, 2021, **76**, 297–304.
8. J. P. R. Furlan, D. G. Sanchez, I. F. L. Gallo and E. G. Stehling, Replicon typing of plasmids in environmental *Achromobacter* sp. producing quinolone-resistant determinants, *APMIS*, 2018, **126**(11), 864-869.
9. V. Cattoir, L. Poirel, V. Rotimi, C. J. Soussy and P. Nordmann, Multiplex PCR for detection of plasmid-mediated quinolone resistance *qnr* genes in ESBL-producing enterobacterial isolates, *J. Antimicrob. Chemother.*, 2007, **60**(2), 394-397.
10. S. Alipour, M. Owrang, M. Rajabnia, M. Olfatifar, H. Kazemian and H. Houri, Prevalence of plasmid-mediated quinolone resistance genes in *Escherichia coli* isolates from colonic biopsies of Iranian patients with inflammatory bowel diseases: a cross-sectional study. *Health Sci Rep*. 2024, **7**(12), e70204.
11. J. Felsenstein, Evolutionary trees from DNA sequences: a maximum likelihood approach, *J. Mol. Evol*., 1981, **17**(6), 368-376.
12. M. Nei and S. Kumar, *Molecular evolution and phylogenetics*. Oxford university press, 2000.
13. D. T. Jones, W. R. Taylor and J. M. Thornton, The rapid generation of mutation data matrices from protein sequences, *Bioinformatics*, 1992, **8**(3), 275-282.
14. K. Tamura, G. Stecher and S. Kumar, MEGA11: Molecular evolutionary genetics analysis version 11. *Mol. Biol. Evol.*, 2021, **38**(7), 3022-3027.
